# A single-cell and single-nucleus RNA-seq toolbox for fresh and frozen human tumors

**DOI:** 10.1101/761429

**Authors:** Michal Slyper, Caroline B. M. Porter, Orr Ashenberg, Julia Waldman, Eugene Drokhlyansky, Isaac Wakiro, Christopher Smillie, Gabriela Smith-Rosario, Jingyi Wu, Danielle Dionne, Sébastien Vigneau, Judit Jané-Valbuena, Sara Napolitano, Mei-Ju Su, Anand G. Patel, Asa Karlstrom, Simon Gritsch, Masashi Nomura, Avinash Waghray, Satyen H. Gohil, Alexander M. Tsankov, Livnat Jerby-Arnon, Ofir Cohen, Johanna Klughammer, Yanay Rosen, Joshua Gould, Bo Li, Lan Nguyen, Catherine J. Wu, Benjamin Izar, Rizwan Haq, F. Stephen Hodi, Charles H. Yoon, Aaron N. Hata, Suzanne J. Baker, Mario L. Suvà, Raphael Bueno, Elizabeth H. Stover, Ursula A. Matulonis, Michael R. Clay, Michael A. Dyer, Natalie B. Collins, Nikhil Wagle, Asaf Rotem, Bruce E. Johnson, Orit Rozenblatt-Rosen, Aviv Regev

## Abstract

Single cell genomics is essential to chart the complex tumor ecosystem. While single cell RNA-Seq (scRNA-Seq) profiles RNA from cells dissociated from fresh tumor tissues, single nucleus RNA-Seq (snRNA-Seq) is needed to profile frozen or hard-to-dissociate tumors. Each strategy requires modifications to fit the unique characteristics of different tissue and tumor types, posing a barrier to adoption. Here, we developed a systematic toolbox for profiling fresh and frozen clinical tumor samples using scRNA-Seq and snRNA-Seq, respectively. We tested eight tumor types of varying tissue and sample characteristics (resection, biopsy, ascites, and orthotopic patient-derived xenograft): lung cancer, metastatic breast cancer, ovarian cancer, melanoma, neuroblastoma, pediatric sarcoma, glioblastoma, pediatric high-grade glioma, and chronic lymphocytic leukemia. Analyzing 212,498 cells and nuclei from 39 clinical samples, we evaluated protocols by cell quality, recovery rate, and cellular composition. We optimized protocols for fresh tissue dissociation for different tumor types using a decision tree to account for the technical and biological variation between clinical samples. We established methods for nucleus isolation from OCT embedded and fresh-frozen tissues, with an optimization matrix varying mechanical force, buffer, and detergent. scRNA-Seq and snRNA-Seq from matched samples recovered the same cell types and intrinsic expression profiles, but at different proportions. Our work provides direct guidance across a broad range of tumors, including criteria for testing and selecting methods from the toolbox for other tumors, thus paving the way for charting tumor atlases.

## Introduction

Single cell RNA-Seq (scRNA-Seq) has transformed our ability to analyze tumors, revealing cell types, states, genetic diversity, and interactions in the complex tumor ecosystem(Cieslik and Chinnaiyan, 2018; Filbin et al., 2018; Jerby-Arnon et al., 2018; Puram et al., 2017; Tirosh et al., 2016; Venteicher et al., 2017). However, successful scRNA-Seq requires dissociation tailored to the tumor type, and involves enzymatic digestion that can lead to loss of sensitive cells or changes in gene expression. Moreover, obtaining fresh tissue is time-sensitive and requires tight coordination between tissue acquisition and processing teams, posing a challenge in clinical settings. Conversely, single-nucleus RNA-Seq (snRNA-Seq) allows profiling of single nuclei isolated from frozen tissues, decoupling tissue acquisition from immediate sample processing. snRNA-Seq can also handle samples that cannot be successfully dissociated even when fresh, due to size or cell fragility(Habib et al., 2017; Habib et al., 2016), as well as multiplexed analysis of longitudinal samples from the same individual(Gaublomme et al., 2019). However, nuclei have lower amounts of mRNA compared to cells, and are more challenging to enrich or deplete. Both scRNA-Seq and snRNA-Seq pose experimental challenges when applied to different tumor types, due to distinct cellular composition and extracellular matrix (ECM) in different tumors.

To address these challenges, we developed a systematic toolbox for fresh and frozen tumor processing using single cell (sc) and single nucleus (sn) RNA-Seq, respectively (**Fig. 1A**). We tested eight tumor types with different tissue characteristics (**Fig. 1B**), including comparisons of matched fresh and frozen preparations from the same tumor specimen. The tumor types span different cell-of-origin (*e.g.*, epithelial, neuronal), solid and non-solid, patient ages, and transitions (*e.g.*, primary, metastatic, **Fig. 1B**).

**Figure 1.**
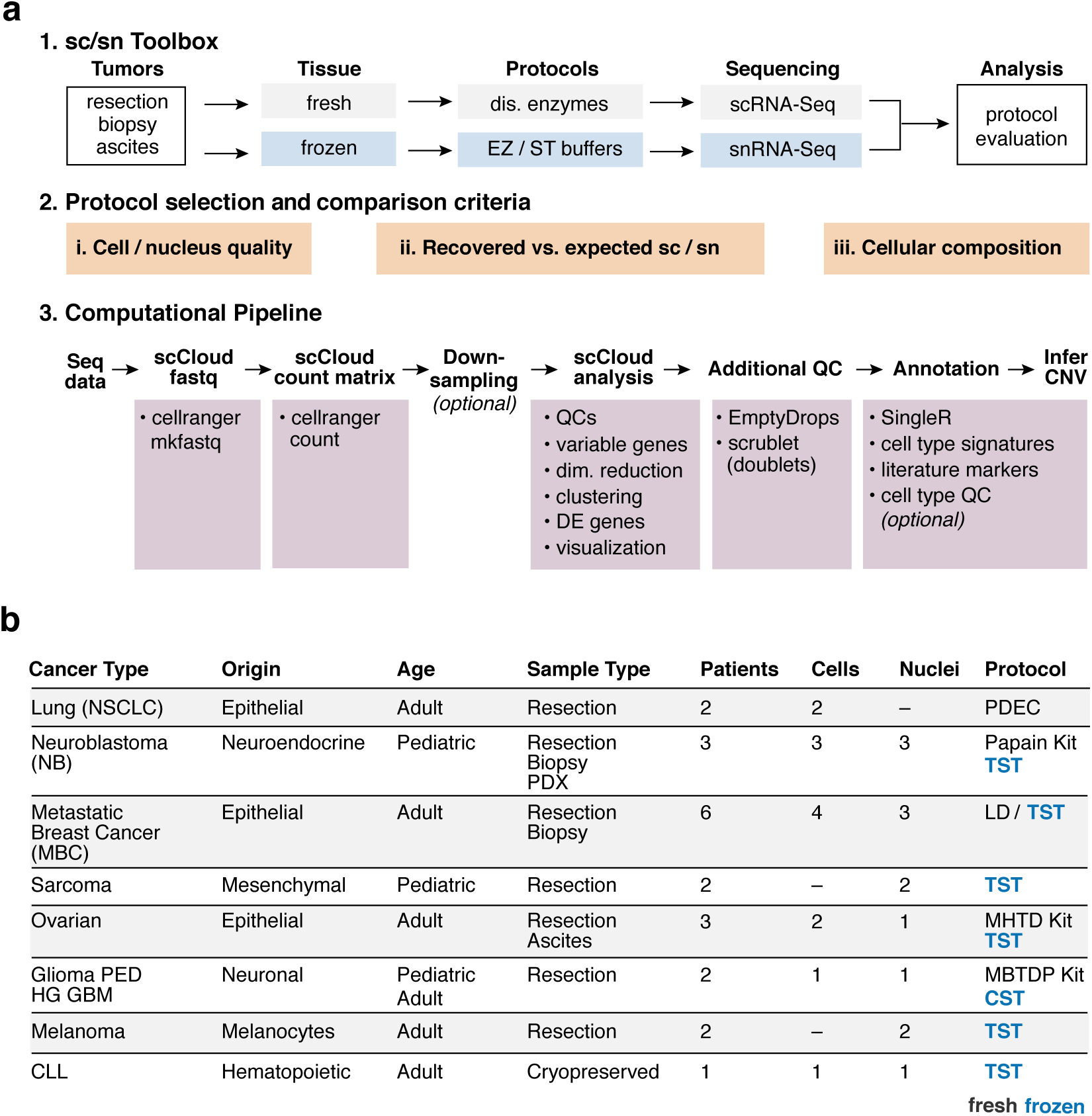
Overview of single-cell genomics toolboxes and tumor types profiled. **(A,B)** Study Overview. (**A**) sc/snRNA-Seq workflow, experimental and computational pipelines, and protocol selection criteria. **(B)** Tumor types in the study. Right column: recommended protocols for fresh (black/cells) or frozen (blue/nuclei) tumor samples.

## Results

We evaluated and compared protocols based on (i) cell/nucleus quality; (ii) number of recovered *vs*. expected cells/nuclei; and (iii) cellular composition (**Fig. 1A**). For “cell/nucleus quality”, we considered both experimental and computational metrics. Experimentally, we measured cell viability (for scRNA-Seq), the extent of doublets or aggregates in the cell/nucleus suspension, and cDNA quality recovered after Whole Transcriptome Amplification (**Methods**). Computationally, we evaluated the overall number of sequencing reads in a library, the percent of reads mapping to the transcriptome, genome, and intergenic regions, the number of cells/nuclei exceeding a minimal number of genes and unique transcripts (reflected by Unique Molecular Identifiers; UMI), the number of reads, transcripts (UMIs), and genes detected per cell/nucleus, and the percent of UMIs from mitochondrial genes (**Methods**). For “number of recovered *vs*. expected cells/nuclei”, we considered the proportion of droplets scored as likely empty (*i.e.*, containing only ambient RNA rather than the RNA from an encapsulated cell(Lun et al., 2019)), and the proportion of doublets(Wolock et al., 2019) (**Methods**). Finally, for “cellular composition”, we considered the diversity of cell types captured, the proportion of cells/nuclei recovered from each subset, and the copy number aberration (CNA) pattern classes that are recovered in malignant cells (**Methods**). We annotated the malignant cells based on the presence of CNAs (when detectable) and the cell type signature they most closely resembled (**Methods**). We conducted most data analysis using scCloud, a Cloud based single-cell analysis pipeline (Li et al., 2019) (BL, JG, YR, ORR and AR, **Methods**, **Fig. 1A**).

For scRNA-seq, our toolbox encompasses successful protocols for five types of fresh tumors: non-small cell lung carcinoma (NSCLC), metastatic breast cancer (MBC), ovarian cancer, glioblastoma (GBM), and neuroblastoma, as well as a cryopreserved non-solid, chronic lymphocytic leukemia (CLL) (**Fig. 1B**, **Supp. Fig. 1**). We constructed workflows that minimize the time interval between removal of the sample from the patient in a clinical setting and its dissociation into cells, to maximize cell viability and preservation of RNA profiles. We determined dissociation conditions for each of the tumor types and constructed specific steps as a decision tree to adjust for differences between types of clinical samples (*e.g.*, size, presence of red blood cells) (**Fig. 2A**, **Methods**). To choose the best performing dissociation method, we apportioned large tumor specimens into smaller pieces (∼0.5-2 cm), dissociating each piece with a different protocol.

**Figure 2.**
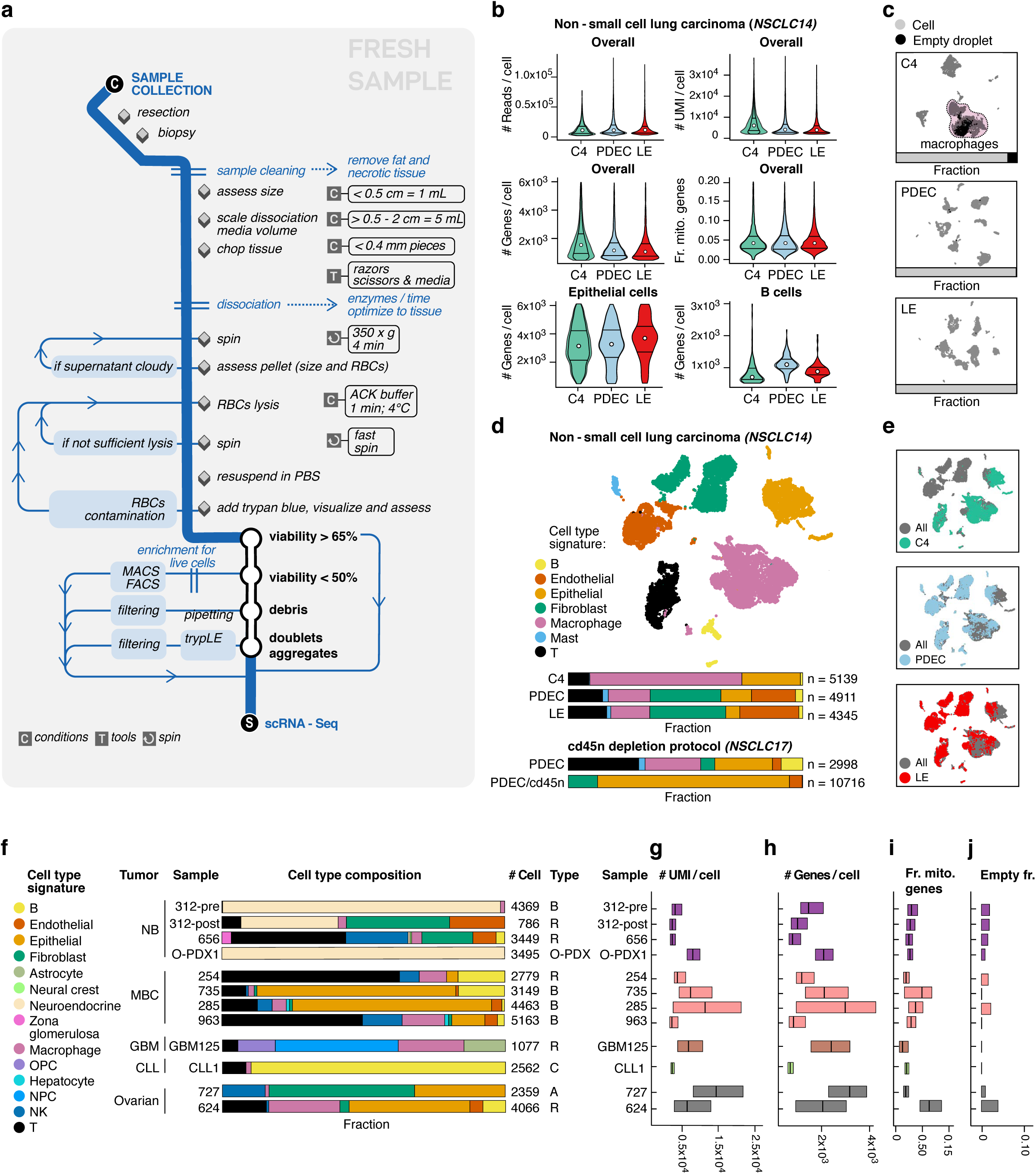
scRNA-Seq toolbox for fresh tumor samples. **(A)** Flow chart for collection and processing of fresh tumor samples. **(B-E)** Comparison of three dissociation protocols applied to one NSCLC sample. (**B**) Protocol performance varies across cell types. Top and middle: Distribution (median and first and third quartiles) of number of reads per cell, number of UMIs per cell, number of genes per cell, and fraction of UMIs per cell mapping to mitochondrial genes (*y* axes) in each protocol (*x* axis) across the entire dataset, Bottom: Distribution (median and first and third quartiles) of number of genes/cell (*y* axis) only in epithelial cells (left) or in B cells (right). Number of cells (*k*) in C4, PDEC, LE respectively is Overall: 5,139, 4,911, 4,345; Epithelial: 1,284, 641, 260; B cells: 100, 121, 78. **(C)** Protocols vary in number of empty drops. UMAP embedding of single cell profiles (dots) for each protocol, colored by assignment as cell (grey) or empty drop (black). Horizontal bars: fraction of assigned cells (grey) and empty drops (black). (**D,E)** Protocols vary in diversity of cell types captured. (**D**) Top: UMAP embedding of single cell profiles (dots) from all three protocols, colored by assigned cell subset signature. Bottom: Proportion of cells in each subset in each of the three protocols, and in an analysis using CD45 depletion; *n* indicates the number of recovered cells passing QC. **(E)** UMAP embedding as in (D) colored by protocol. **(F-J)** Protocol comparison across tumor types. (**F**) Cell type composition. Proportion of cells assigned to each cell subset signature (color) for each sample. R: Resection; B: Biopsy; A: Ascites; C: Cryopreserved; O-PDX: Orthotopic patient-derived xenograft. **(G-J)** QC metrics. The number of UMIs per cell, number of genes per cell, fraction of UMIs per cell mapping to mitochondrial genes, and fraction of empty drops (*x* axes) for each sample in (**F**) (*y* axis). The median of the distributions along with the first and third quartiles are shown in **(G-I**).

We selected enzymatic mixtures for processing fresh tissues based on the specific characteristics of each tumor type, such as cell type composition and ECM components, and ultimately recommend the method that sufficiently breaks down the ECM and cell-to-cell adhesions, while minimizing processing time and supporting the cell type diversity in the sample. For example, to break down collagen fibers in breast cancer(Al-Hajj et al., 2003; McDivitt et al., 1984), we used Liberase TM (**Methods**), whereas to break down ECM in GBM(Neftel et al., 2019) we used papain (cysteine protease). We also included DNase I to digest DNA released from dead cells to decrease viscosity in all dissociation mixtures. We subjected the samples that yielded high quality single cell suspensions to droplet-based scRNA-Seq (**Methods**).

As an example of the optimization process, consider NSCLC (sample NSCLC14, **Supp. Fig. 1-3**) where we used three processing protocols: (**1**) Collagenase 4 [NSCLC-C4]; (**2**) a mixture of Pronase, Dispase, Elastase, and Collagenases A and 4 [PDEC]; or (**3**) Liberase TM and Elastase [LE]; each in combination with DNase I (**Methods**) (**Fig. 2B-E, Supp. Fig. 2-3**). For the other tumor types, we show the application of our recommended protocol out of those tested (**Fig. 2F-J, Supp. Fig. 1**).

Protocols often performed similarly on standard quality control measures (*e.g.*, number of cells recovered), but differed markedly in cellular diversity or in the fraction of droplets predicted to contain only ambient RNA (“empty drops”) — two evaluation criteria that we prioritized. For example, in the NSCLC resection sample above, all methods yielded a similar number of cells with high-quality expression profiles and similar CNA patterns in malignant cells (**Fig. 2B-E, Supp. Fig. 2A-L**). However, only the PDEC and LE protocols recovered stromal and endothelial cells (**Fig. 2D, Supp. Fig. 2G**), and C4 had a 100-fold higher fraction of drops called as “empty” (7% *vs*. 0.08% and 0.04% in PDEC and LE, respectively, **Supp. Fig. 2A**). The drops designated “empty” in C4 clustered within macrophages (**Fig. 2C, Supp. Fig. 2E,G-I**), the most prevalent cell type, suggesting that these cell barcodes either had lower sequencing saturation or that the sample itself had higher ambient RNA content. While we estimated similarly low levels of ambient RNA(Young and Behjati, 2018) across the three protocols (**Supp. Fig. 2M-O**), NCSLC-C4 indeed had lower sequencing saturation and lower reads per cell (**Supp. Fig. 2A,C**). Ultimately, taking all of these features into consideration, we recommend the PDEC protocol for processing NSCLC tumor samples.

Comparing QC metrics across protocols can be challenging due to differences in cell type recovery and in sequencing depth between preparations, which we controlled for by also evaluating QC metrics *within* each cell type and down-sampling by total reads across protocols (**Supp. Fig. 2D and 3**). For example, overall, for the NSCLC sample, C4 had a significantly higher median number of detected genes (*P*=1.3*10^-90^ *vs*. PDEC; 1.4*10^-62^ *vs*. LE, two-sided Mann-Whitney U test), but within B cells, PDEC had a significantly higher number of detected genes (*P*=2*10^-15^ *vs*. C4; 2*10^-10^ *vs*. LE), whereas within epithelial cells, LE had the highest number (*P*=5*10^-6^ *vs*. C4; 2*10^-4^ *vs*. PDEC) (**Fig. 2B, Supp. Fig. 2D**). Because cell type proportions may vary between protocols, and the number of detected genes (and other metrics) varies between cell types, it is important also to assess cell-type specific QCs when choosing a protocol. Down-sampling by total reads did not qualitatively change any of our protocol evaluation metrics (**Supp. Fig. 3**).

**Figure 3.**
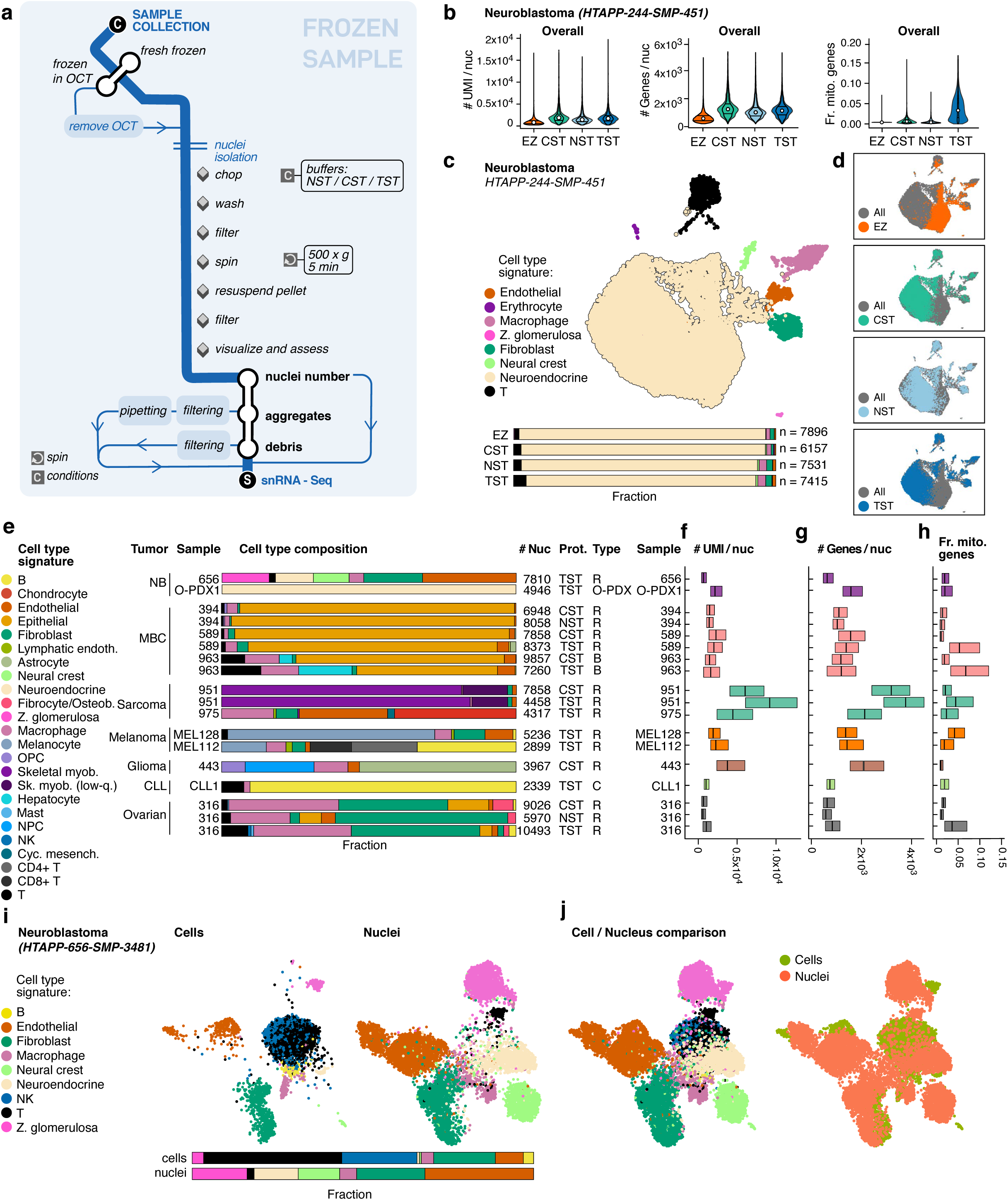
snRNA-Seq toolbox for frozen tumor samples. **(A)** Flow chart for collection and processing of frozen tumor samples. **(B-D)** Comparison of four nucleus isolation protocols in one neuroblastoma sample. (**B**) Variation in protocol performance. Distribution (median and first and third quartiles) of number of UMIs per nucleus, number of genes per nucleus, and fraction of UMIs per nucleus mapping to mitochondrial genes (*y* axes) in each protocol (*x* axis) across all nuclei in the dataset. **(C,D)** Protocols vary in diversity of cell types captured. (**C**) Top: UMAP embedding of single nucleus profiles (dots) from all four protocols, colored by assigned cell subset signature. Bottom: Proportion of cells from each subset in each of the four protocols. **(D)** UMAP embedding as in (c) colored by protocol. **(E-H)** Protocol comparison across tumor types. (**E**) Cell-type composition. Proportion of cells assigned with each cell subset signature (color) for each sample. R: Resection; B: Biopsy; A: Ascites; C: Cryopreserved; O-PDX: Orthotopic patient-derived xenograft. **(F-H)** QC metrics. Number of UMIs per nucleus, number of genes per nucleus, and fraction of UMIs per nucleus mapping to mitochondrial genes for each sample in (E). The median of the distributions along with the first and third quartiles are shown in (F-H). **(I-J)** scRNA-seq and snRNA-seq comparison in neuroblastoma. **(I)** Compositional differences between scRNA-Seq and snRNA-Seq of the same sample. UMAP embedding of scRNA-seq and snRNA-Seq profiles of the same sample combined by CCA(Butler et al., 2018) (**Methods**) showing profiles (dots) from either scRNA-seq (left) or snRNA-Seq (right), colored by assigned cell type signatures. Bottom: Proportion of cells in each subset in the two protocols. **(J)** Agreement in scRNA-seq and snRNA-seq intrinsic profiles. UMAP embedding as in (I) showing both scRNA-seq and snRNA-Seq profiles, colored by assigned cell type signatures (left, colored as in **(I)**) or by protocol (right).

Because in some tumor specimens the proportion of malignant cells is relatively low, we further optimized an immune-cell depletion strategy (**Methods**). Depletion of CD45^+^ expressing cells circumvents the need for enriching with specific surface markers (*e.g.*, EpCAM), which might otherwise bias the selection of specific cell populations, such as loss of representation of malignant cells undergoing EMT. Depletion applied to another NSCLC tumor sample (NSCLC17) increased the proportion of malignant epithelial cells from 26% in non-depleted scRNA-seq to 82% (**Fig. 2D, bottom, Supp. Fig. 4**), and the proportion of non-immune (CD45^-^) cells from 0.75% (by flow cytometry analysis) to 29.5% when applied to an ovarian ascites sample (**Fig. 2F, sample 727; Supp. Fig. 5**).

We also successfully applied the scRNA-Seq toolbox to much smaller core biopsy clinical samples. For example, in MBC, we applied the LD (Liberase TM and DNase I) protocol to a resection (HTAPP-254) and a biopsy (HTAPP-735) from lymph node metastases from two patients, yielding similarly successful QCs (**Fig. 2F-J, Supp. Fig. 6,7**). The resection and biopsy of the two patients had, however, different cellular compositions (**Fig. 2F**): a higher proportion of epithelial, endothelial, and fibroblast cells and a lower proportion of T cells in the biopsy compared to the resection. We similarly successfully profiled biopsies of MBC liver metastases (HTAPP-285, HTAPP-963) with the same protocol (**Fig. 2F-J, Supp. Fig. 8,9**), recovering some hepatocytes in addition to a similar set of cell types as was recovered in the lymph node biopsy (**Fig. 2F**). Thus, this protocol can be used across breast cancer metastases from different anatomical metastatic sites.

The scRNA-Seq toolbox performs well on samples obtained post-treatment, which can be challenging as a result of cell death and changes in cell type composition with treatment. We demonstrate this in profiling a pre-treatment diagnostic biopsy and post-treatment resection from the same neuroblastoma patient using the NB-C4 protocol (**Fig. 2F-J,** HTAPP-312-pre, HTAPP-312-post, **Supp. Fig. 10,11**). More cells but of fewer cell types were recovered in the pre-treatment biopsy (4,369 cells: neuroendocrine, T cells, and macrophages) than the post-treatment resection (786 cells: neuroendocrine, T cells, macrophages, as well as endothelial cells, and fibroblasts), consistent with observed post-treatment fibrosis. We tested an additional dissociation protocol in a neuroblastoma orthotopic patient-derived xenograft (O-PDX) sample (O-PDX1)(Stewart et al., 2017; Stewart et al., 2015), which is not expected to include non-malignant human cells, and indeed resulted in high quality malignant cell profiles (**Supp. Fig. 12**).

In addition to NSCLC, MBC, ovarian cancer ascites, and neuroblastoma samples (**Fig. 2F-J**, **Supplementary Figs. 2-13**), we established effective scRNA-Seq protocols for GBM, ovarian cancer, and CLL (**Fig. 2F-J, Supp. Fig. 14-16**). In particular, in CLL, we successfully recovered the expected cell types from a cryopreserved sample, containing viable cells. This reflects the increased resilience of immune cells to freezing compared to other cell types, also observed in other settings(Hermansen et al., 2018), and the lack of a dissociation step in CLL scRNA-Seq (**Methods**). Cryopreservation, however, can increase the proportion of damaged cells(Guillaumet-Adkins et al., 2017) and may not successfully recover all the malignant and other non-malignant cells in the tumor.

Thus, for frozen specimens from solid tumors, we optimized snRNA-Seq, focusing on different methods for nucleus isolation (**Fig. 3A**) across seven tumor types: MBC, neuroblastoma, ovarian cancer, pediatric sarcoma, melanoma, pediatric high-grade glioma, and CLL (**Fig. 1B, Supp. Fig. 1**). We initially divided larger samples or used multiple biopsies to compare four isolation methods (EZPrep(Habib et al., 2017), Nonidet™ P40 with salts and Tris (NST) [modified from Gao, R., et al (Gao et al., 2017)], CHAPS, with salts and Tris (CST) (Drokhlyansky et al., 2019), and Tween with salts and Tris (TST) (Drokhlyansky et al., 2019), which differ primarily in the mechanical force (*e.g.*, chopping or douncing), buffer, and/or detergent composition (**Fig. 3A**, **Methods**). Because in early tests EZPrep routinely underperformed CST, NST, and TST (data not shown), we only included it in initial comparisons (below). To evaluate protocols, we used the *post-hoc* computational criteria above, except we excluded the estimation of empty drops, because it was only developed and tested on single-cell RNA-seq data. We further customized scCloud for snRNA-Seq data, mapping reads to both exons and introns, and adapted the QC thresholds for transcript (UMI) and gene counts to reflect the lower expected mRNA content in nuclei (**Methods**). Experimentally, we added in-process light microscopy QCs to ensure complete nuclei isolation, and to estimate doublets, aggregates, and debris (**Fig. 3A, Methods**).

Overall, three nucleus isolation methods — TST, CST, and NST — had comparable performance based on the assessed nucleus quality (**Fig. 3B-H**), with TST typically yielding the greatest cell type diversity and number of nuclei per cell type, together with highest expression of mitochondrial genes, and NST typically having the fewest genes per nucleus and lowest diversity of types. For example, in neuroblastoma, testing each of the four protocols on a single resection sample (HTAPP-244) (**Fig. 3B-D, Supp. Fig. 17**) yielded a similar number of high-quality nuclei (7,896, 6,157, 7,531, and 7,415 for EZ, CST, NST, and TST, respectively), malignant cells with similarly detectable CNAs, and the expected cell types — with malignant neuroendocrine cells being the most prevalent (**Fig. 3C, Supp. Fig.17D, F-M**). However, nuclei prepared with the EZ protocol had lower numbers of UMIs and genes detected (**Fig. 3B**), while TST recovered more endothelial cells, fibroblasts, neural crest cells, and T cells than the other protocols (**Fig. 3C**). TST yielded a higher expression of mitochondrial genes (**Fig. 3B**), in this and all other tumors tested (**Fig. 3H**), since the nuclear membrane, ER, and ribosomes remain attached to the nucleus when using this method (Drokhlyansky et al., 2019). The same trends were preserved when down-sampling by the total number of sequencing reads (**Supp. Fig. 18**), as well as for cell-type specific QCs (**Supp. Fig. 17D**).

The CST, NST, and TST nucleus isolation methods had similar performance characteristics when tested with MBC, ovarian cancer, and pediatric sarcoma samples, with TST again providing the most diversity in cell types, especially in non-malignant cells. In MBC, we compared CST and NST in one metastatic brain resection (HTAPP-394), and CST and TST in another metastatic brain resection (HTAPP-589) and in a metastatic liver biopsy (HTAPP-963) (**Fig. 3E-H, Supp. Fig. 19-21**). In all cases, QC statistics (**Fig. 3F-H, Supp. Fig. 19-21A-D**) and CNA patterns **(Supp. Fig. 19-21G-H**) were similar between protocols, and nuclei from epithelial cells were the most prevalent (**Fig. 3E**). CST and NST captured a very similar distribution of cell types, while TST captured more non-malignant cells, including T cells (**Fig. 3E**) and a higher fraction of mitochondrial reads (**Fig. 3H**). In ovarian cancer, CST, NST, and TST recovered similar CNA patterns from the same sample (HTAPP-316, **Supp. Fig. 22**), but NST recovered fewer cells, genes per cell, and UMIs per cell (**Fig. 3E-G**), and had a lower cell type diversity, despite having greater overall sequencing depth, whereas TST captured the greatest cell type diversity (**Fig. 3E, Supp. Fig. 22A**). In a rhabdomyosarcoma sample (HTAPP-951), CST and TST captured the same cell types at similar proportions (**Fig. 3E**) and showed similar CNA patterns (**Supp. Fig. 23**).

Overall, we recommend the TST protocol for most tumor types, and CST for tumors from neuronal tissues, such as pediatric high-grade glioma (**Supp. Fig. 1, 24**). With the recommended protocols (**Fig. 1B**, right column), we profiled additional neuroblastoma tumors as well as Ewing sarcoma, melanoma, pediatric high-grade glioma, and CLL tumor samples — spanning biopsies, resections, and treated samples (**Fig. 1B, Fig. 3E-H**, **Supp. Fig. 24-30**). We also tested a pediatric rhabdomyosarcoma sample (HTAPP-951) by two different chemistries for droplet based snRNA-Seq (v2 *vs*. v3 from 10x Genomics, **Methods**), obtaining overall similar results in terms of cell types detected, an improved number of recovered *vs*. expected nuclei and higher complexity per nucleus in v3 (**Supp. Fig. 31**).

Finally, when we compared scRNA-Seq and snRNA-Seq by testing matching samples from the same specimen each in CLL, MBC, neuroblastoma, and O-PDX (**Fig. 3I-J, Supp. Fig. 32-35**), the methods typically recovered similar cell types with similar transcriptional profiles, but sometimes at varying proportions. In both neuroblastoma and MBC, immune cells were more prevalent in scRNA-Seq, and parenchymal (especially malignant) cells were more prevalent in snRNA-Seq (**Supp. Fig. 33,34**). Cell and nucleus profiles were comparable based on grouping together when using batch correction by canonical correlation analysis (CCA)(Butler et al., 2018) (**Methods**) (**Fig. 3J**, **Supp. Fig. 32-35**).

## Discussion

In conclusion, we developed a toolbox for processing fresh and frozen clinical tumor samples by single cell and single nucleus RNA-Seq, and demonstrated it across eight tumor types. For fresh tissues, we recommend testing 2-3 dissociation methods based on the tumor type, the tissue composition and the decision tree (**Fig. 1A**), and choose to apply the best performing protocol by assessing both experimental and computational QC metrics, and, if desired, adding a depletion step. For frozen tissues, we recommend testing the NST, TST, and CST protocols (**Fig. 3A**). While TST is often favorable due to its superior ability to capture the most diverse set of cells, in some tumors we recommend CST or NST (*e.g.*, CST for pediatric high-grade glioma, **Supp. Fig. 1**). CST also yields fewer mitochondrial reads, reducing sequencing cost. When possible, we recommend testing both scRNA-Seq and snRNA-Seq for the same tumor type, as the two approaches differ in the distribution of cell types detected. Processing frozen samples by snRNA-Seq allows studying many rare, unusual, and longitudinal banked tumor samples. Our toolbox will help researchers systematically profile additional human tumors, leading to a better understanding of tumor biology and ultimately to an era of precision medicine.

## Acknowledgments

We thank all patients and their families. We thank Jennifer Rood for help in editing, Anna Hupalowska for help with figure preparation, the clinical research teams who supplied the samples, Ellen Todres Gelfand, Nichole Straub, Laura DelloStritto, Karla Helvie and Shreevidya Periyasamy for project management, Miraj Patel, Ana Lako, and Scott J. Rodig for pathology review, the scientific team at Leidos Biomedical Research, Inc., Frederick National Laboratory for Cancer Research, especially Rachana Agarwal and Yuriko Mori, and the team at NCI, especially Shannon Hughes, Philipp Oberdoerffer, and Dinah Singer. This project has been funded in part with Federal funds from the National Cancer Institute, National Institutes of Health, Task Order No. HHSN261100039 under Contract No. HHSN261201500003I. The content of this publication does not necessarily reflect the views or policies of the Department of Health and Human Services, nor does mention of trade names, commercial products, or organizations imply endorsement by the U.S. Government. This project was also funded in part by the “START: Standardization of Single-Cell and Single-Nucleus RNA-seq Protocols for Tumors” project from the Chan Zuckerberg Initiative and the Klarman Cell Observatory. AR is an Investigator of the Howard Hughes Medical Institute. CJW is a Scholar of the Leukemia and Lymphoma Society. JK was supported by an EMBO Long-Term Fellowship (ALTF 738-2017).

AR is a founder of and equity holder in Celsius Therapeutics and an SAB member of Syros Pharmaceuticals, ThermoFisher Scientific and NeoGene. MS, OA, ED, ORR, and AR are co-inventors on patent applications filed by the Broad Institute to inventions relating to work in this manuscript, such as in PCT/US2018/060860 and US Provisional Application No. 62/745,259. CJW is a founder and member of the scientific advisory board of Neon Therapeutics, and receives research funding from Pharmacyclics. ASR is a consultant and equity holder in Celsius Therapeutics. FSH reports grants, personal fees from Bristol-Myers Squibb and Novartis; personal fees from Merck, EMD Serono, Takeda, Surface, Genentech/Roche, Compass Therapeutics, Apricity, Bayer, Aduro, Sanofi, Pfizer, Pionyr, Verastem, Torque, Rheos; In addition, Dr. Hodi has a patent Methods for Treating MICA-Related Disorders (#20100111973) with royalties paid, a patent Tumor antigens and uses thereof (#7250291) issued, a patent Angiopoiten-2 Biomarkers Predictive of Anti-immune checkpoint response (#20170248603) pending, a patent Compositions and Methods for Identification, Assessment, Prevention, and Treatment of Melanoma using PD-L1 Isoforms (#20160340407) pending, a patent Therapeutic peptides (#20160046716) pending, a patent Therapeutic Peptides (#20140004112) pending, a patent Therapeutic Peptides (#20170022275) pending, a patent Therapeutic Peptides (#20170008962) pending, a patent THERAPEUTIC PEPTIDES Therapeutic Peptides Patent number: 9402905 issued, and a patent METHODS OF USING PEMBROLIZUMAB AND TREBANANIB pending. RH has received research support from Novartis and Bristol-Myers and is a consultant for Tango Therapeutics. RB has received research support from Roche, Genentech, Merck, Siemens, Verastem, Gritstone, Epizyme, Medgenome, HTG and has equity in Navigation Sciences. This publication is part of the Human Cell Atlas - www.humancellatlas.org/publications/.

## Supplementary Figure Legends

**Supplementary Figure 1.**
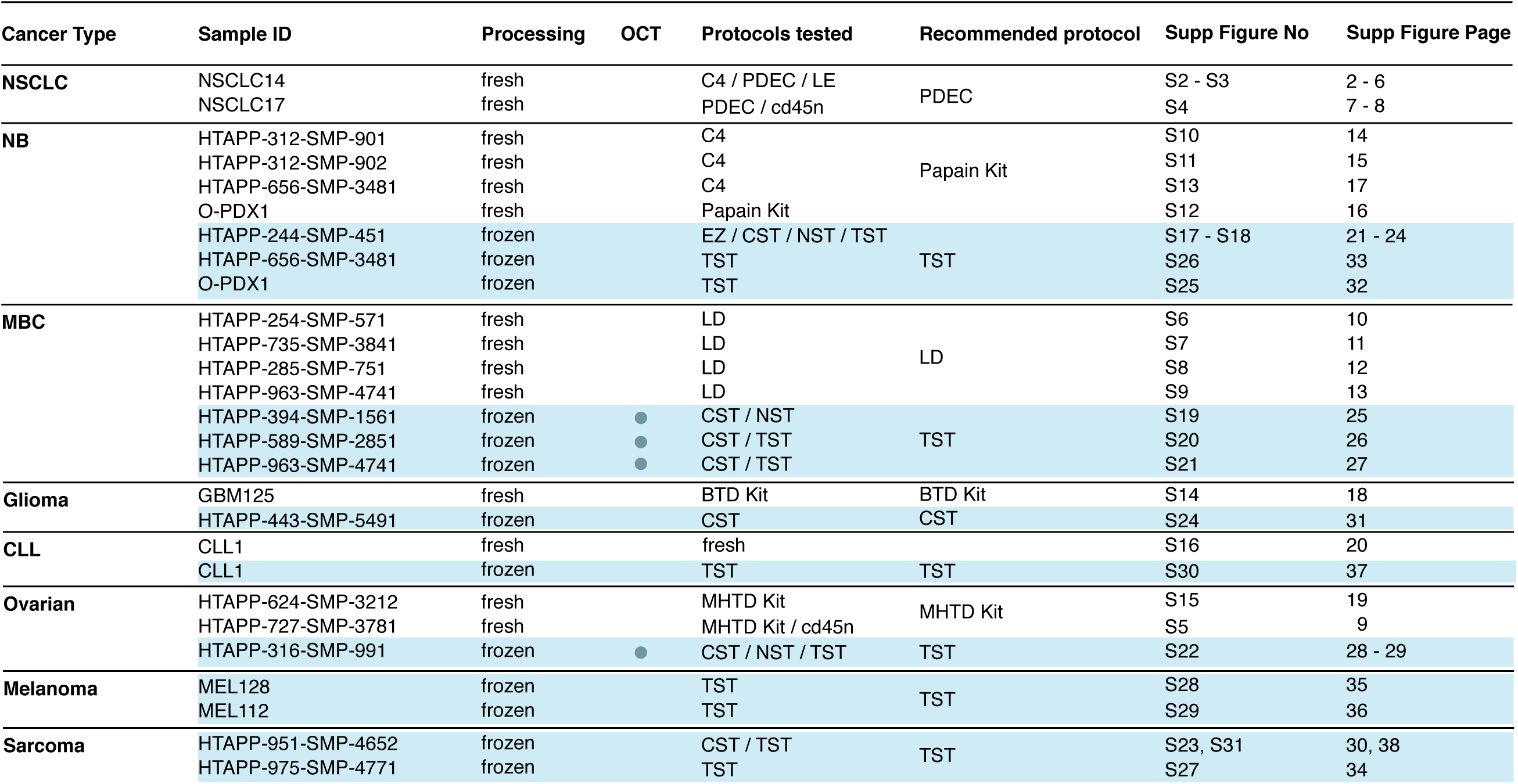
Overview of processed samples. Samples processed in this study are listed by tumor type (rows), along with their ID, tissue source (fresh or frozen, and OCT embedding), processing protocols tested, the recommended protocol, and the Supplementary Figure showing the sample’s analysis.

**Supplementary Figure 2.**
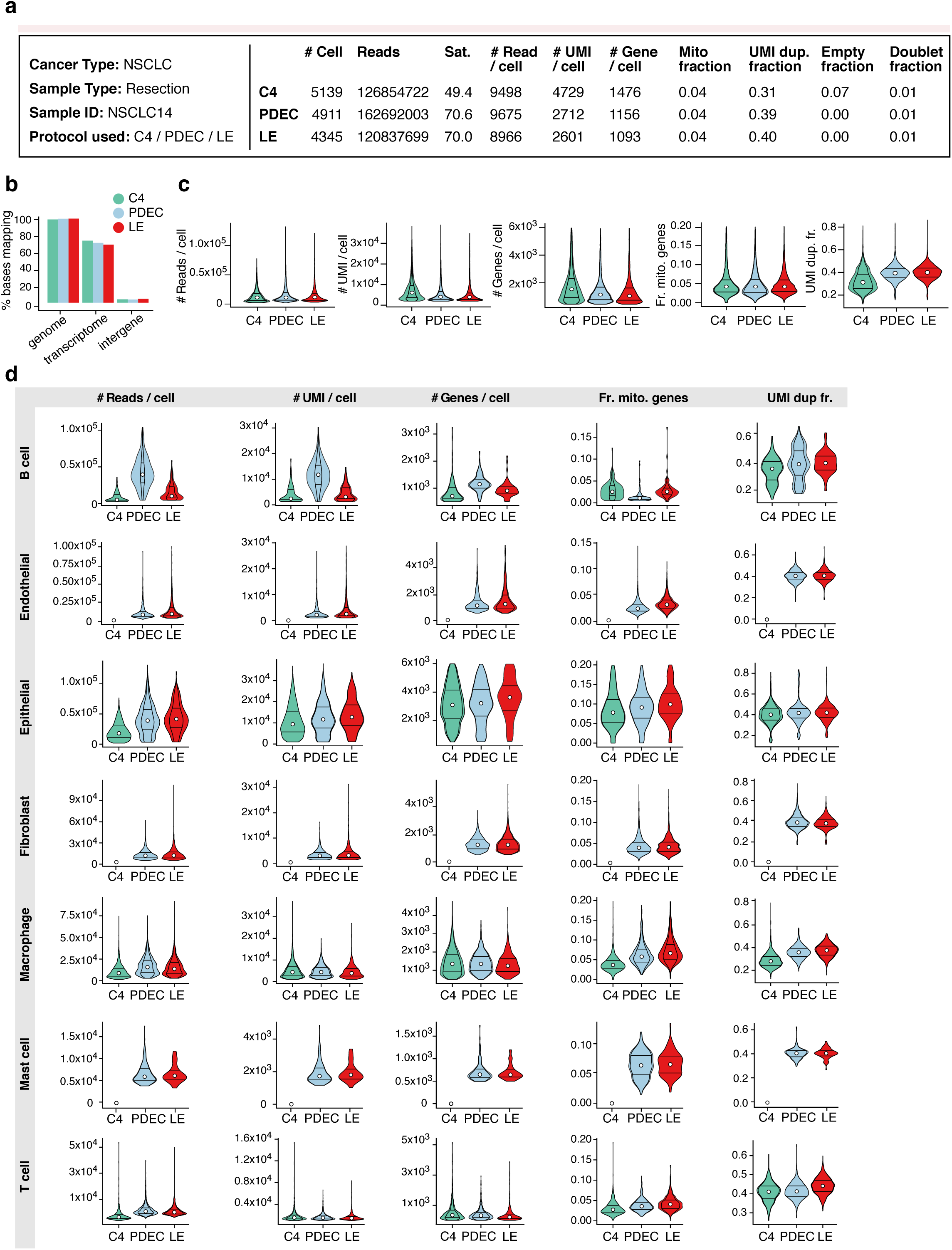
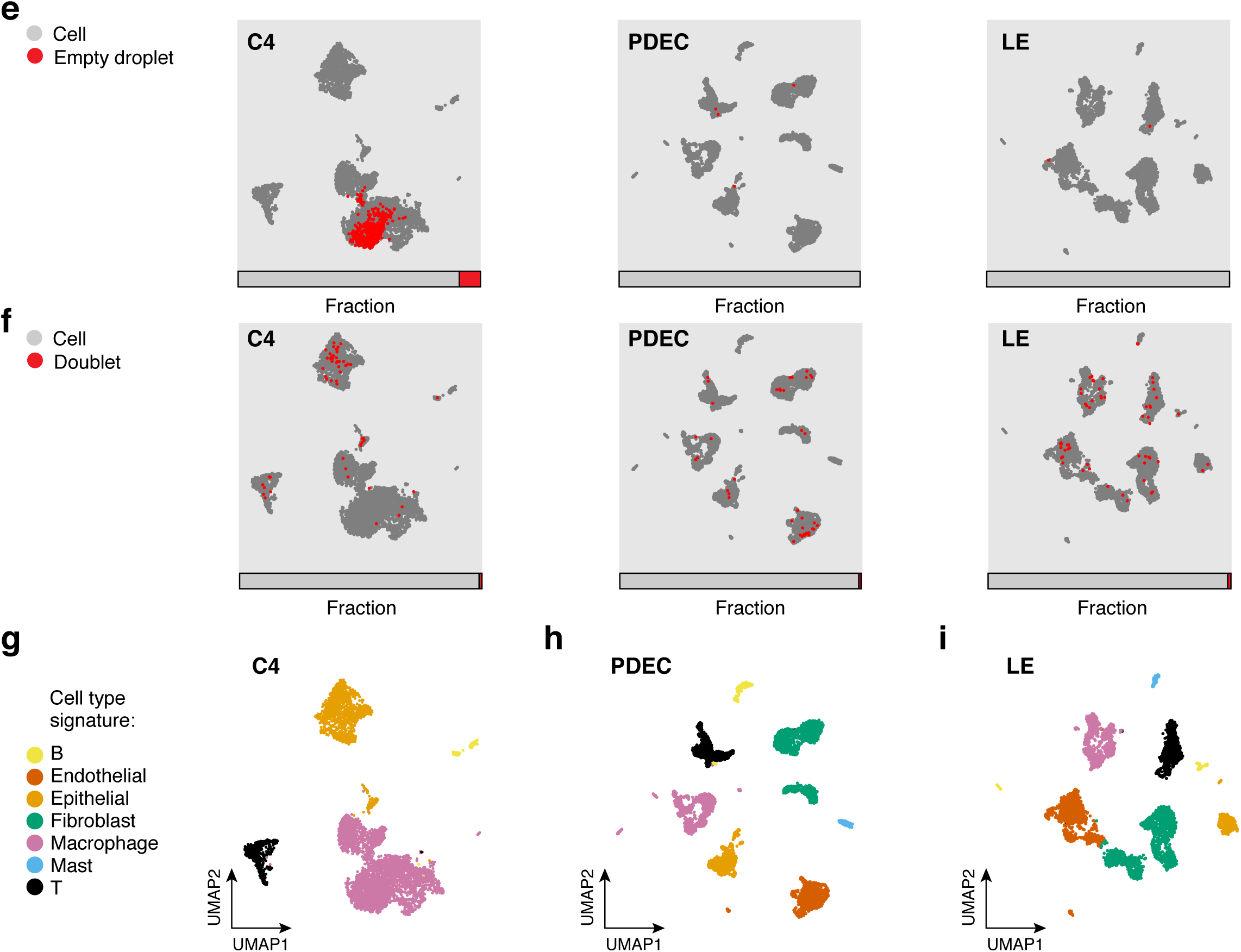
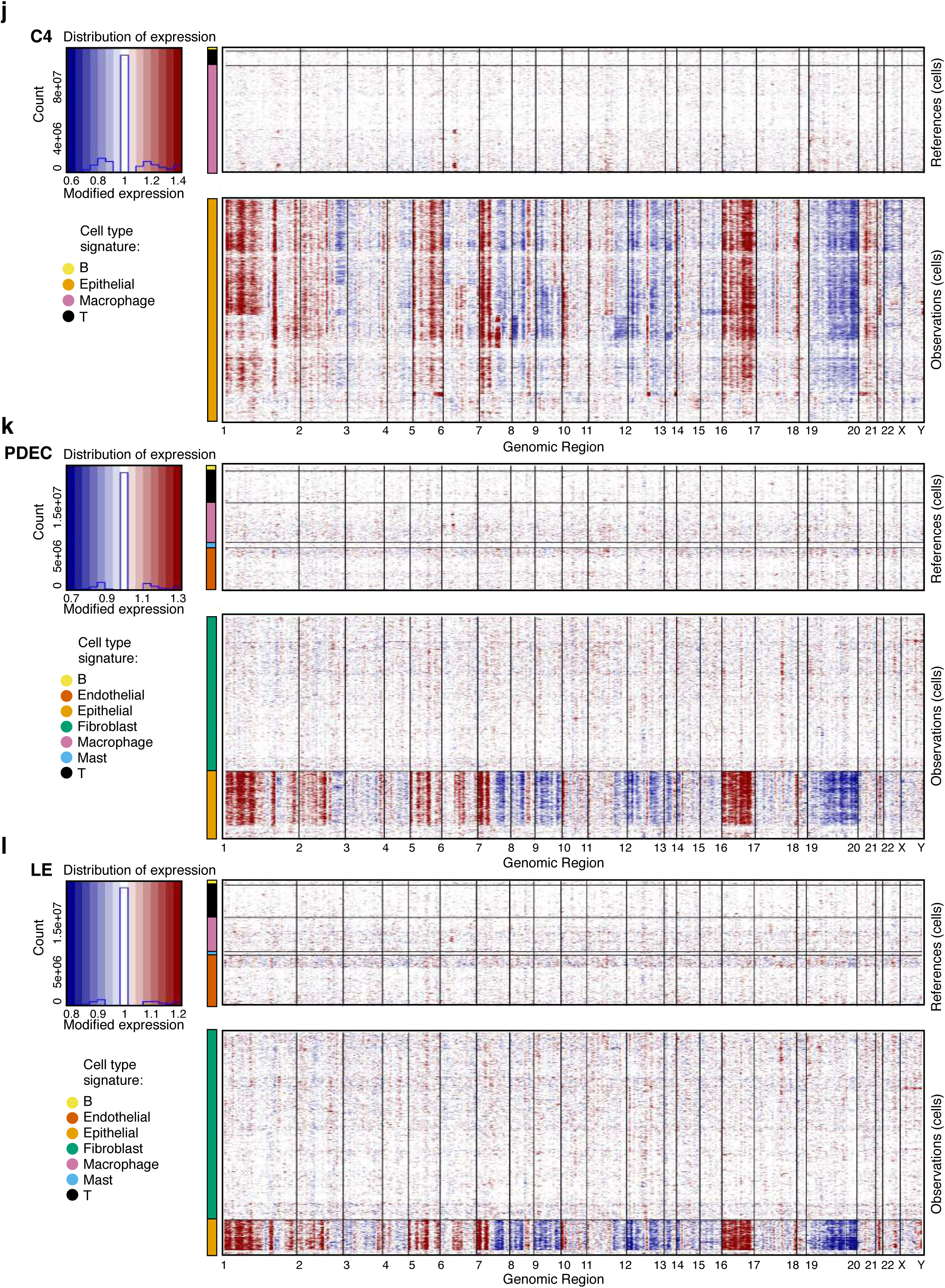
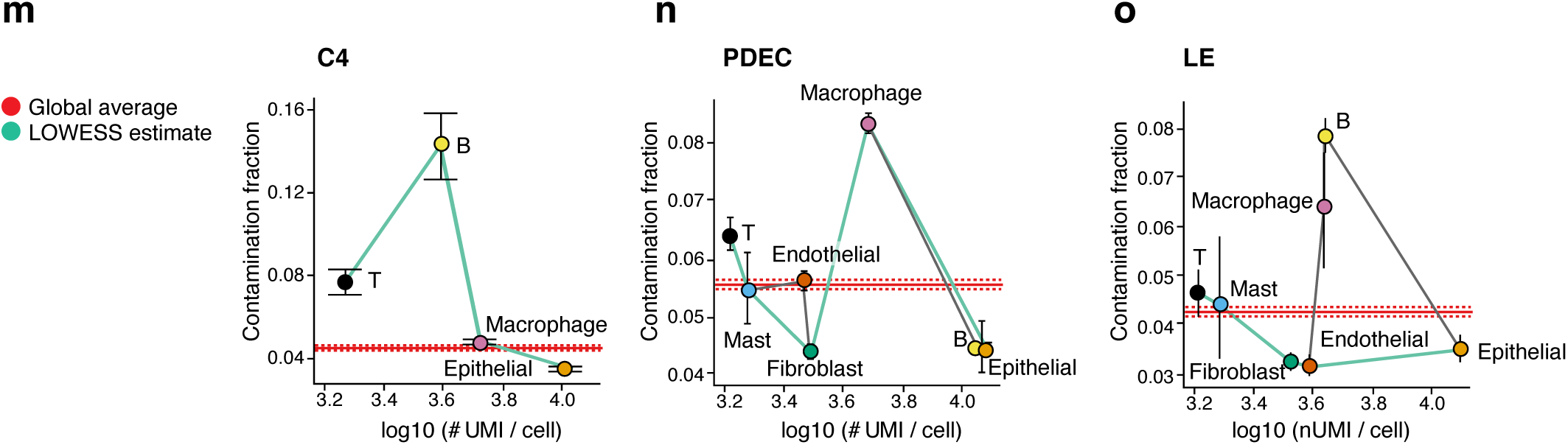
ScRNA-Seq protocol comparison for a single NSCLC sample. **(A)** Sample processing and QC overview. For each protocol, shown are the number of cells passing QC, and the number of sequencing reads and sequencing saturation across all cells. The remaining metrics are reported for those cells passing QC: the median number of reads per cell, median number of UMIs per cell, median number of genes per cell, median fraction of UMIs mapping to mitochondrial genes, median fraction of duplicated UMIs per cell, fraction of cell barcodes called as empty droplets, and fraction of cell barcodes called as doublets. **(B)** Read mapping QCs. The percent of bases in the sequencing reads (*y* axis) mapping to the genome, transcriptome, and intergenic regions (*x* axis) across the three protocols (colored bars). **(C-D)** Overall and cell types specific QCs. Distribution (median and first and third quartiles) of the number of reads per cell, number of UMIs per cell, number of genes per cell, fraction of UMIs mapping to mitochondrial genes in each cell, and fraction of duplicated UMIs per cell (*y* axes) in each of the three protocols (*x* axis), for all cells passing QC (**C**) and for cells passing QC from each cell type (**D**, rows). **(E,F)** Relation of empty droplets and doublets to cell types. UMAP embedding of single cell (grey), “empty droplet” (red, top), and doublet (red, bottom) profiles for each protocol. **(G-I)** Cell type assignment. UMAP embedding of single cell profiles from each protocol colored by assigned cell type signature. **(J-L)** Inferred CNA profiles. Chromosomal amplification (red) and deletion (blue) inferred in each chromosomal position (columns) across the single cells (rows). Top: reference cells not expected to contain CNA in this cancer type. Bottom: cells tested for CNA relative to the reference cells. Color bar: assigned cell type signature for each cell. **(M-O)** Ambient RNA estimates. SoupX(Young and Behjati, 2018) estimates of the fraction of RNA in each cell type derived from ambient RNA contamination (*y* axis), with cell types ordered by their mean number of UMIs/cell (*x* axis). Red line: global average of contamination fraction; Green line: LOWESS smoothed estimate of the contamination fraction within each cell type, along with the associated confidence interval.

**Supplementary Figure 3.**
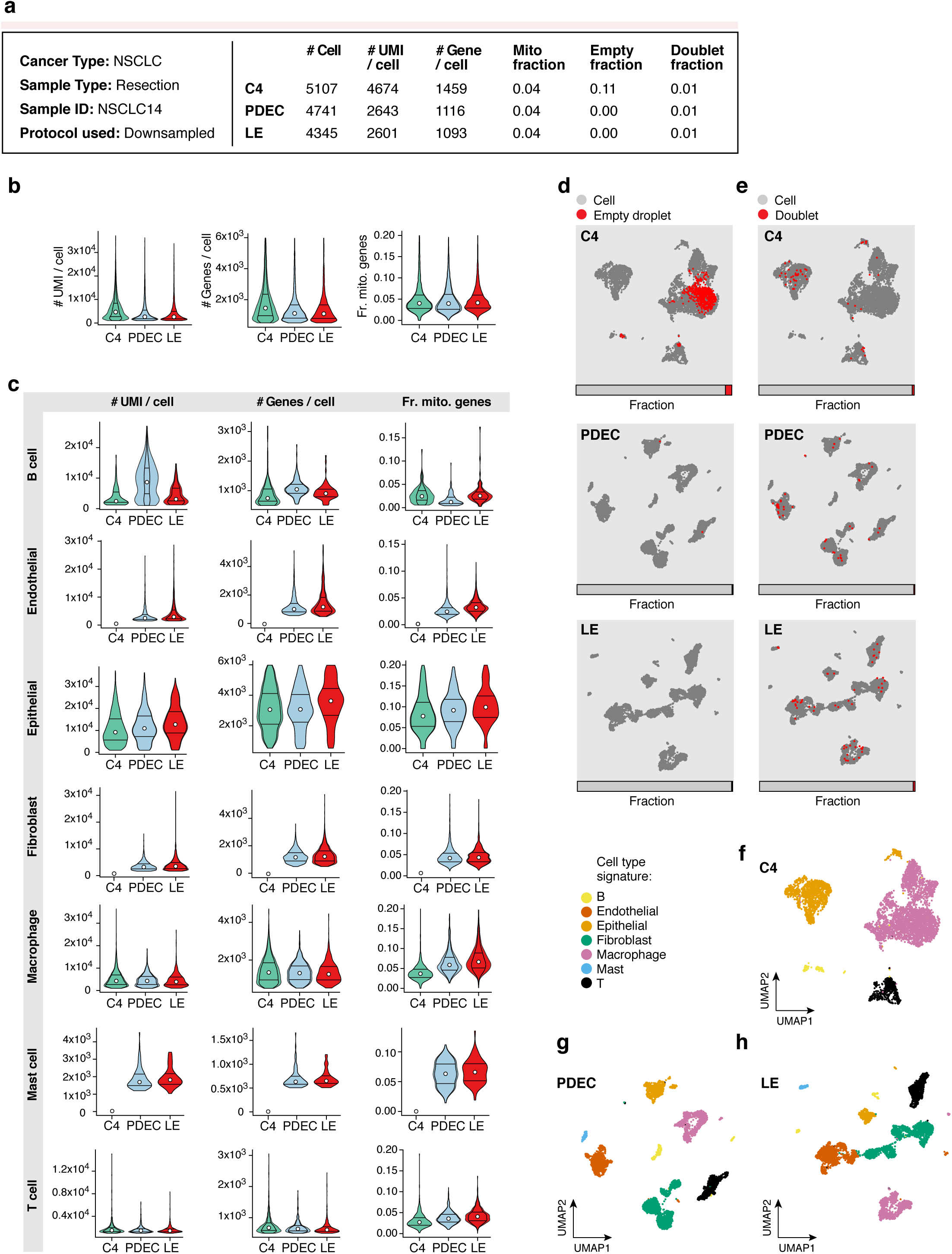
ScRNA-Seq protocol comparison for NSCLC following read down-sampling. Shown are analyses for NSCLC14 (as in **Supp.** Fig. 2), but after the total number of sequencing reads within each sample was down-sampled to match the protocol with the fewest total sequencing reads. **(A)** Sample processing and QC overview. For each protocol, shown are the number of cells passing QC. The remaining metrics are reported for those cells passing QC: median number of UMIs per cell, median number of genes per cell, median fraction of UMIs mapping to mitochondrial genes in each cell, fraction of cell barcodes called as empty droplets, and fraction of cell barcodes called as doublets. **(B,C)** Overall and cell types specific QCs. Distribution (median and first and third quartiles) of the number of UMIs per cell, number of genes per cell, and fraction of gene expression per cell from mitochondrial genes (*y* axes) in each of the three protocols (*x* axis), for all cells passing QC **(B)** and for cells from each cell type (**C,** rows). **(D,E)** Relation of empty droplets and doublets to cell types. UMAP embedding and fraction (horizontal bar) of single cell (grey), “empty droplet” (red, left), and doublet (red, right) profiles for each protocol **(F-H)** Cell type assignment. UMAP embedding of single cell profiles from each protocol colored by assigned cell type signature.

**Supplementary Figure 4.**
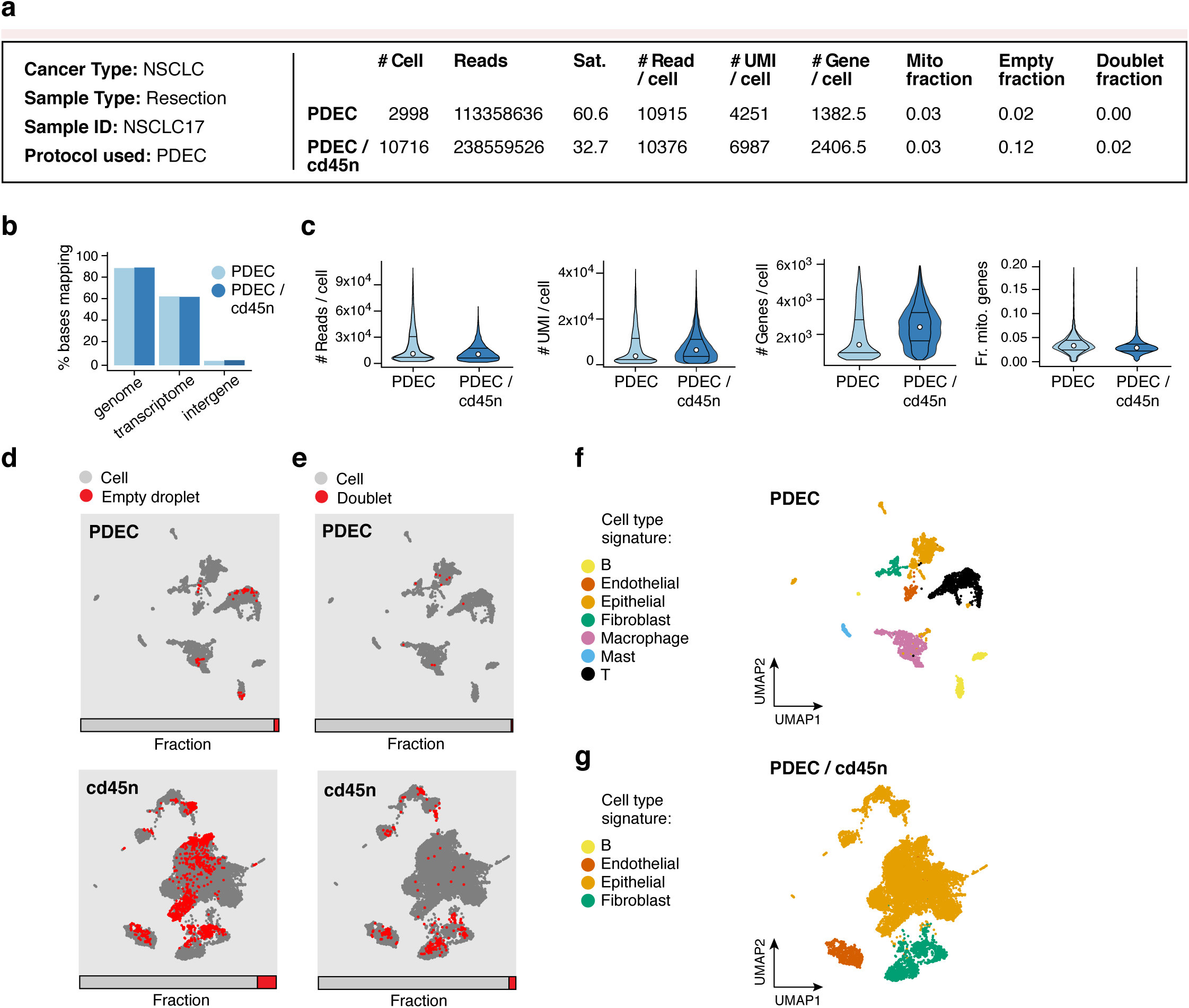
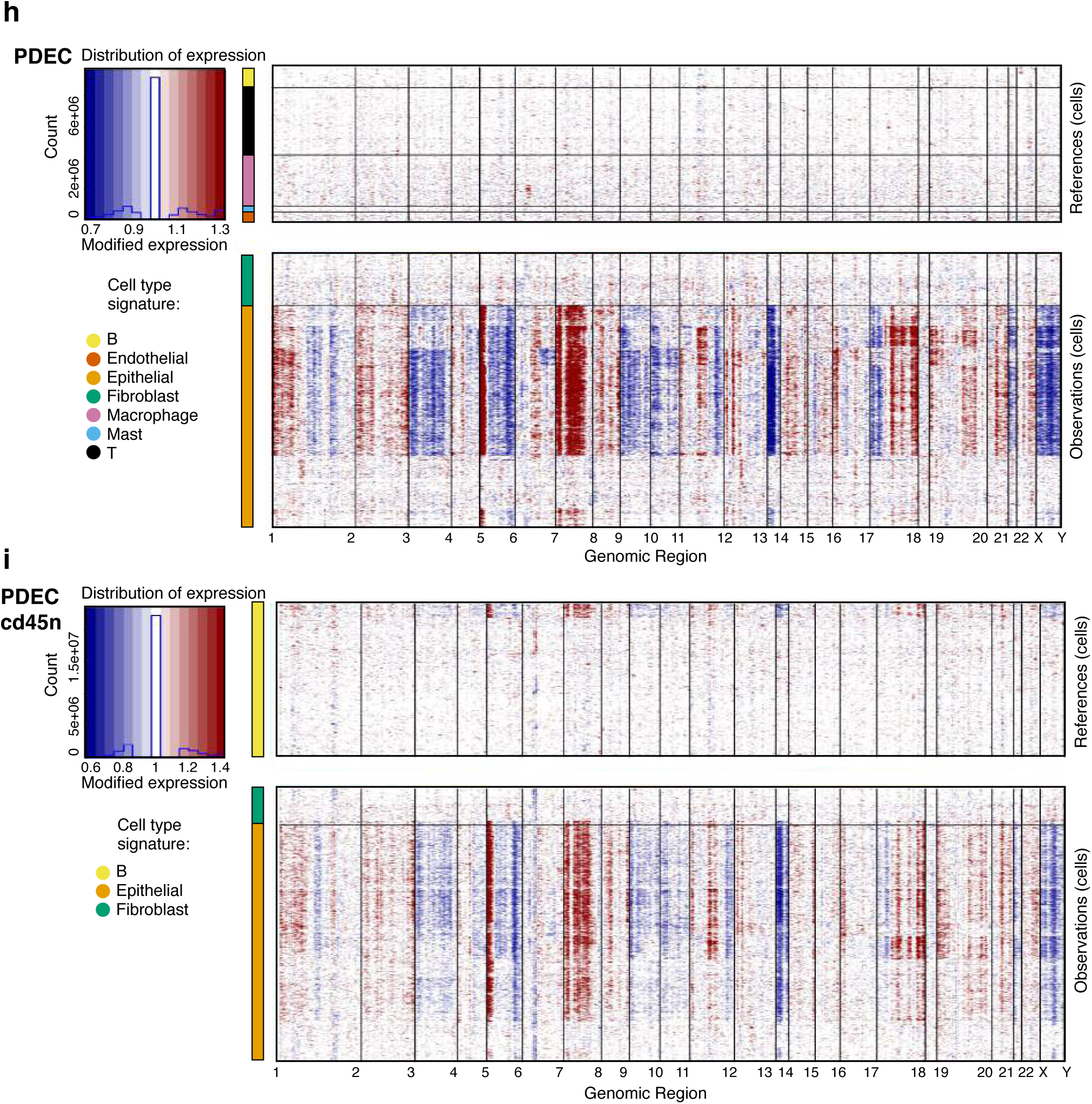
ScRNA-Seq depletion protocol enriches for malignant cells in freshly processed NSCLC. Cells were processed using the PDEC protocol or the PDEC protocol combined with depletion of CD45^+^ cells. **(A)** Sample processing and QC overview. For each protocol, shown are the number of cells passing QC, and the number of sequencing reads and sequencing saturation across all cells. The remaining metrics are reported for those cells passing QC: median number of reads per cell, median number of UMIs per cell, median number of genes per cell, median fraction of UMIs mapping to mitochondrial genes in each cell, fraction of cell barcodes called as empty droplets, and fraction of cell barcodes called as doublets. **(B)** Read mapping QCs. The percent of bases in the sequencing reads (*y* axis) mapping to the genome, transcriptome, and intergenic regions (*x* axis) in each of the two protocols (colored bars). **(C)** Overall QCs. Distribution (median and first and third quartiles) of the number of reads per cell, number of UMIs per cell, number of genes per cell, and fraction of UMIs mapping to mitochondrial genes in each cell (*y* axes) in each of the three protocols (*x* axis) for all cells passing QC. **(D,E)** Relation of empty droplets and doublets to cell types. UMAP embedding and fraction (horizontal bar) of single cell (grey), “empty droplet” (red, left) and doublet (red, right) profiles for each protocol. **(F-G)** Cell type assignment. UMAP embedding of single cell profiles from each protocol colored by assigned cell type signature. **(H-I)** Inferred CNA profiles for cells from each protocol. Chromosomal amplification (red) and deletion (blue) inferred in each chromosomal position (columns) across the single cells (rows). Top: reference cells not expected to contain CNA in this cancer type. Bottom: cells tested for CNA relative to the reference cells. Color bar: assigned cell type signature for each cell.

**Supplementary Figure 5.**
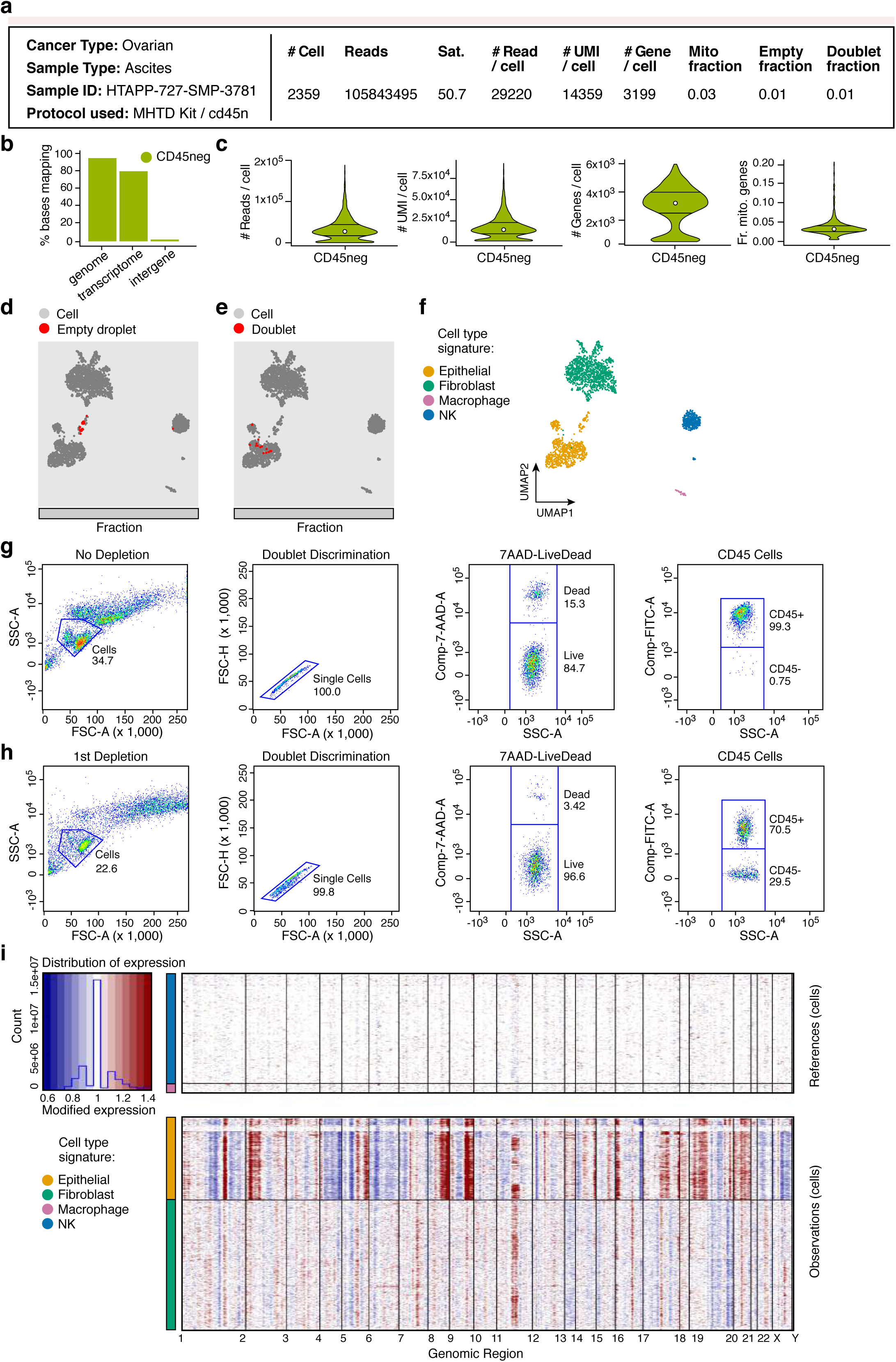
Application of CD45^+^ cell depletion scRNA-Seq protocol for processing ascites from ovarian cancer. **(A)** Sample processing and QC overview. Shown are the number of cells passing QC, and the number of sequencing reads and sequencing saturation across all cells. The remaining metrics are reported for those cells passing QC: median number of reads per cell, median number of UMIs per cell, median number of genes per cell, median fraction of UMIs mapping to mitochondrial genes in each cell, fraction of cell barcodes called as empty droplets, and fraction of cell barcodes called as doublets. **(B)** Read mapping QCs. The percent of bases in the sequencing reads (*y* axis) mapping to the genome, transcriptome, and intergenic regions (*x* axis). **(C)** Overall QCs. Distribution (median and first and third quartiles) of the number of reads per cell, number of UMIs per cell, number of genes per cell, and fraction of UMIs mapping to mitochondrial genes in each cell (*y* axes) for all cells passing QC. **(D,E)** Relation of empty droplets and doublets to cell types. UMAP embedding and fraction (horizontal bar) of single cell (grey), “empty droplet” (red, left) and doublet (red, right) profiles. **(F)** Cell type assignment. UMAP embedding of single cell profiles colored by assigned cell type signature. (**G,H)** Flow-cytometry comparison of single cells isolated (G) without or (H) with depletion of CD45^+^ cells. Cells were gated by FSC and SSC (first column), doublets removed using FSC-A and FSC-H (second column), live cells identified using 7AAD (third column), and the distribution of immune and non-immune cells quantified using a CD45 antibody (fourth column). Number of cells without and with depletion, respectively are: 10,000, 10,000 (1^st^ column), 3,468, 2,256 (2^nd^ column), 3,467, 2,251 (3^rd^ column), 2,936, 2,174 (4^th^ column). **(I)** Inferred CNA profiles for cells. Chromosomal amplification (red) and deletion (blue) inferred in each chromosomal position (columns) across the single cells (rows). Top: reference cells not expected to contain CNA in this cancer type. Bottom: cells tested for CNA relative to the reference cells. Color bar: assigned cell type signature for each cell.

**Supplementary Figure 6.**
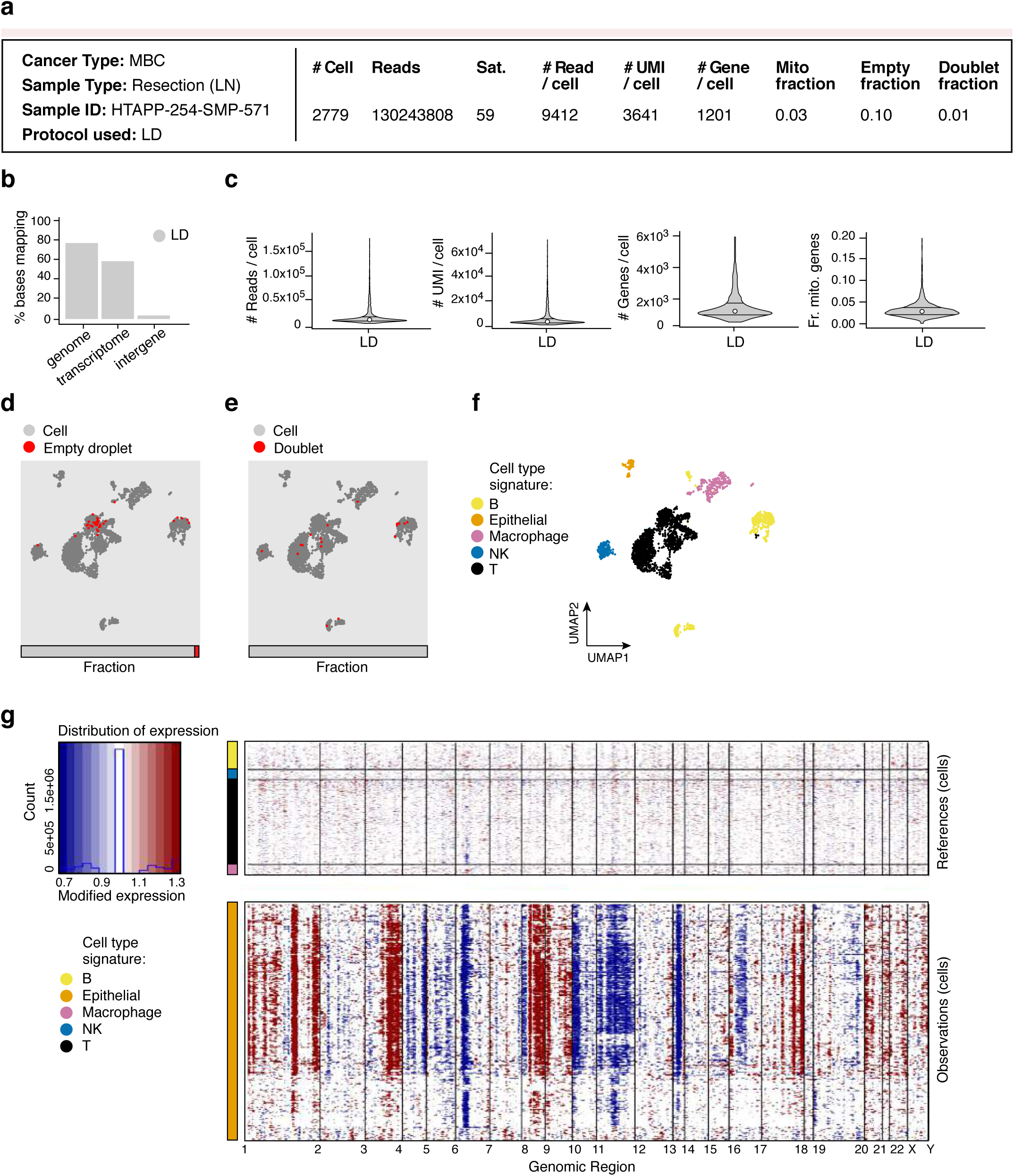
Evaluation of scRNA-Seq protocol for lymph node resection of metastatic breast cancer. **(A)** Sample processing and QC overview. Shown are the number of cells passing QC, and the number of sequencing reads and sequencing saturation across all cells. The remaining metrics are reported for those cells passing QC: median number of reads per cell, median number of UMIs per cell, median number of genes per cell, median fraction of UMIs mapping to mitochondrial genes in each cell, fraction of cell barcodes called as empty droplets, and fraction of cell barcodes called as doublets. **(B)** Read mapping QCs. The percent of bases in the sequencing reads (*y* axis) mapping to the genome, transcriptome, and intergenic regions (*x* axis). **(C)** Overall QCs. Distribution (median and first and third quartiles) of the number of reads per cell, number of UMIs per cell, number of genes per cell, and fraction of UMIs mapping to mitochondrial genes in each cell (*y* axes) for all cells passing QC. **(D,E)** Relation of empty droplets and doublets to cell types. UMAP embedding and fraction (horizontal bar) of single cell (grey), “empty droplet” (red, left) and doublet (red, right) profiles. **(F)** Cell type assignment. UMAP embedding of single cell profiles colored by assigned cell type signature. **(G)** Inferred CNA profiles for cells. Chromosomal amplification (red) and deletion (blue) inferred in each chromosomal position (columns) across the single cells (rows). Top: reference cells not expected to contain CNA in this cancer type. Bottom: cells tested for CNA relative to the reference cells. Color bar: assigned cell type signature for each cell.

**Supplementary Figure 7.**
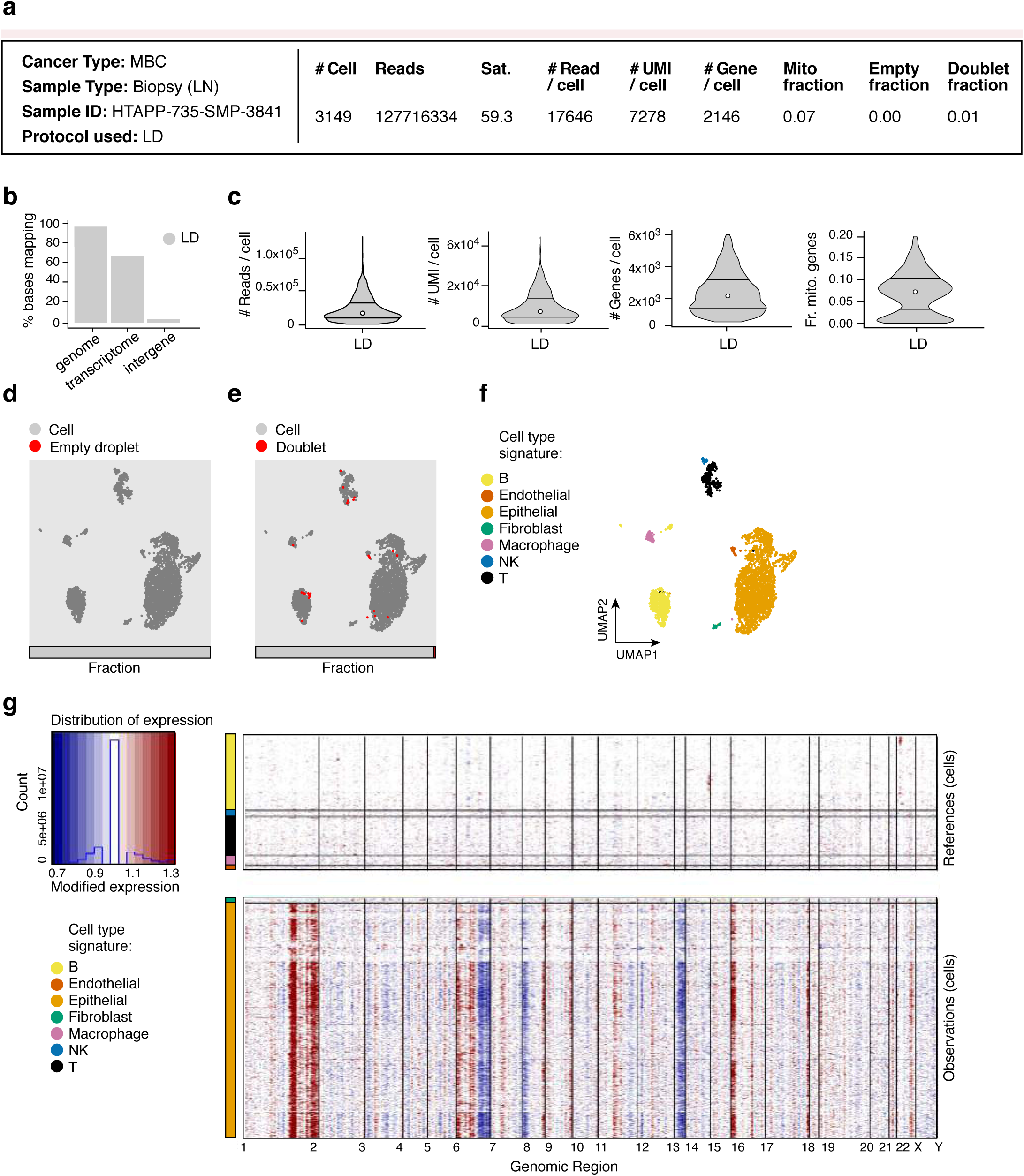
Evaluation of scRNA-Seq protocol for lymph node biopsy of metastatic breast cancer. **(A)** Sample processing and QC overview. Shown are the number of cells passing QC, and the number of sequencing reads and sequencing saturation across all cells. The remaining metrics are reported for those cells passing QC: median number of reads per cell, median number of UMIs per cell, median number of genes per cell, median fraction of fraction of UMIs mapping to mitochondrial genes in each cell, fraction of cell barcodes called as empty droplets, and fraction of cell barcodes called as doublets. **(B)** Read mapping QCs. The percent of bases in the sequencing reads (*y* axis) mapping to the genome, transcriptome, and intergenic regions (*x* axis). **(C)** Overall QCs. Distribution (median and first and third quartiles) of the number of reads per cell, number of UMIs per cell, number of genes per cell, and fraction of UMIs mapping to mitochondrial genes in each cell (*y* axes) for all cells passing QC. **(D,E)** Relation of empty droplets and doublets to cell types. UMAP embedding and fraction (horizontal bar) of single cell (grey), “empty droplet” (red, left) and doublet (red, right) profiles. **(F)** Cell type assignment. UMAP embedding of single cell profiles colored by assigned cell type signature. **(G)** Inferred CNA profiles for cells. Chromosomal amplification (red) and deletion (blue) inferred in each chromosomal position (columns) across the single cells (rows). Top: reference cells not expected to contain CNA in this cancer type. Bottom: cells tested for CNA relative to the reference cells. Color bar: assigned cell type signature for each cell.

**Supplementary Figure 8.**
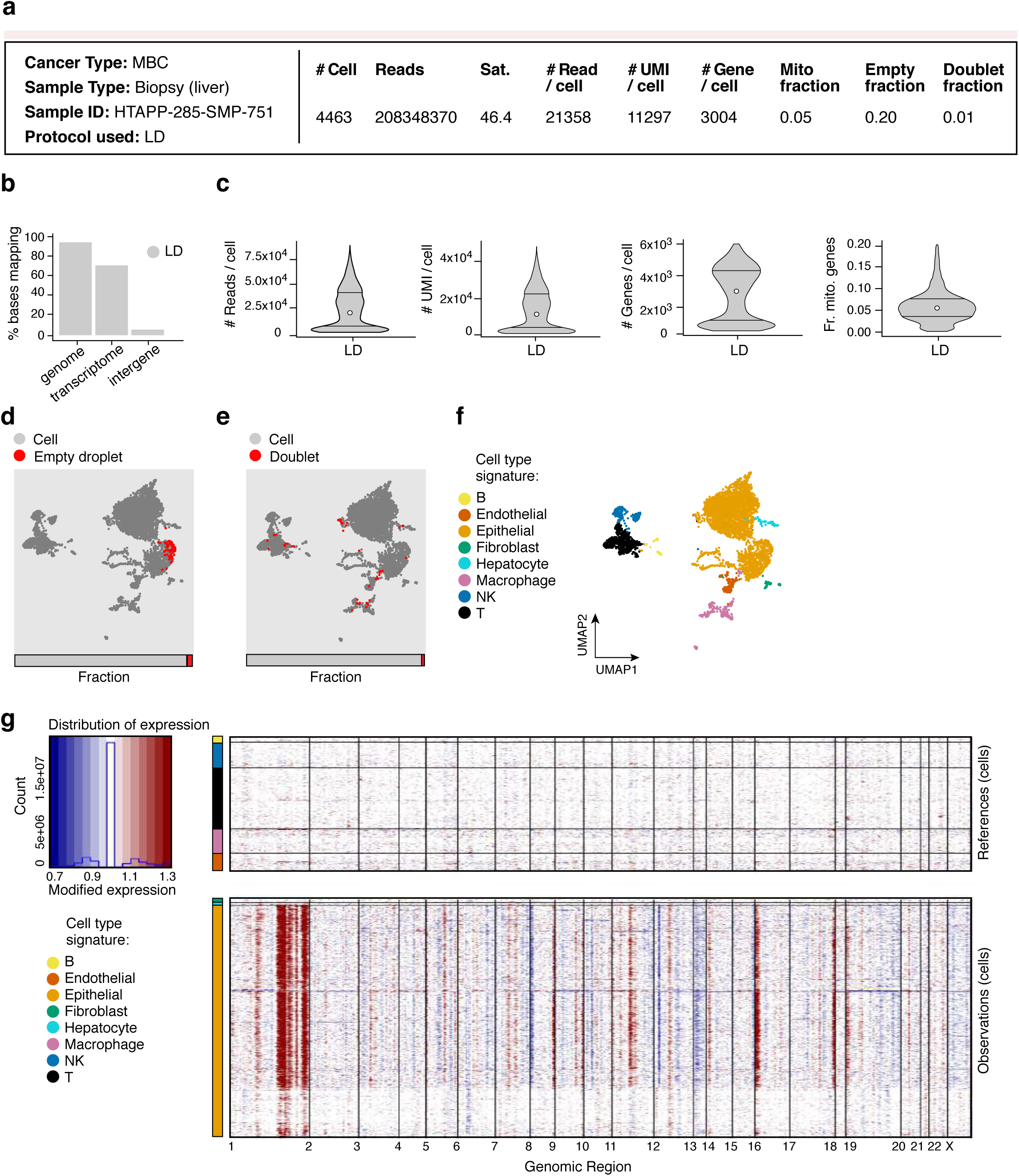
Evaluation of scRNA-Seq protocol for liver biopsy of metastatic breast cancer. **(A)** Sample processing and QC overview. Shown are the number of cells passing QC, and the number of sequencing reads and sequencing saturation across all cells. The remaining metrics are reported for those cells passing QC: median number of reads per cell, median number of UMIs per cell, median number of genes per cell, median fraction of UMIs mapping to mitochondrial genes in each cell, fraction of cell barcodes called as empty droplets, and fraction of cell barcodes called as doublets. **(B)** Read mapping QCs. The percent of bases in the sequencing reads (*y* axis) mapping to the genome, transcriptome, and intergenic regions (*x* axis). **(C)** Overall QCs. Distribution (median and first and third quartiles) of the number of reads per cell, number of UMIs per cell, number of genes per cell, and fraction of UMIs mapping to mitochondrial genes in each cell (*y* axes) for all cells passing QC. **(D,E)** Relation of empty droplets and doublets to cell types. UMAP embedding and fraction (horizontal bar) of single cell (grey), “empty droplet” (red, left) and doublet (red, right) profiles. **(F)** Cell type assignment. UMAP embedding of single cell profiles colored by assigned cell type signature. **(G)** Inferred CNA profiles for cells. Chromosomal amplification (red) and deletion (blue) inferred in each chromosomal position (columns) across the single cells (rows). Top: reference cells not expected to contain CNA in this cancer type. Bottom: cells tested for CNA relative to the reference cells. Color bar: assigned cell type signature for each cell.

**Supplementary Figure 9.**
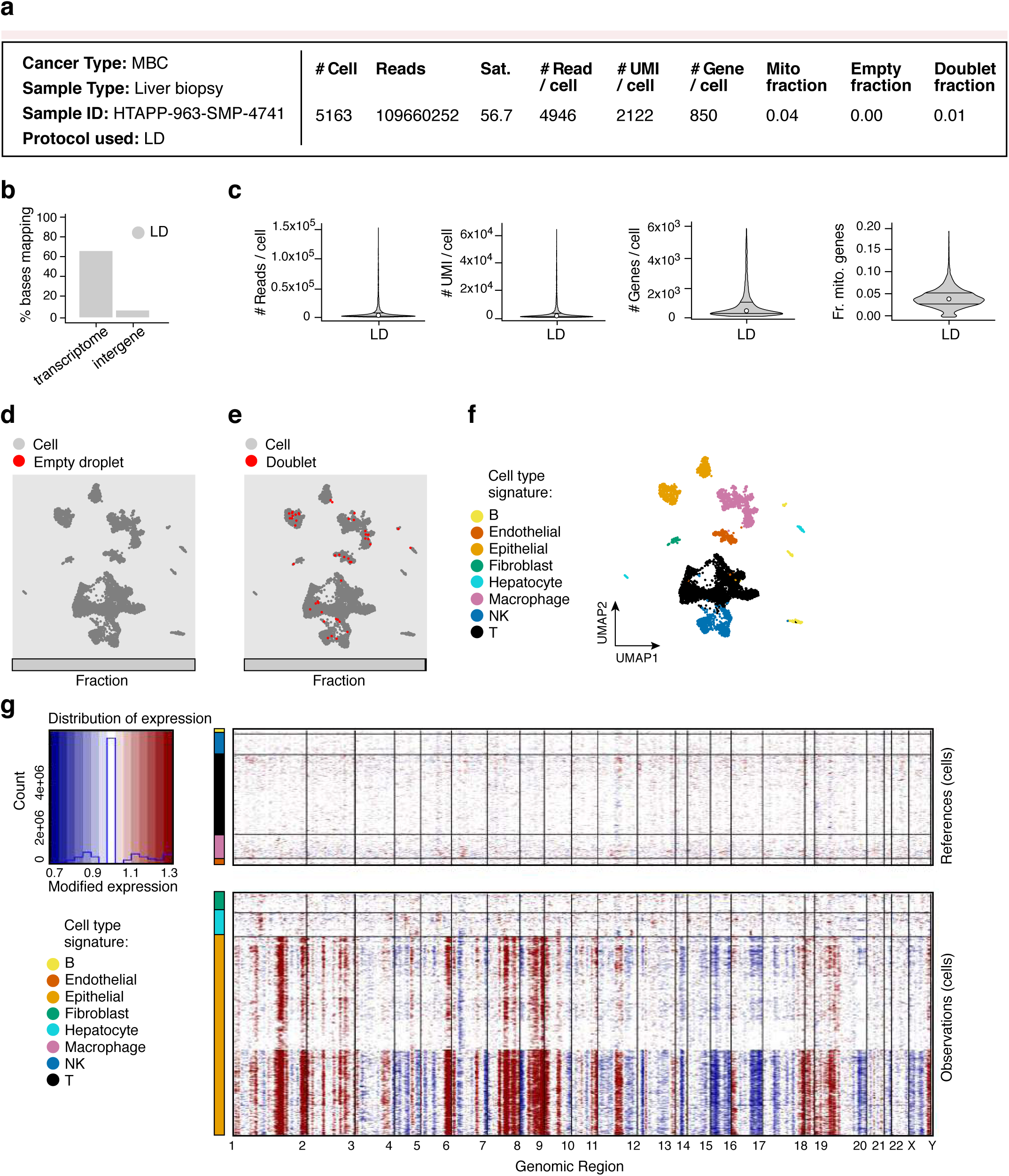
Evaluation of scRNA-Seq protocol for liver biopsy of metastatic breast cancer. **(A)** Sample processing and QC overview. Shown are the number of cells passing QC, and the number of sequencing reads and sequencing saturation across all cells. The remaining metrics are reported for those cells passing QC: median number of reads per cell, median number of UMIs per cell, median number of genes per cell, median fraction of UMIs mapping to mitochondrial genes in each cell, fraction of cell barcodes called as empty droplets, and fraction of cell barcodes called as doublets. **(B)** Read mapping QCs. The percent of bases in the sequencing reads (*y* axis) mapping to the transcriptome and intergenic regions (*x* axis). **(C)** Overall QCs. Distribution (median and first and third quartiles) of the number of reads per cell, number of UMIs per cell, number of genes per cell, and fraction of UMIs mapping to mitochondrial genes in each cell (*y* axes) for all cells passing QC. **(D,E)** Relation of empty droplets and doublets to cell types. UMAP embedding and fraction (horizontal bar) of single cell (grey), “empty droplet” (red, left) and doublet (red, right) profiles. **(F)** Cell type assignment. UMAP embedding of single cell profiles colored by assigned cell type signature. **(G)** Inferred CNA profiles for cells. Chromosomal amplification (red) and deletion (blue) inferred in each chromosomal position (columns) across the single cells (rows). Top: reference cells not expected to contain CNA in this cancer type. Bottom: cells tested for CNA relative to the reference cells. Color bar: assigned cell type signature for each cell.

**Supplementary Figure 10.**
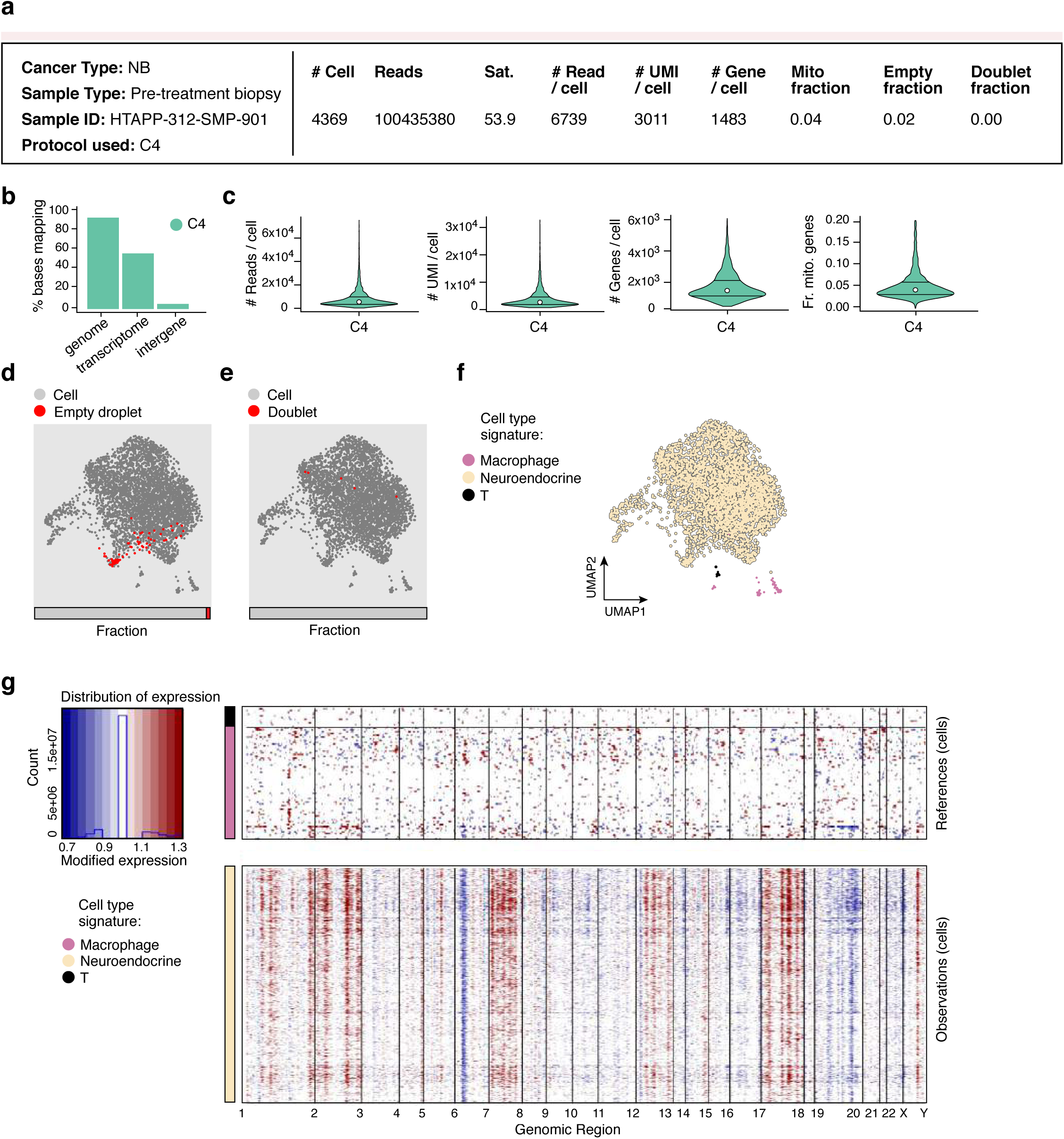
Evaluation of scRNA-Seq protocol for pre-treatment biopsy of neuroblastoma. **(A)** Sample processing and QC overview. Shown are the number of cells passing QC, and the number of sequencing reads and sequencing saturation across all cells. The remaining metrics are reported for those cells passing QC: median number of reads per cell, median number of UMIs per cell, median number of genes per cell, median fraction of UMIs mapping to mitochondrial genes in each cell, fraction of cell barcodes called as empty droplets, and fraction of cell barcodes called as doublets. **(B)** Read mapping QCs. The percent of bases in the sequencing reads (*y* axis) mapping to the genome, transcriptome, and intergenic regions (*x* axis). **(C)** Overall QCs. Distribution (median and first and third quartiles) of the number of reads per cell, number of UMIs per cell, number of genes per cell, and fraction of UMIs mapping to mitochondrial genes in each cell (*y* axes) for all cells passing QC. **(D,E)** Relation of empty droplets and doublets to cell types. UMAP embedding and fraction (horizontal bar) of single cell (grey), “empty droplet” (red, left) and doublet (red, right) profiles. **(F)** Cell type assignment. UMAP embedding of single cell profiles colored by assigned cell type signature. **(G)** Inferred CNA profiles for cells. Chromosomal amplification (red) and deletion (blue) inferred in each chromosomal position (columns) across the single cells (rows). Top: reference cells not expected to contain CNA in this cancer type. Bottom: cells tested for CNA relative to the reference cells. Color bar: assigned cell type signature for each cell.

**Supplementary Figure 11.**
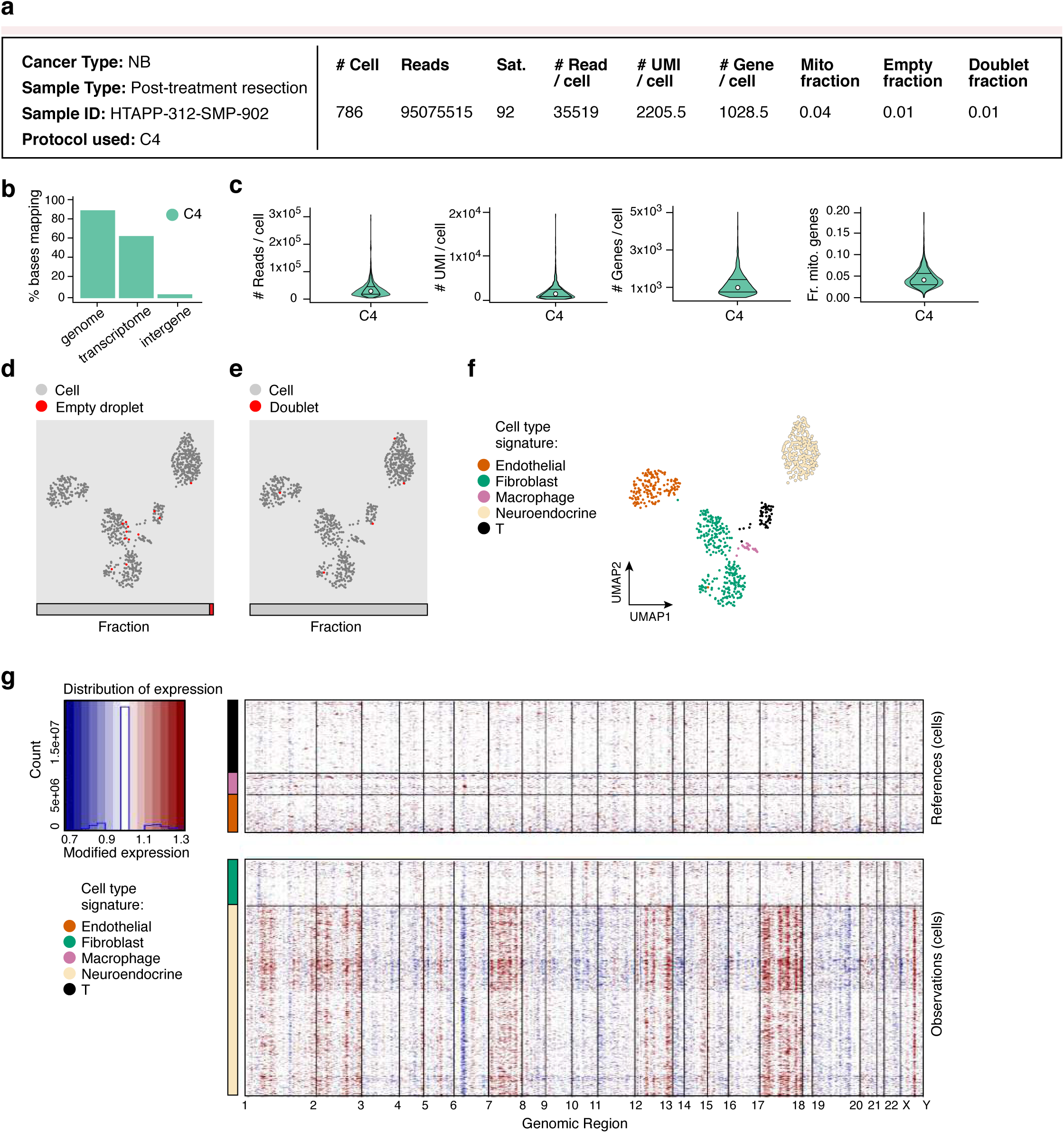
Evaluation of scRNA-Seq protocol for post-treatment resection of neuroblastoma. **(A)** Sample processing and QC overview. Shown are the number of cells passing QC, and the number of sequencing reads and sequencing saturation across all cells. The remaining metrics are reported for those cells passing QC: median number of reads per cell, median number of UMIs per cell, median number of genes per cell, median fraction of UMIs mapping to mitochondrial genes in each cell, fraction of cell barcodes called as empty droplets, and fraction of cell barcodes called as doublets. **(B)** Read mapping QCs. The percent of bases in the sequencing reads (*y* axis) mapping to the genome, transcriptome, and intergenic regions (*x* axis). **(C)** Overall QCs. Distribution (median and first and third quartiles) of the number of reads per cell, number of UMIs per cell, number of genes per cell, and fraction of UMIs mapping to mitochondrial genes in each cell (*y* axes) for all cells passing QC. **(D,E)** Relation of empty droplets and doublets to cell types. UMAP embedding and fraction (horizontal bar) of single cell (grey), “empty droplet” (red, left) and doublet (red, right) profiles. **(F)** Cell type assignment. UMAP embedding of single cell profiles colored by assigned cell type signature. **(G)** Inferred CNA profiles for cells. Chromosomal amplification (red) and deletion (blue) inferred in each chromosomal position (columns) across the single cells (rows). Top: reference cells not expected to contain CNA in this cancer type. Bottom: cells tested for CNA relative to the reference cells. Color bar: assigned cell type signature for each cell.

**Supplementary Figure 12.**
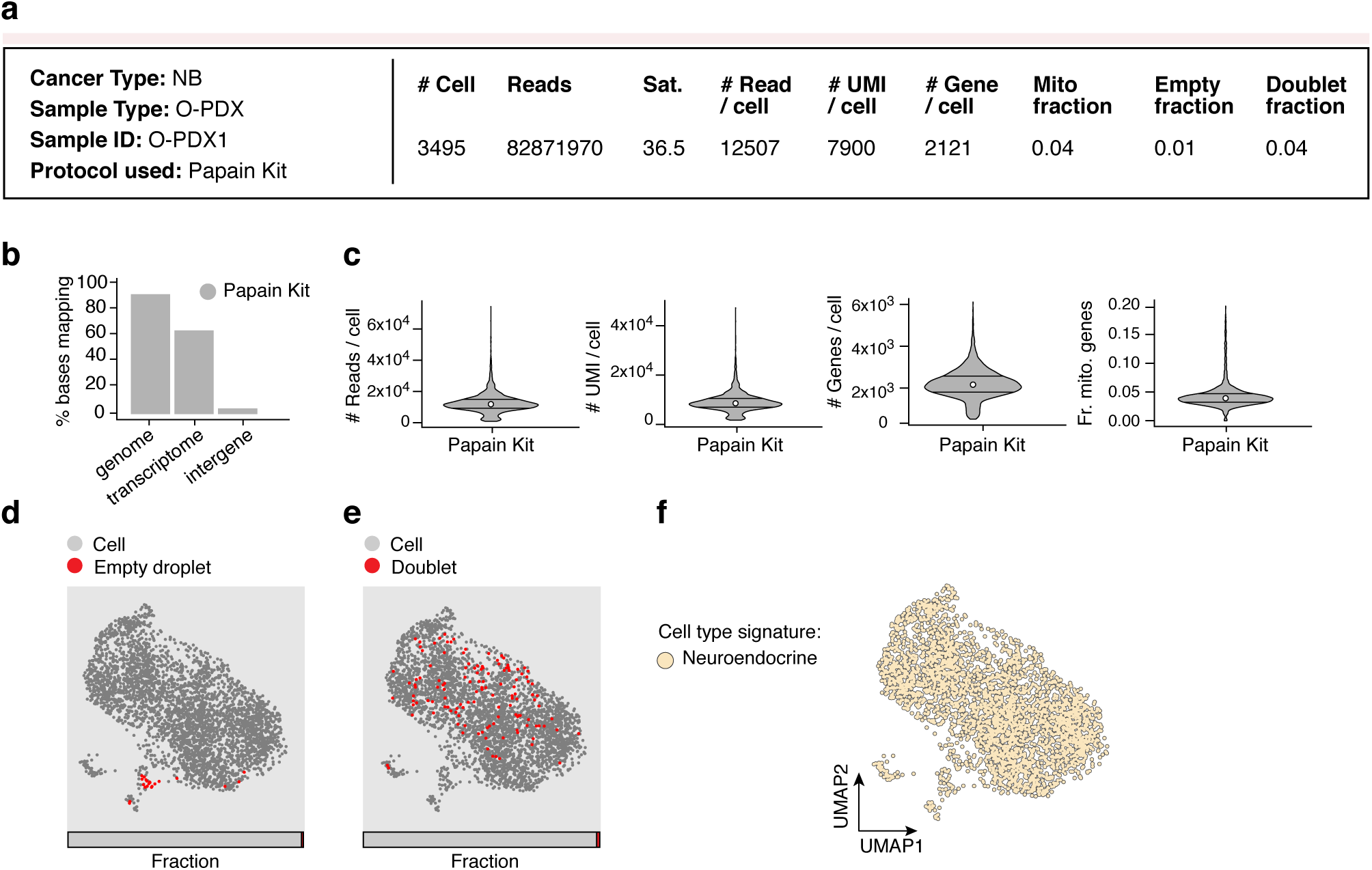
Evaluation of scRNA-Seq protocol for neuroblastoma O-PDX. **(A)** Sample processing and QC overview. Shown are the number of cells passing QC, and the number of sequencing reads and sequencing saturation across all cells. The remaining metrics are reported for those cells passing QC: median number of reads per cell, median number of UMIs per cell, median number of genes per cell, median fraction of UMIs mapping to mitochondrial genes in each cell, fraction of cell barcodes called as empty droplets, and fraction of cell barcodes called as doublets. **(B)** Read mapping QCs. The percent of bases in the sequencing reads (*y* axis) mapping to the genome, transcriptome, and intergenic regions (*x* axis). **(C)** Overall QCs. Distribution (median and first and third quartiles) of the number of reads per cell, number of UMIs per cell, number of genes per cell, and fraction of UMIs mapping to mitochondrial genes in each cell (*y* axes) for all cells passing QC. **(D,E)** Relation of empty droplets and doublets to cell types. UMAP embedding and fraction (horizontal bar) of single cell (grey), “empty droplet” (red, left) and doublet (red, right) profiles. **(F)** Cell type assignment. UMAP embedding of single cell profiles colored by assigned cell type signature.

**Supplementary Figure 13.**
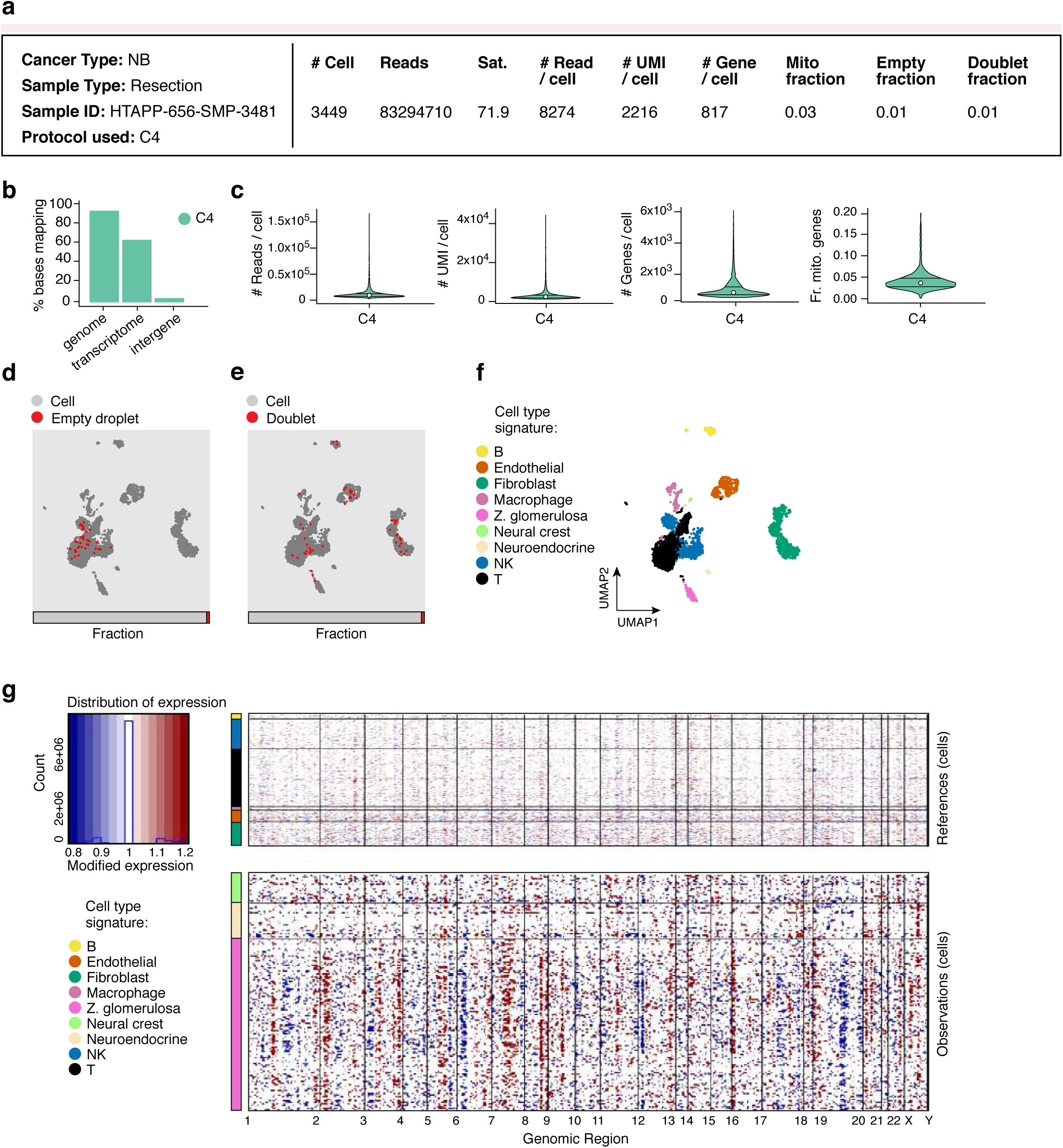
Evaluation of scRNA-Seq protocol for neuroblastoma resection. **(A)** Sample processing and QC overview. Shown are the number of cells passing QC, and the number of sequencing reads and sequencing saturation across all cells. The remaining metrics are reported for those cells passing QC: median number of reads per cell, median number of UMIs per cell, median number of genes per cell, median fraction of UMIs mapping to mitochondrial genes in each cell, fraction of cell barcodes called as empty droplets, and fraction of cell barcodes called as doublets. **(B)** Read mapping QCs. The percent of bases in the sequencing reads (*y* axis) mapping to the genome, transcriptome, and intergenic regions (*x* axis). **(C)** Overall QCs. Distribution (median and first and third quartiles) of the number of reads per cell, number of UMIs per cell, number of genes per cell, and fraction of UMIs mapping to mitochondrial genes in each cell (*y* axes) for all cells passing QC. **(D,E)** Relation of empty droplets and doublets to cell types. UMAP embedding and fraction (horizontal bar) of single cell (grey), “empty droplet” (red, left) and doublet (red, right) profiles. **(F)** Cell type assignment. UMAP embedding of single cell profiles colored by assigned cell type signature. **(G)** Inferred CNA profiles for cells. Chromosomal amplification (red) and deletion (blue) inferred in each chromosomal position (columns) across the single cells (rows). Top: reference cells not expected to contain CNA in this cancer type. Bottom: cells tested for CNA relative to the reference cells. Color bar: assigned cell type signature for each cell.

**Supplementary Figure 14.**
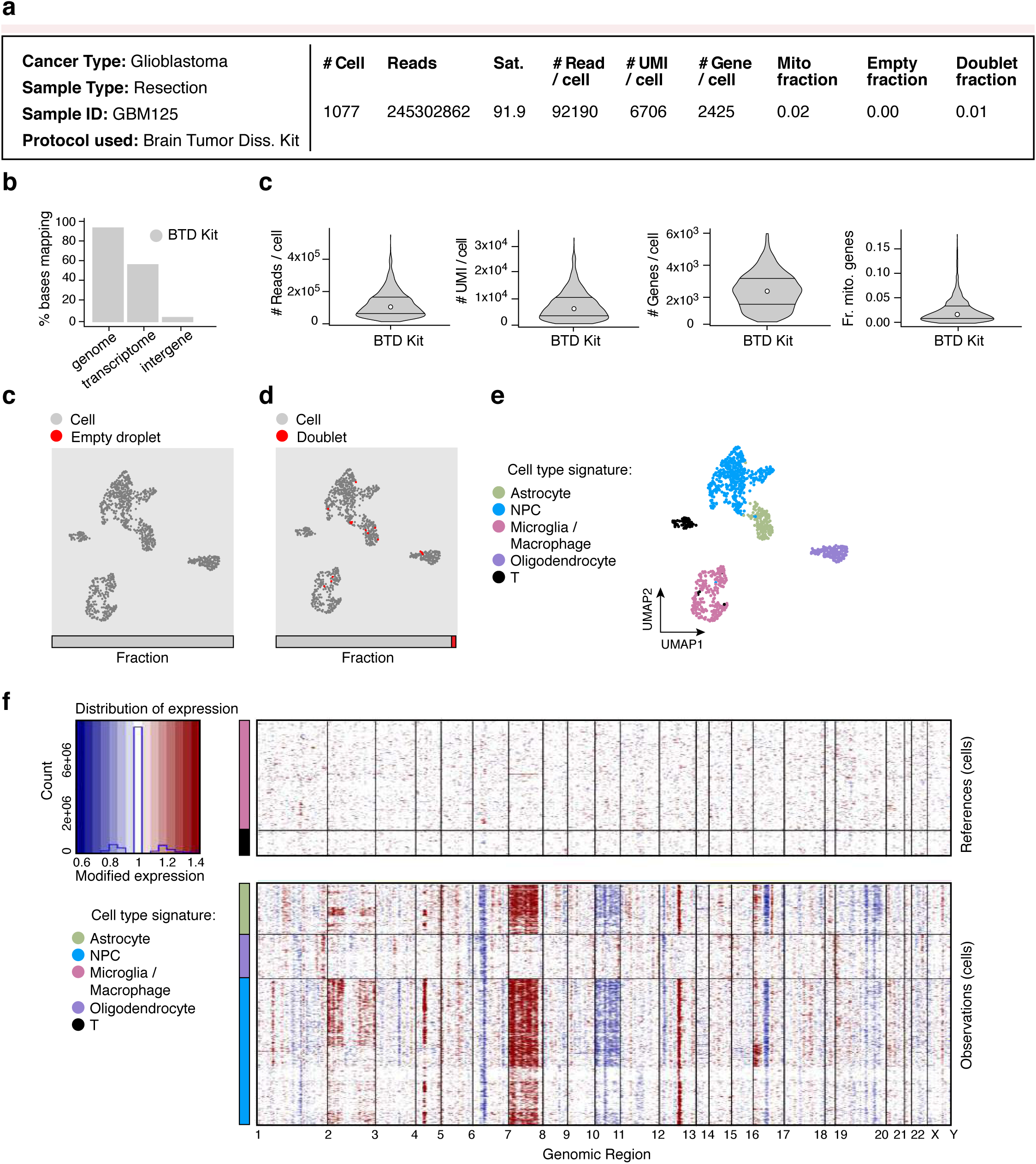
Evaluation of scRNA-Seq protocol for glioma resection. **(A)** Sample processing and QC overview. Shown are the number of cells passing QC, and the number of sequencing reads and sequencing saturation across all cells. The remaining metrics are reported for those cells passing QC: median number of reads per cell, median number of UMIs per cell, median number of genes per cell, median fraction of UMIs mapping to mitochondrial genes in each cell, fraction of cell barcodes called as empty droplets, and fraction of cell barcodes called as doublets. **(B)** Read mapping QCs. The percent of bases in the sequencing reads (*y* axis) mapping to the genome, transcriptome, and intergenic regions (*x* axis). **(C)** Overall QCs. Distribution (median and first and third quartiles) of the number of reads per cell, number of UMIs per cell, number of genes per cell, and fraction of UMIs mapping to mitochondrial genes in each cell (*y* axes) for all cells passing QC. **(D,E)** Relation of empty droplets and doublets to cell types. UMAP embedding and fraction (horizontal bar) of single cell (grey), “empty droplet” (red, left) and doublet (red, right) profiles. **(F)** Cell type assignment. UMAP embedding of single cell profiles colored by assigned cell type signature. **(G)** Inferred CNA profiles for cells. Chromosomal amplification (red) and deletion (blue) inferred in each chromosomal position (columns) across the single cells (rows). Top: reference cells not expected to contain CNA in this cancer type. Bottom: cells tested for CNA relative to the reference cells. Color bar: assigned cell type signature for each cell.

**Supplementary Figure 15.**
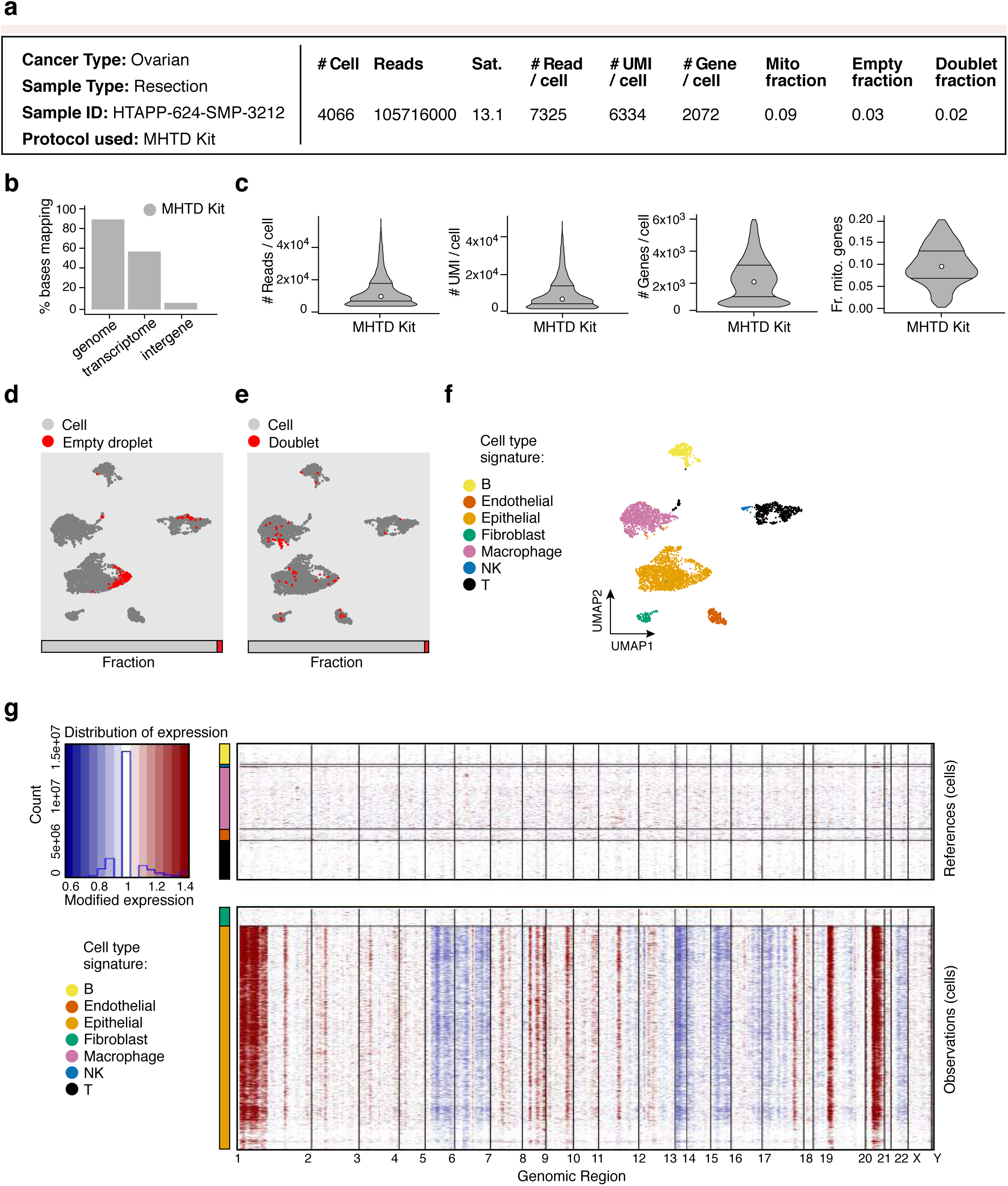
Evaluation of scRNA-Seq protocol for ovarian cancer resection. **(A)** Sample processing and QC overview. Shown are the number of cells passing QC, and the number of sequencing reads and sequencing saturation across all cells. The remaining metrics are reported for those cells passing QC: median number of reads per cell, median number of UMIs per cell, median number of genes per cell, median fraction of UMIs mapping to mitochondrial genes in each cell, fraction of cell barcodes called as empty droplets, and fraction of cell barcodes called as doublets. **(B)** Read mapping QCs. The percent of bases in the sequencing reads (*y* axis) mapping to the genome, transcriptome, and intergenic regions (*x* axis). **(C)** Overall QCs. Distribution (median and first and third quartiles) of the number of reads per cell, number of UMIs per cell, number of genes per cell, and fraction of UMIs mapping to mitochondrial genes in each cell (*y* axes) for all cells passing QC. **(D,E)** Relation of empty droplets and doublets to cell types. UMAP embedding and fraction (horizontal bar) of single cell (grey), “empty droplet” (red, left) and doublet (red, right) profiles. **(F)** Cell type assignment. UMAP embedding of single cell profiles colored by assigned cell type signature. **(G)** Inferred CNA profiles for cells. Chromosomal amplification (red) and deletion (blue) inferred in each chromosomal position (columns) across the single cells (rows). Top: reference cells not expected to contain CNA in this cancer type. Bottom: cells tested for CNA relative to the reference cells. Color bar: assigned cell type signature for each cell.

**Supplementary Figure 16.**
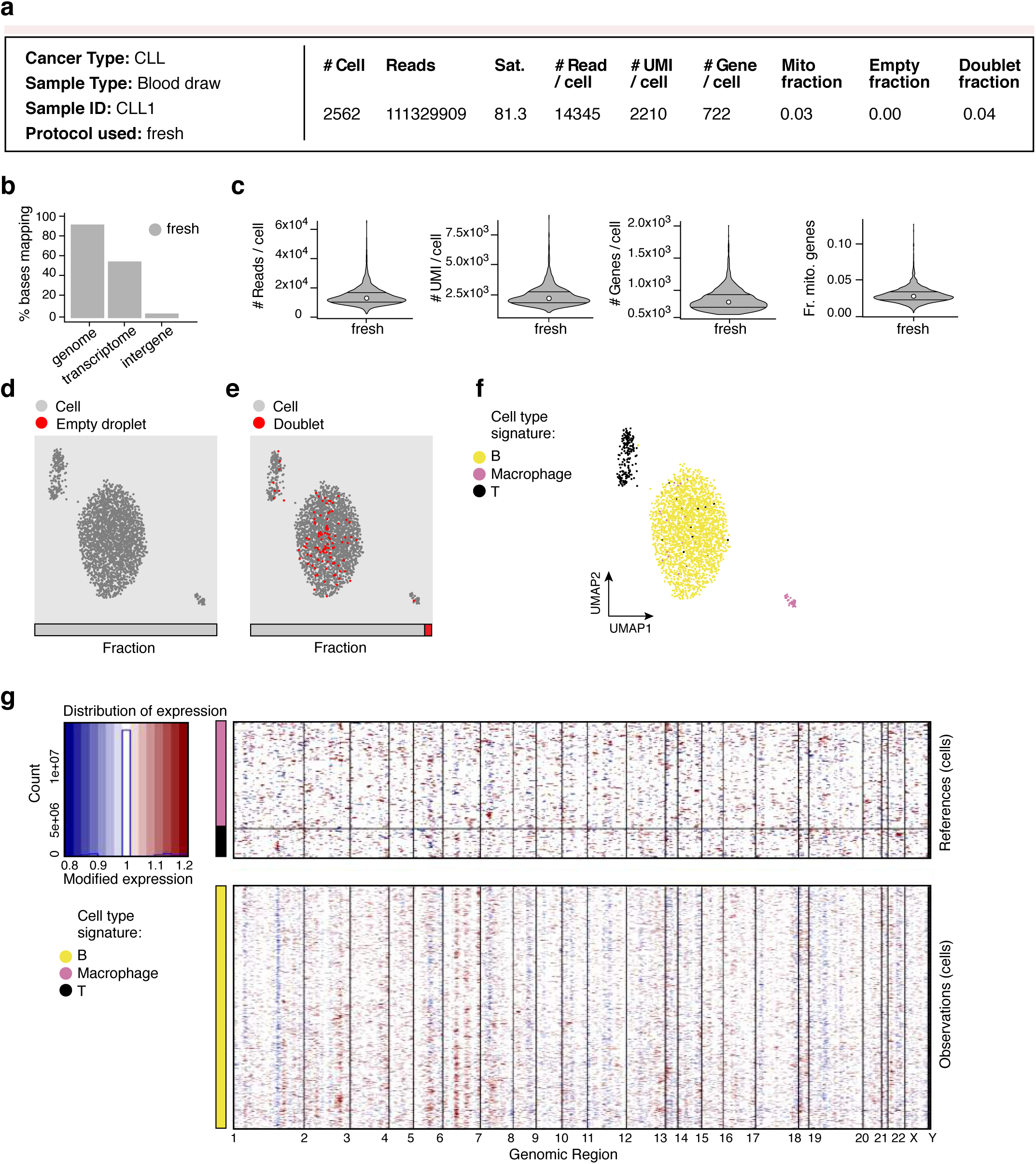
Evaluation of scRNA-Seq protocol for a cryopreserved CLL sample. **(A)** Sample processing and QC overview. Shown are the number of cells passing QC, and the number of sequencing reads and sequencing saturation across all cells. The remaining metrics are reported for those cells passing QC: median number of reads per cell, median number of UMIs per cell, median number of genes per cell, median fraction of UMIs mapping to mitochondrial genes in each cell, fraction of cell barcodes called as empty droplets, and fraction of cell barcodes called as doublets. **(B)** Read mapping QCs. The percent of bases in the sequencing reads (*y* axis) mapping to the genome, transcriptome, and intergenic regions (*x* axis). **(C)** Overall QCs. Distribution (median and first and third quartiles) of the number of reads per cell, number of UMIs per cell, number of genes per cell, and fraction of UMIs mapping to mitochondrial genes in each cell (*y* axes) for all cells passing QC. **(D,E)** Relation of empty droplets and doublets to cell types. UMAP embedding and fraction (horizontal bar) of single cell (grey), “empty droplet” (red, left) and doublet (red, right) profiles. **(F)** Cell type assignment. UMAP embedding of single cell profiles colored by assigned cell type signature. Note that the cell type signature used for macrophages contains macrophage and monocyte markers. **(G)** Inferred CNA profiles for cells. Chromosomal amplification (red) and deletion (blue) inferred in each chromosomal position (columns) across the single cells (rows). Top: reference cells not expected to contain CNA in this cancer type. Bottom: cells tested for CNA relative to the reference cells. Color bar: assigned cell type signature for each cell.

**Supplementary Figure 17.**
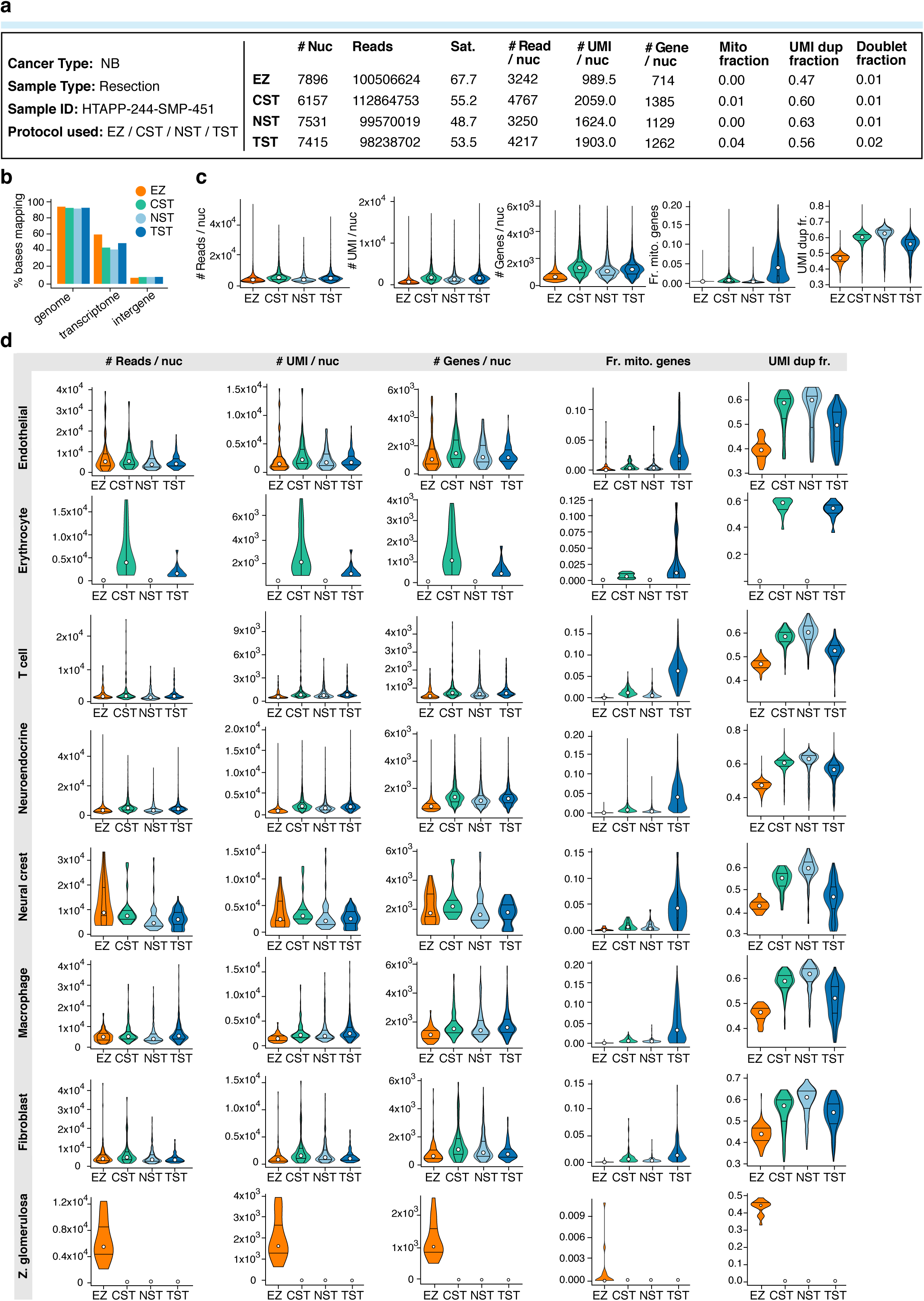
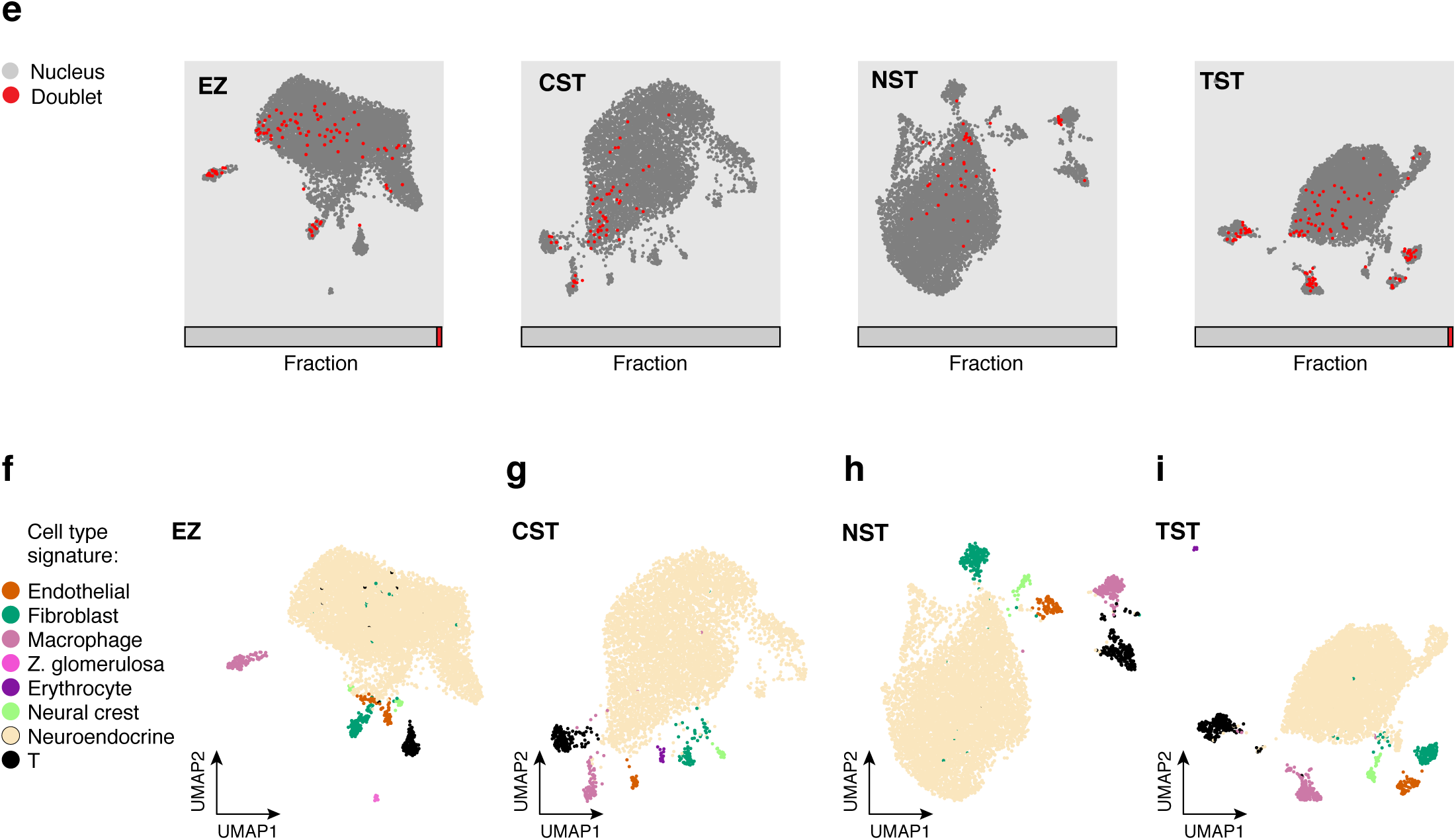
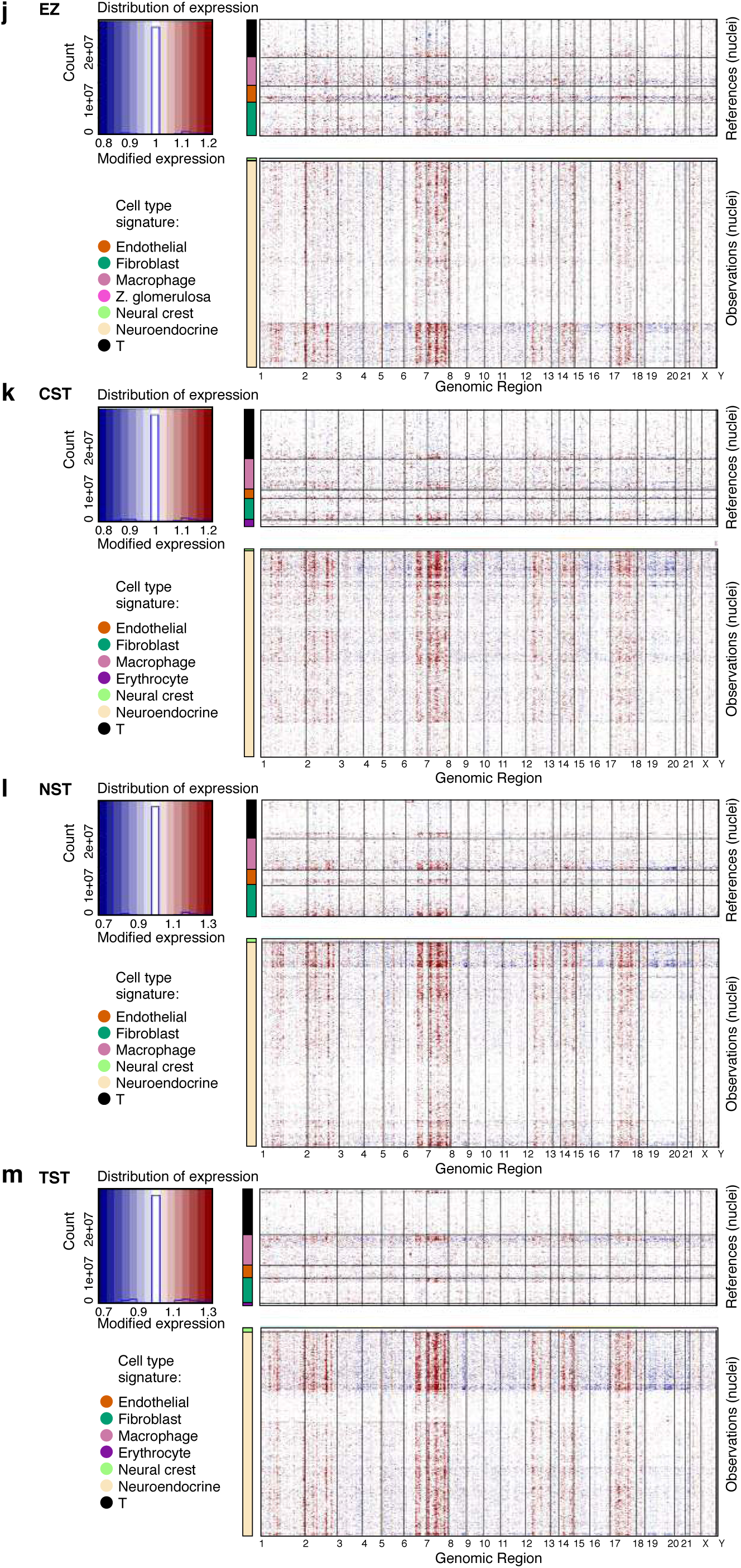
SnRNA-Seq protocol comparison for a single neuroblastoma sample. **(A)** Sample processing and QC overview. For each protocol, shown are the number of nuclei passing QC, and the number of sequencing reads and sequencing saturation across all nuclei. The remaining metrics are reported for those nuclei passing QC: the median number of reads per nucleus, median number of UMIs per nucleus, median number of genes per nucleus, median fraction of UMIs mapping to mitochondrial genes in each nucleus, median fraction of duplicated UMIs per nucleus, and fraction of nucleus barcodes called as doublets. **(B)** Read mapping QCs. The percent of bases in the sequencing reads (*y* axis) mapping to the genome, transcriptome, and intergenic regions (*x* axis) across the four protocols (colored bars). **(C-D)** Overall and cell types specific QCs. Distribution (median and first and third quartiles) of the number of reads per nucleus, number of UMIs per nucleus, number of genes per nucleus, fraction of UMIs mapping to mitochondrial genes in each nucleus, and fraction of duplicated UMIs per nucleus (*y* axes) in each of the four protocols (*x* axis), for all nuclei passing QC (**C**) and for nuclei from each cell type (**D**, rows). **(e)** Relation of doublets to cell types. UMAP embedding and fraction (horizontal bar) of single nucleus (grey) and doublet (red) profiles for each protocol. **(F-I)** Cell type assignment. UMAP embedding of single nucleus profiles from each protocol colored by assigned cell type signature. **(J-M)** Inferred CNA profiles. Chromosomal amplification (red) and deletion (blue) inferred in each chromosomal position (columns) across the single nuclei (rows). Top: reference nuclei not expected to contain CNA in this cancer type. Bottom: nuclei tested for CNA relative to the reference nuclei. Color bar: assigned cell type signature for each nucleus.

**Supplementary Figure 18.**
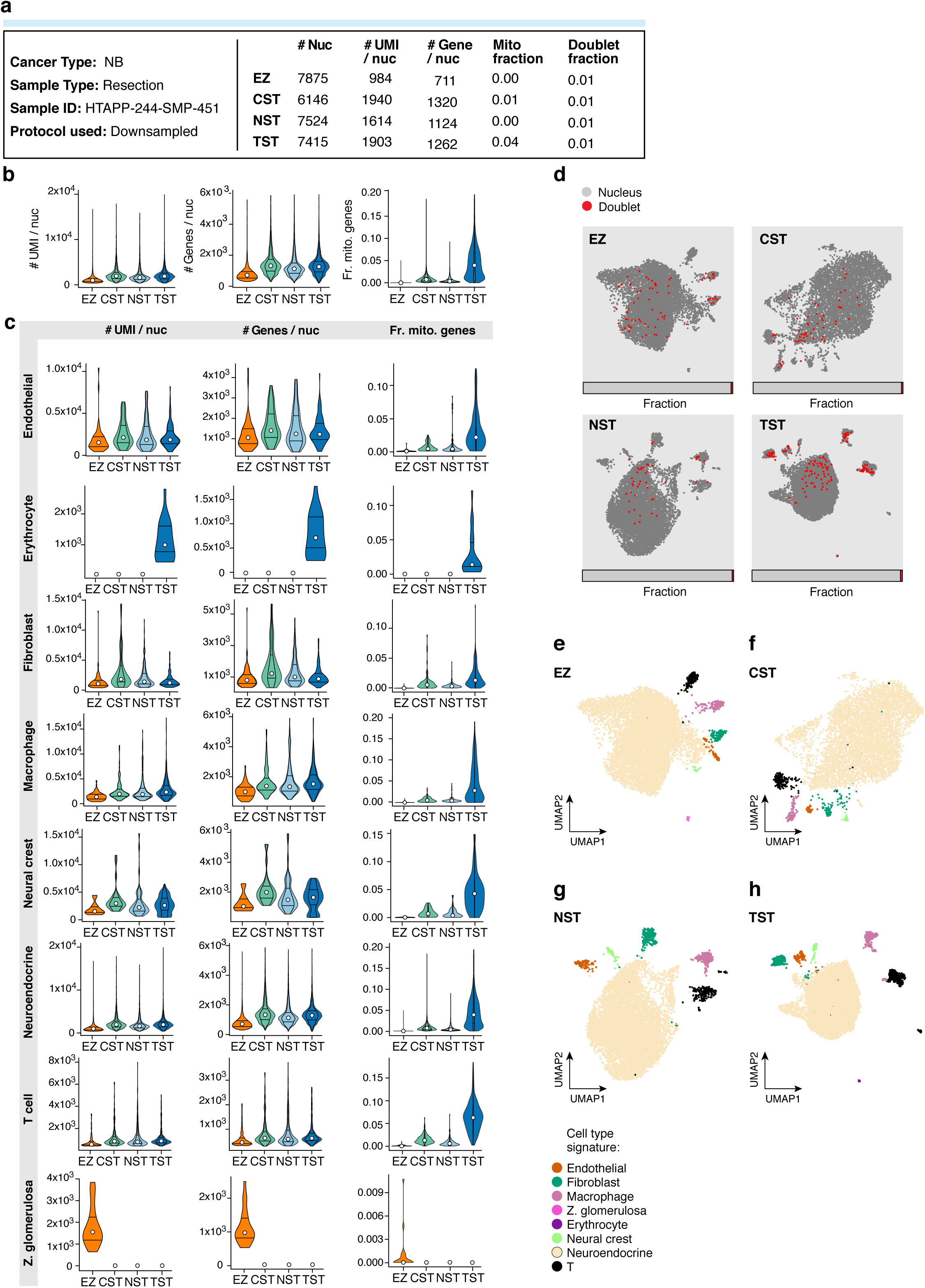
SnRNA-Seq protocol comparison for neuroblastoma following read down-sampling. Shown are analyses for NB HTAPP-244-SMP-451 (as in **Supp.** Fig. 17), but after the total number of sequencing reads within each sample was down-sampled to match the protocol with the fewest total sequencing reads. **(A)** Sample processing and QC overview. For each protocol, shown are the number of nuclei passing QC. The remaining metrics are reported for those nuclei passing QC: median number of UMIs per nucleus, median number of genes per nucleus, median fraction of UMIs mapping to mitochondrial genes in each nucleus, and fraction of nucleus barcodes called as doublets. **(B,C)** Overall and cell types specific QCs. Distribution (median and first and third quartiles) of the number of UMIs per nucleus, number of genes per nucleus, and fraction of UMIs mapping to mitochondrial genes in each nucleus (*y* axes) in each of the four protocols (*x* axis), for all nuclei passing QC **(B)** and for nuclei from each cell type (**C,** rows). **(D)** Relation of doublets to cell types. UMAP embedding and fraction (horizontal bar) of single nucleus (grey) and doublet (red) profiles for each protocol. **(E-H)** Cell type assignment. UMAP embedding of single nucleus profiles from each protocol colored by assigned cell type signature.

**Supplementary Figure 19.**
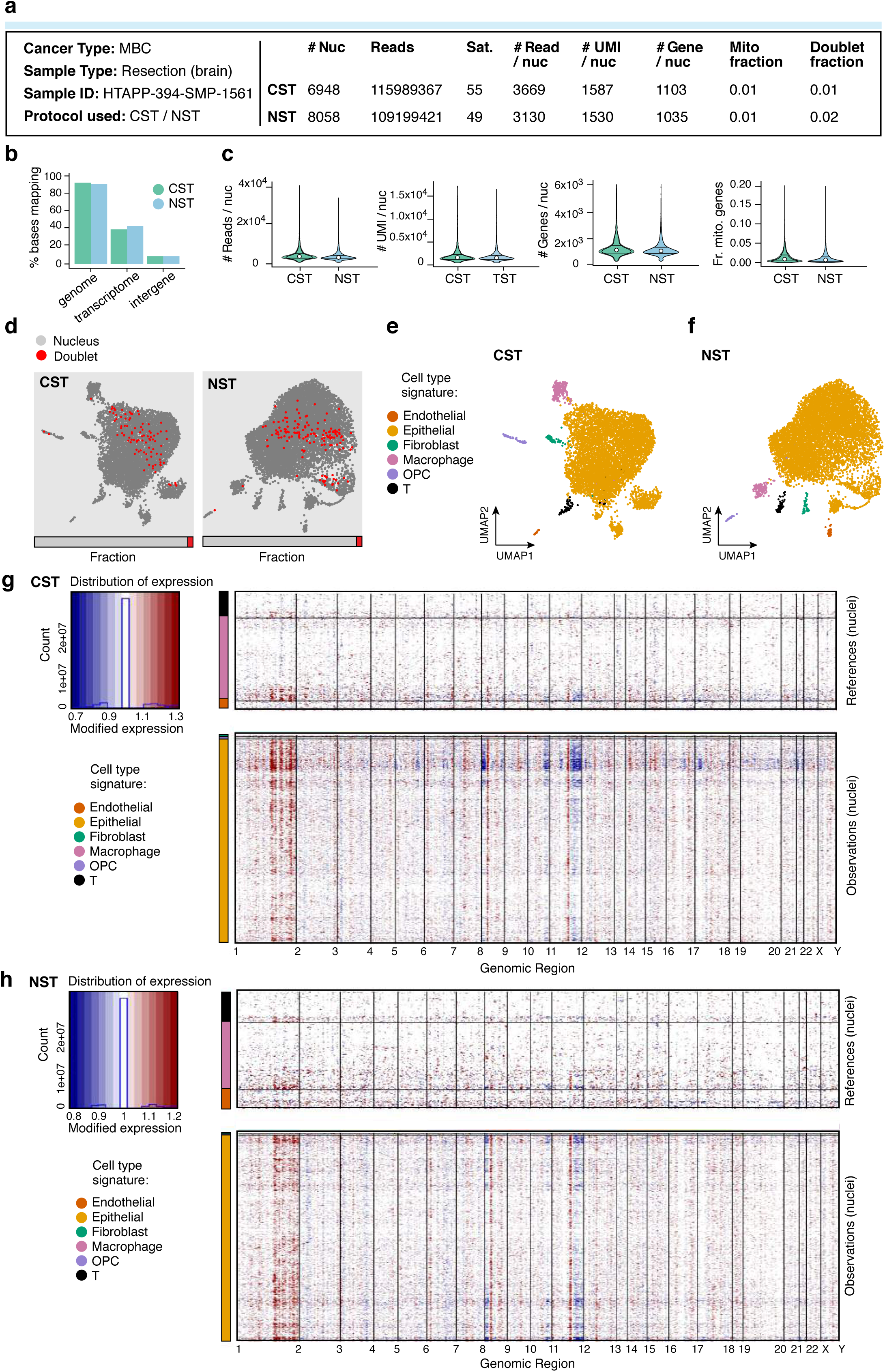
SnRNA-seq protocol comparison for a resection of a breast cancer metastasis from the brain. **(A)** Sample processing and QC overview. For each protocol, shown are the number of nuclei passing QC, and the number of sequencing reads and sequencing saturation across all nuclei. The remaining metrics are reported for those nuclei passing QC: median number of reads per nucleus, median number of UMIs per nucleus, median number of genes per nucleus, median fraction of UMIs mapping to mitochondrial genes in each nucleus, and fraction of nucleus barcodes called as doublets. **(B)** Read mapping QCs. The percent of bases in the sequencing reads (*y* axis) mapping to the genome, transcriptome, and intergenic regions (*x* axis). **(C)** Overall QCs. Distribution (median and first and third quartiles) of the number of reads per nucleus, number of UMIs per nucleus, number of genes per nucleus, and fraction of UMIs mapping to mitochondrial genes in each nucleus (*y* axes) for all nuclei passing QC. **(D)** Relation of doublets to cell types. UMAP embedding and fraction (horizontal bar) of single nucleus (grey) and doublet (red) profiles for each protocol. **(E-F)** Cell type assignment. UMAP embedding of single nucleus profiles from each protocol colored by assigned cell type signature. **(G-H)** Inferred CNA profiles for nuclei. Chromosomal amplification (red) and deletion (blue) inferred in each chromosomal position (columns) across the single nuclei (rows). Top: reference nuclei not expected to contain CNA in this cancer type. Bottom: nuclei tested for CNA relative to the reference nuclei. Color bar: assigned cell type signature for each nucleus.

**Supplementary Figure 20.**
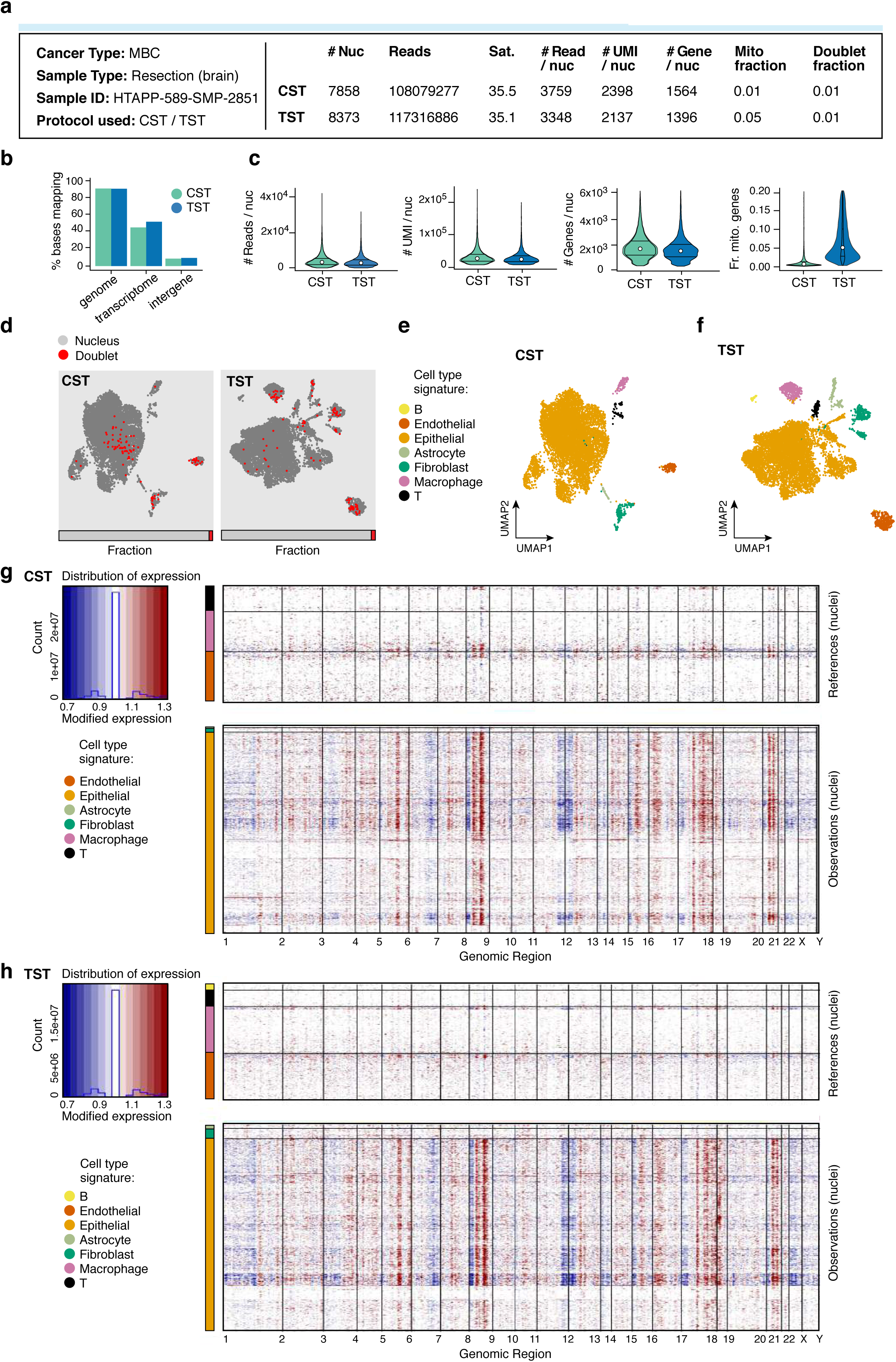
SnRNA-seq protocol comparison for another resection of a breast cancer metastasis from the brain. **(A)** Sample processing and QC overview. For each protocol, shown are the number of nuclei passing QC, and the number of sequencing reads and sequencing saturation across all nuclei. The remaining metrics are reported for those nuclei passing QC: median number of reads per nucleus, median number of UMIs per nucleus, median number of genes per nucleus, median fraction of UMIs mapping to mitochondrial genes in each nucleus, and fraction of nucleus barcodes called as doublets. **(B)** Read mapping QCs. The percent of bases in the sequencing reads (*y* axis) mapping to the genome, transcriptome, and intergenic regions (*x* axis). **(C)** Overall QCs. Distribution (median and first and third quartiles) of the number of reads per nucleus, number of UMIs per nucleus, number of genes per nucleus, and fraction of UMIs mapping to mitochondrial genes in each nucleus (*y* axes) for all nuclei passing QC. **(D)** Relation of doublets to cell types. UMAP embedding and fraction (horizontal bar) of single nucleus (grey) and doublet (red) profiles for each protocol. **(E-F)** Cell type assignment. UMAP embedding of single nucleus profiles from each protocol colored by assigned cell type signature. **(G-H)** Inferred CNA profiles for nuclei. Chromosomal amplification (red) and deletion (blue) inferred in each chromosomal position (columns) across the single nuclei (rows). Top: reference nuclei not expected to contain CNA in this cancer type. Bottom: nuclei tested for CNA relative to the reference nuclei. Color bar: assigned cell type signature for each nucleus.

**Supplementary Figure 21.**
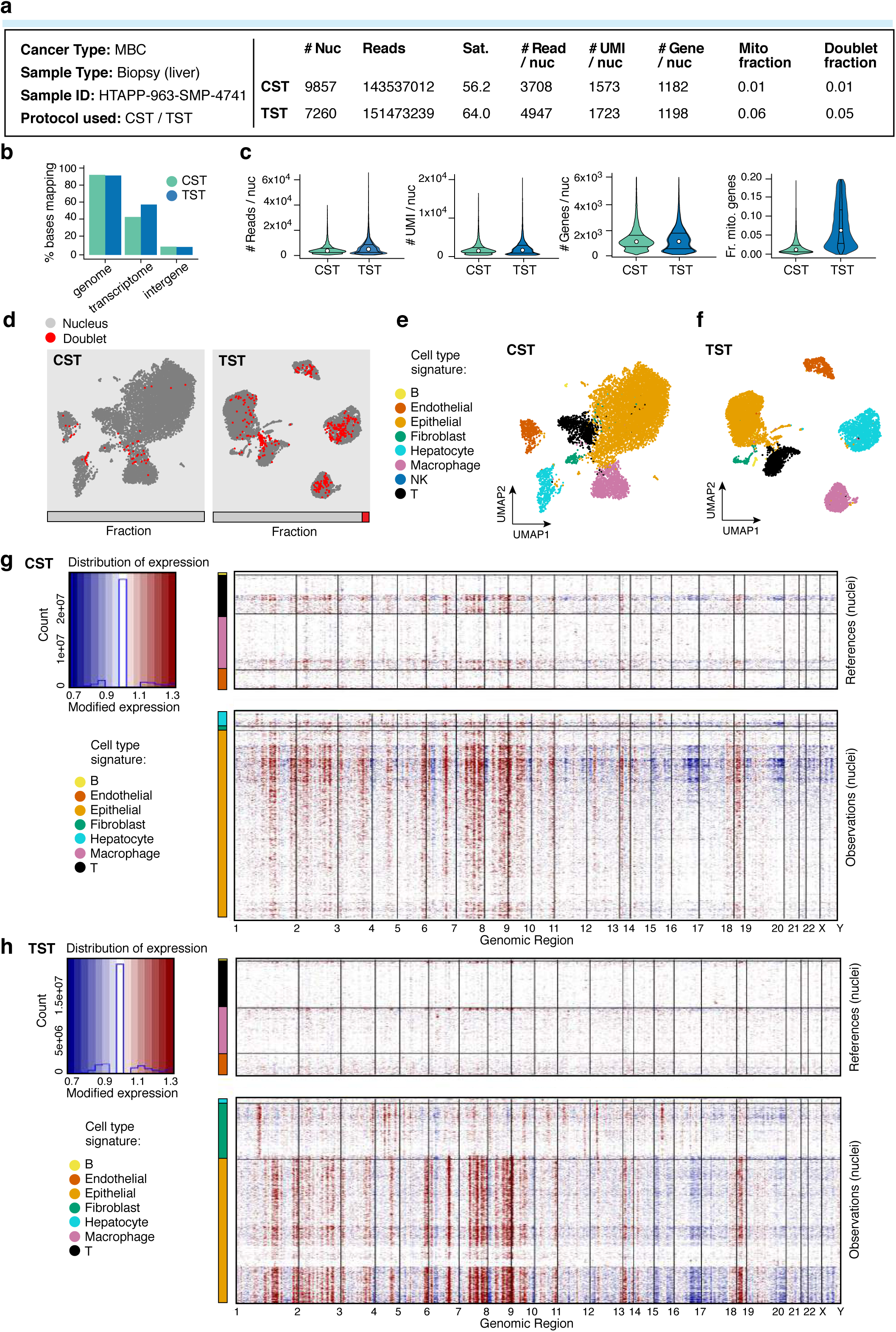
SnRNA-seq protocol comparison for a liver biopsy of metastatic breast cancer. **(A)** Sample processing and QC overview. For each protocol, shown are the number of nuclei passing QC, and the number of sequencing reads and sequencing saturation across all nuclei. The remaining metrics are reported for those nuclei passing QC: median number of reads per nucleus, median number of UMIs per nucleus, median number of genes per nucleus, median fraction of UMIs mapping to mitochondrial genes in each nucleus, and fraction of nucleus barcodes called as doublets. **(B)** Read mapping QCs. The percent of bases in the sequencing reads (*y* axis) mapping to the genome, transcriptome, and intergenic regions (*x* axis). **(C)** Overall QCs. Distribution (median and first and third quartiles) of the number of reads per nucleus, number of UMIs per nucleus, number of genes per nucleus, and fraction of UMIs mapping to mitochondrial genes in each nucleus (*y* axes) for all nuclei passing QC. **(D)** Relation of doublets to cell types. UMAP embedding and fraction (horizontal bar) of single nucleus (grey) and doublet (red) profiles for each protocol. **(E-F)** Cell type assignment. UMAP embedding of single nucleus profiles from each protocol colored by assigned cell type signature. **(G-H)** Inferred CNA profiles for nuclei. Chromosomal amplification (red) and deletion (blue) inferred in each chromosomal position (columns) across the single nuclei (rows). Top: reference nuclei not expected to contain CNA in this cancer type. Bottom: nuclei tested for CNA relative to the reference nuclei. Color bar: assigned cell type signature for each nucleus.

**Supplementary Figure 22.**
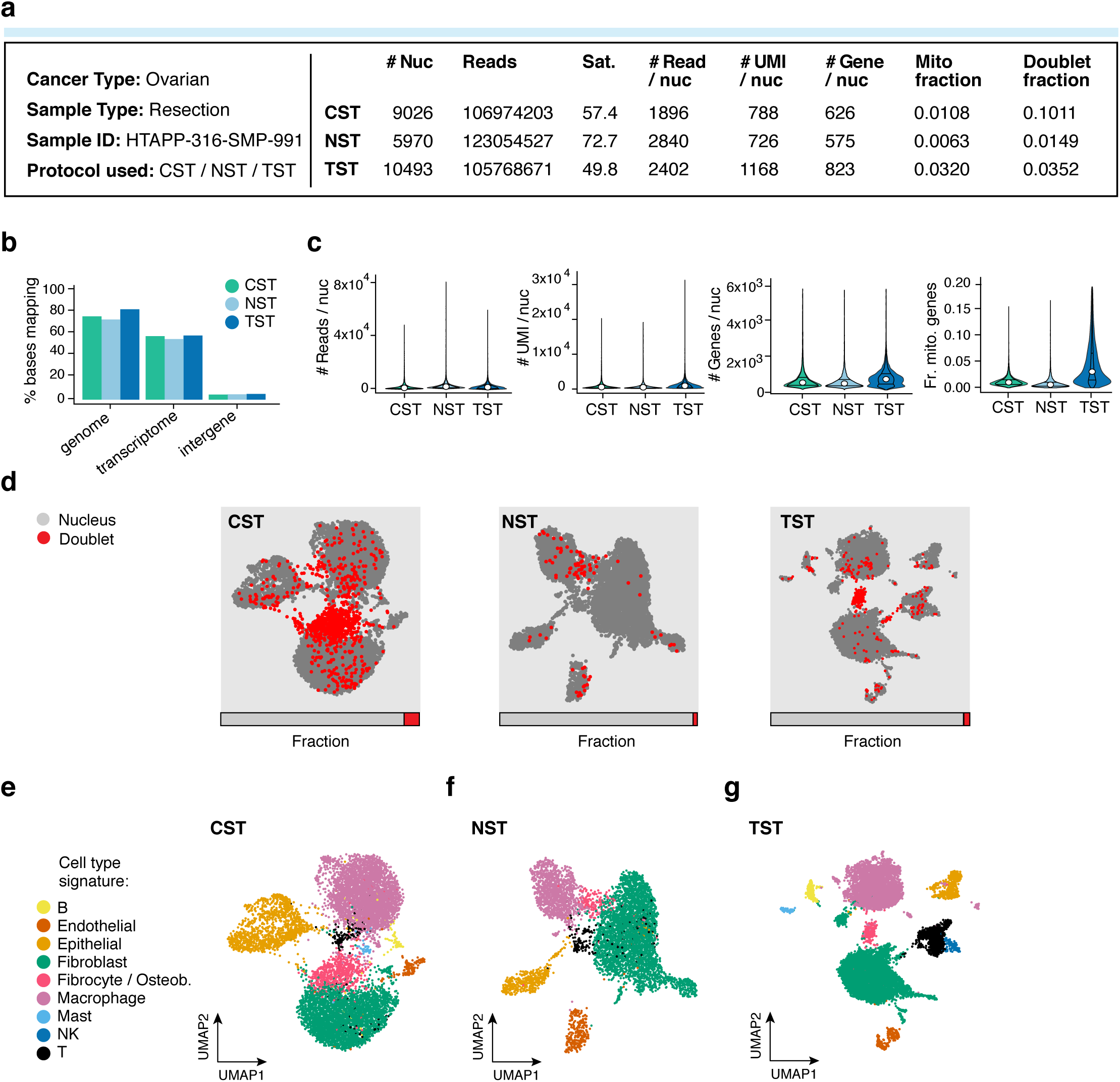
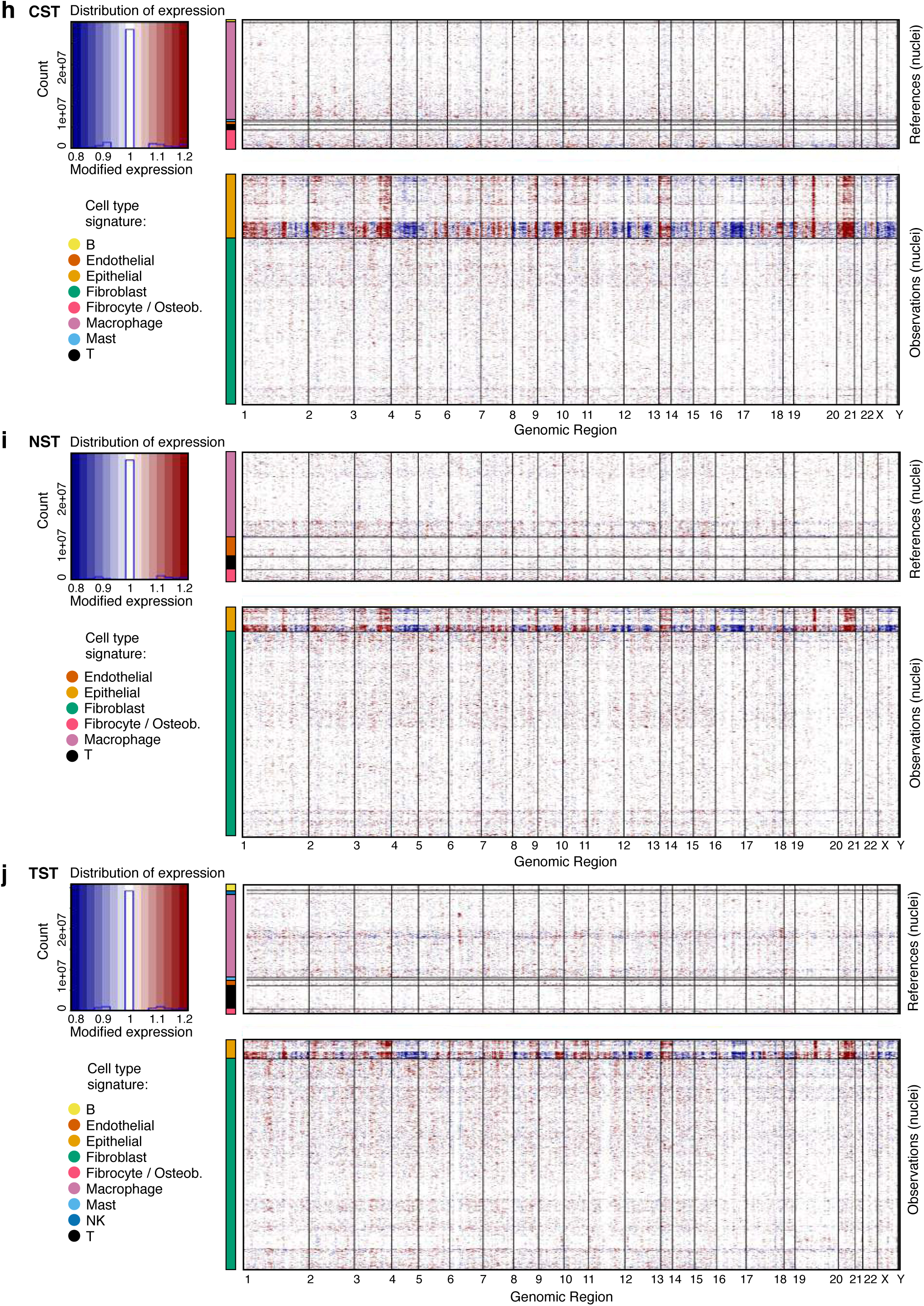
SnRNA-Seq protocol comparison for of ovarian cancer resection. **(A)** Sample processing and QC overview. For each protocol, shown are the number of nuclei passing QC, and the number of sequencing reads and sequencing saturation across all nuclei. The remaining metrics are reported for those nuclei passing QC: median number of reads per nucleus, median number of UMIs per nucleus, median number of genes per nucleus, median fraction of UMIs mapping to mitochondrial genes in each nucleus, and fraction of nucleus barcodes called as doublets. **(B)** Read mapping QCs. The percent of bases in the sequencing reads (*y* axis) mapping to the genome, transcriptome, and intergenic regions (*x* axis). **(C)** Overall QCs. Distribution (median and first and third quartiles) of the number of reads per nucleus, number of UMIs per nucleus, number of genes per nucleus, and fraction of UMIs mapping to mitochondrial genes in each nucleus (*y* axes) for all nuclei passing QC. **(D)** Relation of doublets to cell types. UMAP embedding and fraction (horizontal bar) of single nucleus (grey) and doublet (red) profiles for each protocol. **(E-G)** Cell type assignment. UMAP embedding of single nucleus profiles from each protocol colored by assigned cell type signature. **(H-J)** Inferred CNA profiles for nuclei. Chromosomal amplification (red) and deletion (blue) inferred in each chromosomal position (columns) across the single nuclei (rows). Top: reference nuclei not expected to contain CNA in this cancer type. Bottom: nuclei tested for CNA relative to the reference nuclei. Color bar: assigned cell type signature for each nucleus.

**Supplementary Figure 23.**
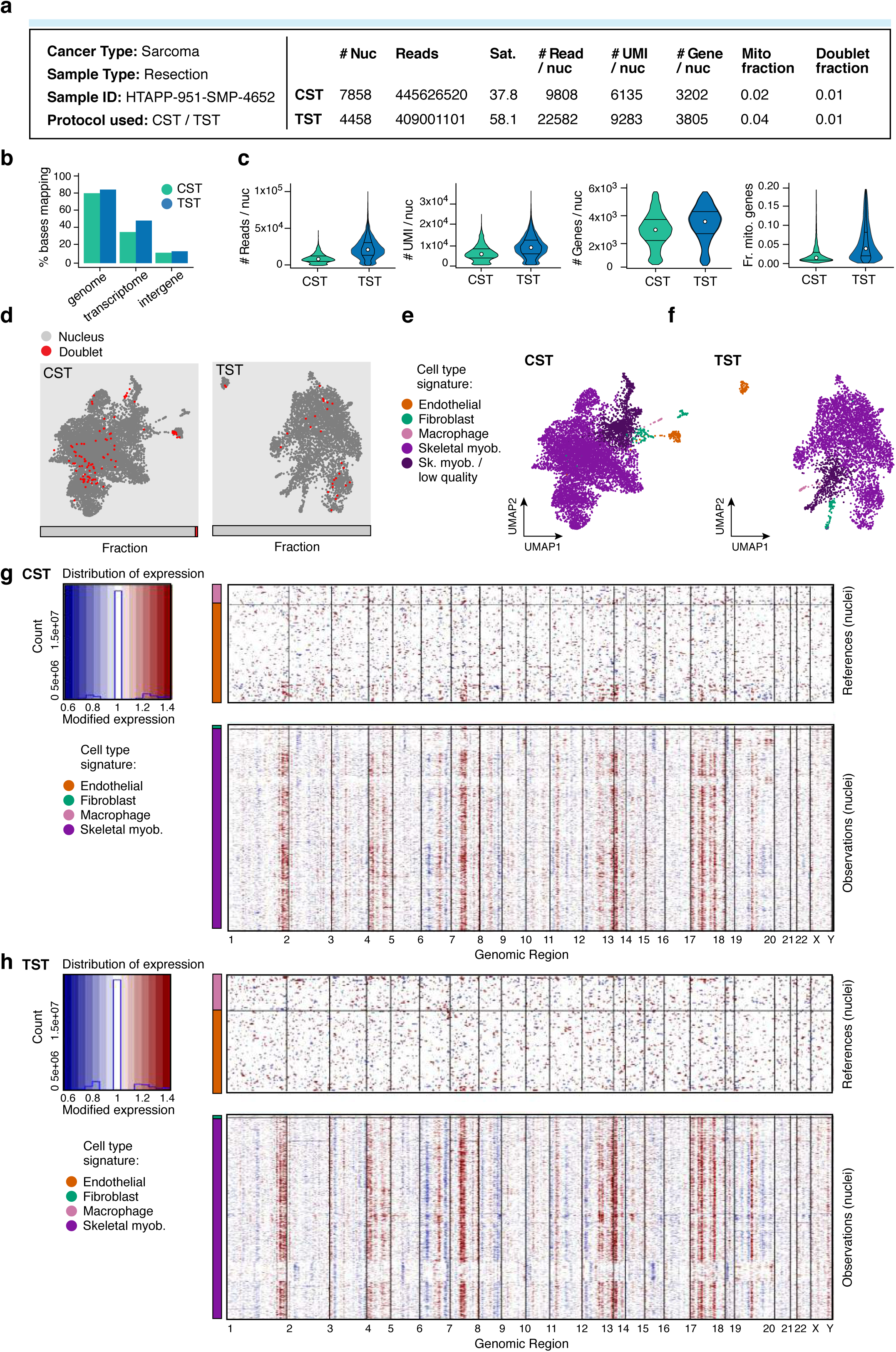
SnRNA-Seq protocol comparison for a sarcoma resection. **(A)** Sample processing and QC overview. For each protocol, shown are the number of nuclei passing QC, and the number of sequencing reads and sequencing saturation across all nuclei. The remaining metrics are reported for those nuclei passing QC: median number of reads per nucleus, median number of UMIs per nucleus, median number of genes per nucleus, median fraction of UMIs mapping to mitochondrial genes, and fraction of nucleus barcodes called as doublets. **(B)** Read mapping QCs. The percent of bases in the sequencing reads (*y* axis) mapping to the genome, transcriptome, and intergenic regions (*x* axis). **(C)** Overall QCs. Distribution (median and first and third quartiles) of the number of reads per nucleus, number of UMIs per nucleus, number of genes per nucleus, and fraction of UMIs mapping to mitochondrial genes in each nucleus (*y* axes) for all nuclei passing QC. **(D)** Relation of doublets to cell types. UMAP embedding and fraction (horizontal bar) of single nucleus (grey) and doublet (red) profiles for each protocol. **(E-F)** Cell type assignment. UMAP embedding of single nucleus profiles from each protocol colored by assigned cell type signature. **(G-H)** Inferred CNA profiles for nuclei. Chromosomal amplification (red) and deletion (blue) inferred in each chromosomal position (columns) across the single nuclei (rows). Top: reference nuclei not expected to contain CNA in this cancer type. Bottom: nuclei tested for CNA relative to the reference nuclei. Color bar: assigned cell type signature for each nucleus.

**Supplementary Figure 24.**
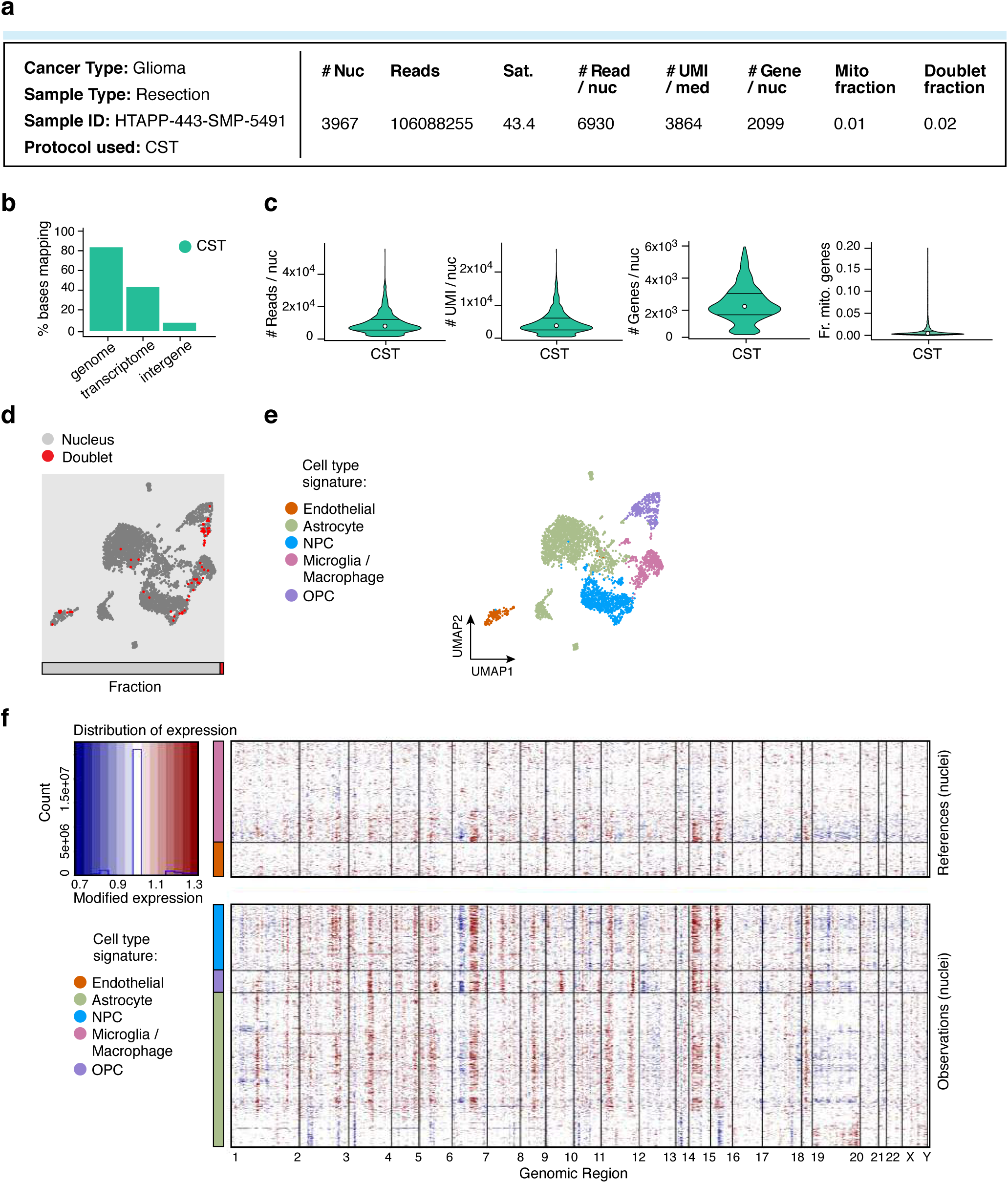
Evaluation of snRNA-Seq protocol for a glioma resection. **(A)** Sample processing and QC overview. Shown are the number of nuclei passing QC, and the number of sequencing reads and sequencing saturation across all nuclei. The remaining metrics are reported for those nuclei passing QC: median number of reads per nucleus, median number of UMIs per nucleus, median number of genes per nucleus, median fraction of UMIs mapping to mitochondrial genes in each nucleus, and fraction of nucleus barcodes called as doublets. **(B)** Read mapping QCs. The percent of bases in the sequencing reads (*y* axis) mapping to the genome, transcriptome, and intergenic regions (*x* axis). **(C)** Overall QCs. Distribution (median and first and third quartiles) of the number of reads per nucleus, number of UMIs per nucleus, number of genes per nucleus, and fraction of UMIs mapping to mitochondrial genes in each nucleus (*y* axes) for all nuclei passing QC. **(D)** Relation of doublets to cell types. UMAP embedding and fraction (horizontal bar) of single nucleus (grey) and doublet (red) profiles. **(E)** Cell type assignment. UMAP embedding of single nucleus profiles colored by assigned cell type signature. **(F)** Inferred CNA profiles for nuclei. Chromosomal amplification (red) and deletion (blue) inferred in each chromosomal position (columns) across the single nuclei (rows). Top: reference nuclei not expected to contain CNA in this cancer type. Bottom: nuclei tested for CNA relative to the reference nuclei. Color bar: assigned cell type signature for each nucleus.

**Supplementary Figure 25.**
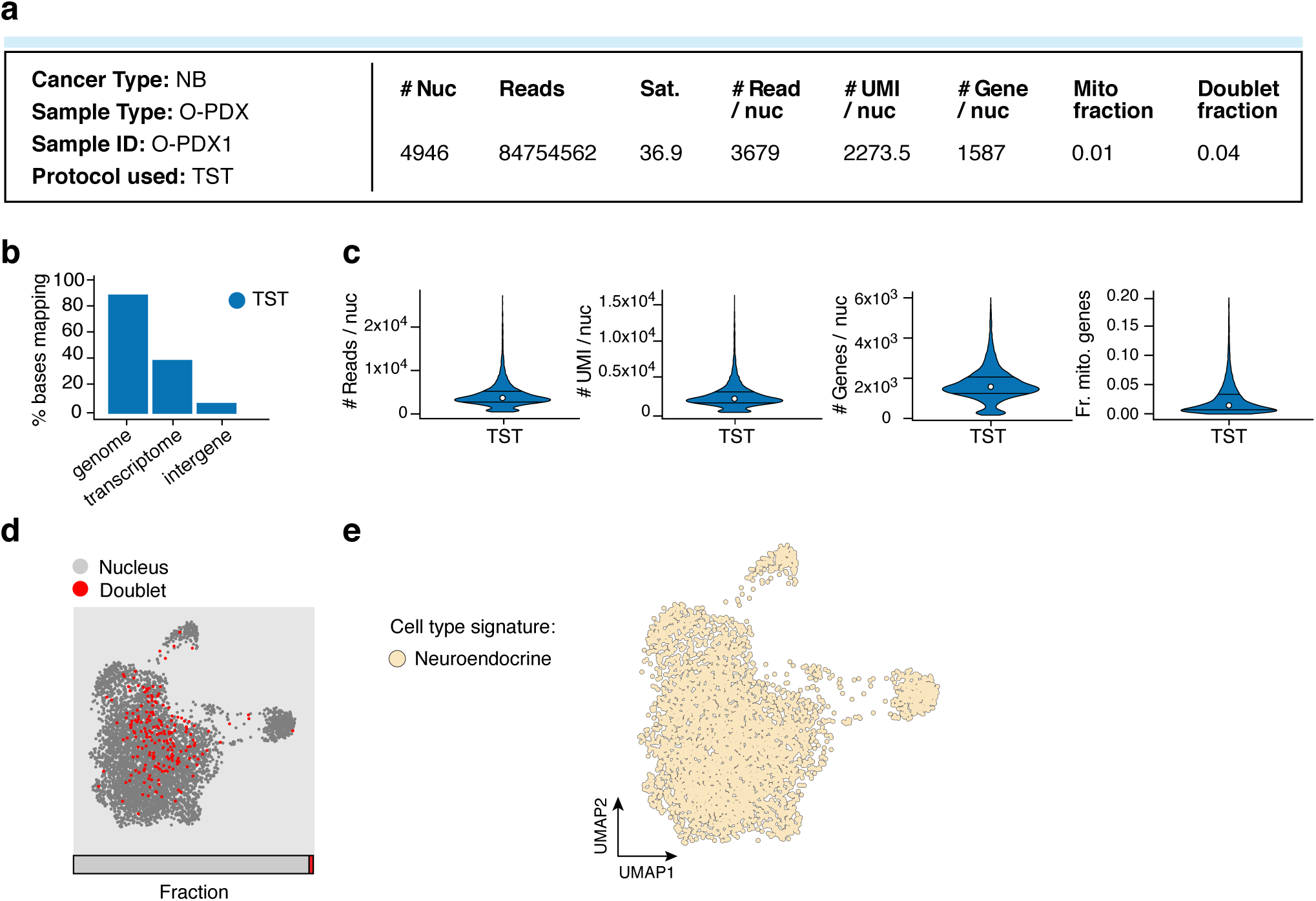
Evaluation of snRNA-Seq protocol for a neuroblastoma O-PDX. **(A)** Sample processing and QC overview. Shown are the number of nuclei passing QC, and the number of sequencing reads and sequencing saturation across all nuclei. The remaining metrics are reported for those nuclei passing QC: median number of reads per nucleus, median number of UMIs per nucleus, median number of genes per nucleus, median fraction of UMIs mapping to mitochondrial genes in each nucleus, and fraction of nucleus barcodes called as doublets. **(B)** Read mapping QCs. The percent of bases in the sequencing reads (*y* axis) mapping to the genome, transcriptome, and intergenic regions (*x* axis). **(C)** Overall QCs. Distribution (median and first and third quartiles) of the number of reads per nucleus, number of UMIs per nucleus, number of genes per nucleus, and fraction of UMIs mapping to mitochondrial genes in each nucleus (*y* axes) for all nuclei passing QC. **(D)** Relation of doublets to cell types. UMAP embedding and fraction (horizontal bar) of single nucleus (grey) and doublet (red) profiles. **(E)** Cell type assignment. UMAP embedding of single nucleus profiles colored by assigned cell type signature.

**Supplementary Figure 26.**
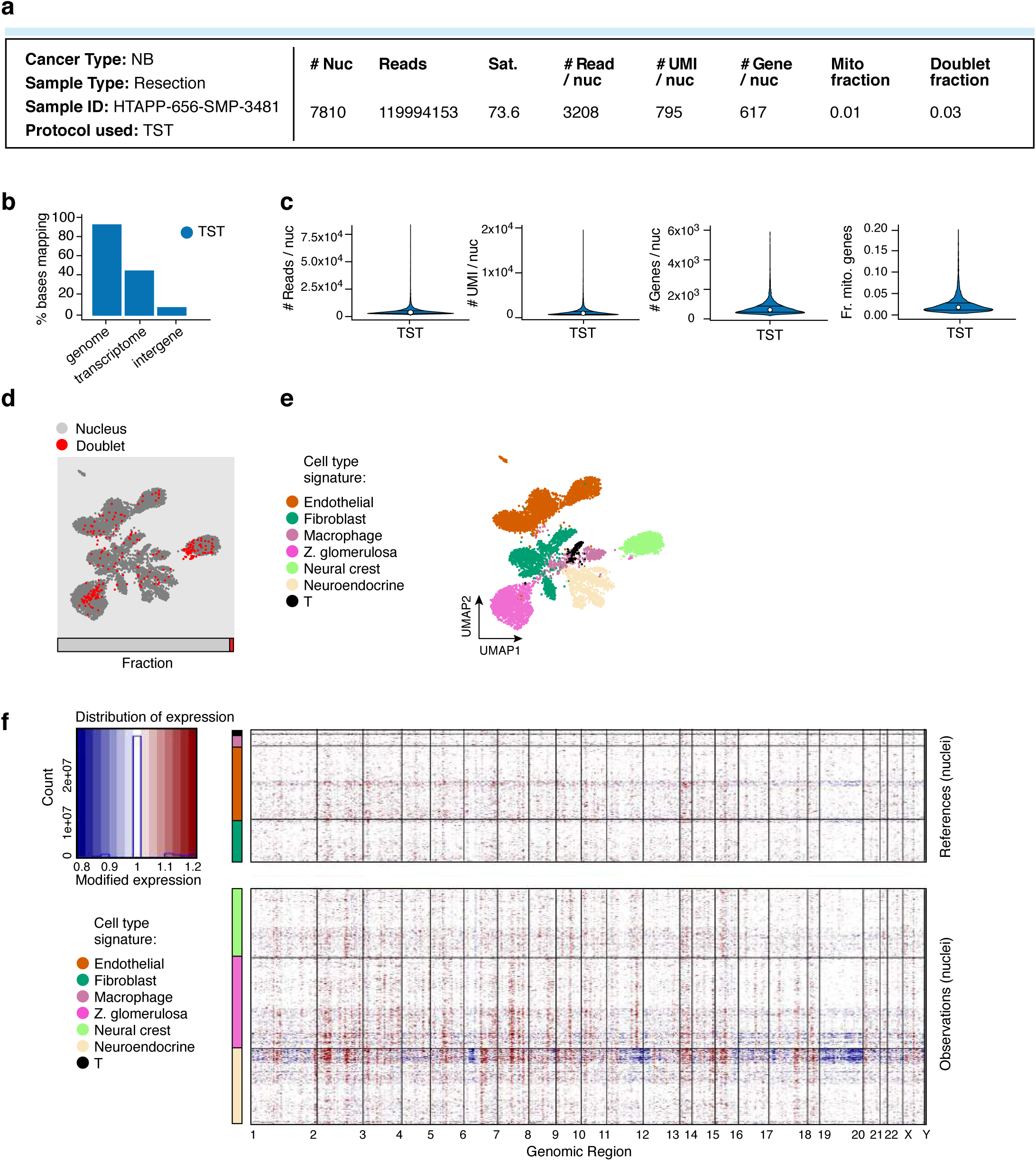
Evaluation of snRNA-Seq protocol for a neuroblastoma resection. **(A)** Sample processing and QC overview. Shown are the number of nuclei passing QC, and the number of sequencing reads and sequencing saturation across all nuclei. The remaining metrics are reported for those nuclei passing QC: median number of reads per nucleus, median number of UMIs per nucleus, median number of genes per nucleus, median fraction of UMIs mapping to mitochondrial genes in each nucleus, and fraction of nucleus barcodes called as doublets. **(B)** Read mapping QCs. The percent of bases in the sequencing reads (*y* axis) mapping to the genome, transcriptome, and intergenic regions (*x* axis). **(C)** Overall QCs. Distribution (median and first and third quartiles) of the number of reads per nucleus, number of UMIs per nucleus, number of genes per nucleus, and fraction of UMIs mapping to mitochondrial genes in each nucleus (*y* axes) for all nuclei passing QC. **(D)** Relation of doublets to cell types. UMAP embedding and fraction (horizontal bar) of single nucleus (grey) and doublet (red) profiles for each protocol. **(E)** Cell type assignment. UMAP embedding of single nucleus profiles from each protocol colored by assigned cell type signature. **(F)** Inferred CNA profiles for nuclei. Chromosomal amplification (red) and deletion (blue) inferred in each chromosomal position (columns) across the single nuclei (rows). Top: reference nuclei not expected to contain CNA in this cancer type. Bottom: nuclei tested for CNA relative to the reference nuclei. Color bar: assigned cell type signature for each nucleus.

**Supplementary Figure 27.**
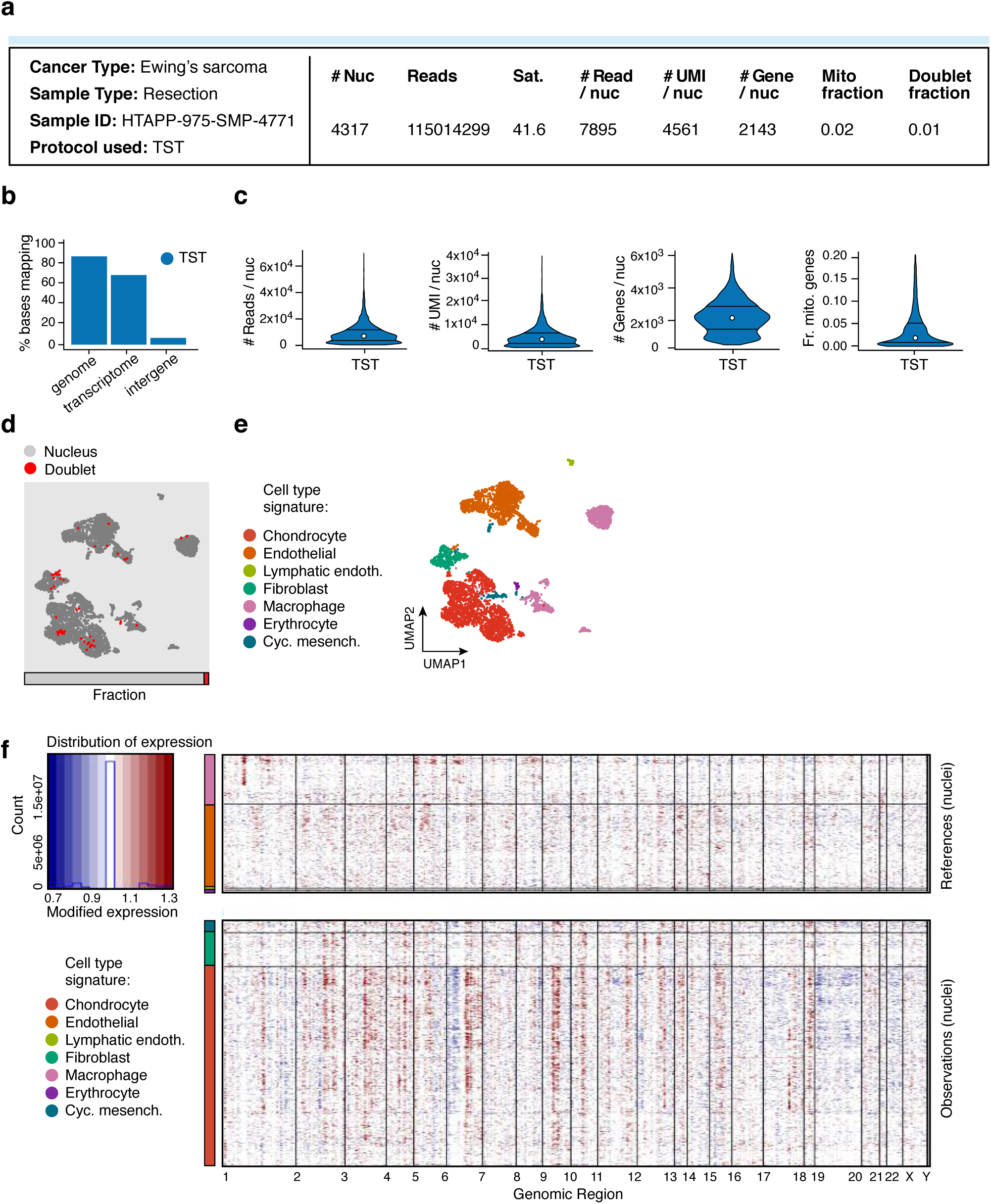
Evaluation of snRNA-Seq protocol for a sarcoma resection. **(A)** Sample processing and QC overview. Shown are the number of nuclei passing QC, the number of sequencing reads, and sequencing saturation across all nuclei. The remaining metrics are reported for those nuclei passing QC: median number of reads per nucleus, median number of UMIs per nucleus, median number of genes per nucleus, median fraction of UMIs mapping to mitochondrial genes in each nucleus, and fraction of nucleus barcodes called as doublets. **(B)** Read mapping QCs. The percent of bases in the sequencing reads (*y* axis) mapping to the genome, transcriptome, and intergenic regions (*x* axis). **(C)** Overall QCs. Distribution (median and first and third quartiles) of the number of reads per nucleus, number of UMIs per nucleus, number of genes per nucleus, and fraction of UMIs mapping to mitochondrial genes in each nucleus (*y* axes) for all nuclei passing QC. **(D)** Relation of doublets to cell types. UMAP embedding and fraction (horizontal bar) of single nucleus (grey) and doublet (red) profiles. **(E)** Cell type assignment. UMAP embedding of single nucleus profiles colored by assigned cell type signature. **(F)** Inferred CNA profiles for nuclei. Chromosomal amplification (red) and deletion (blue) inferred in each chromosomal position (columns) across the single nuclei (rows). Top: reference nuclei not expected to contain CNA in this cancer type. Bottom: nuclei tested for CNA relative to the reference nuclei. Color bar: assigned cell type signature for each nucleus.

**Supplementary Figure 28.**
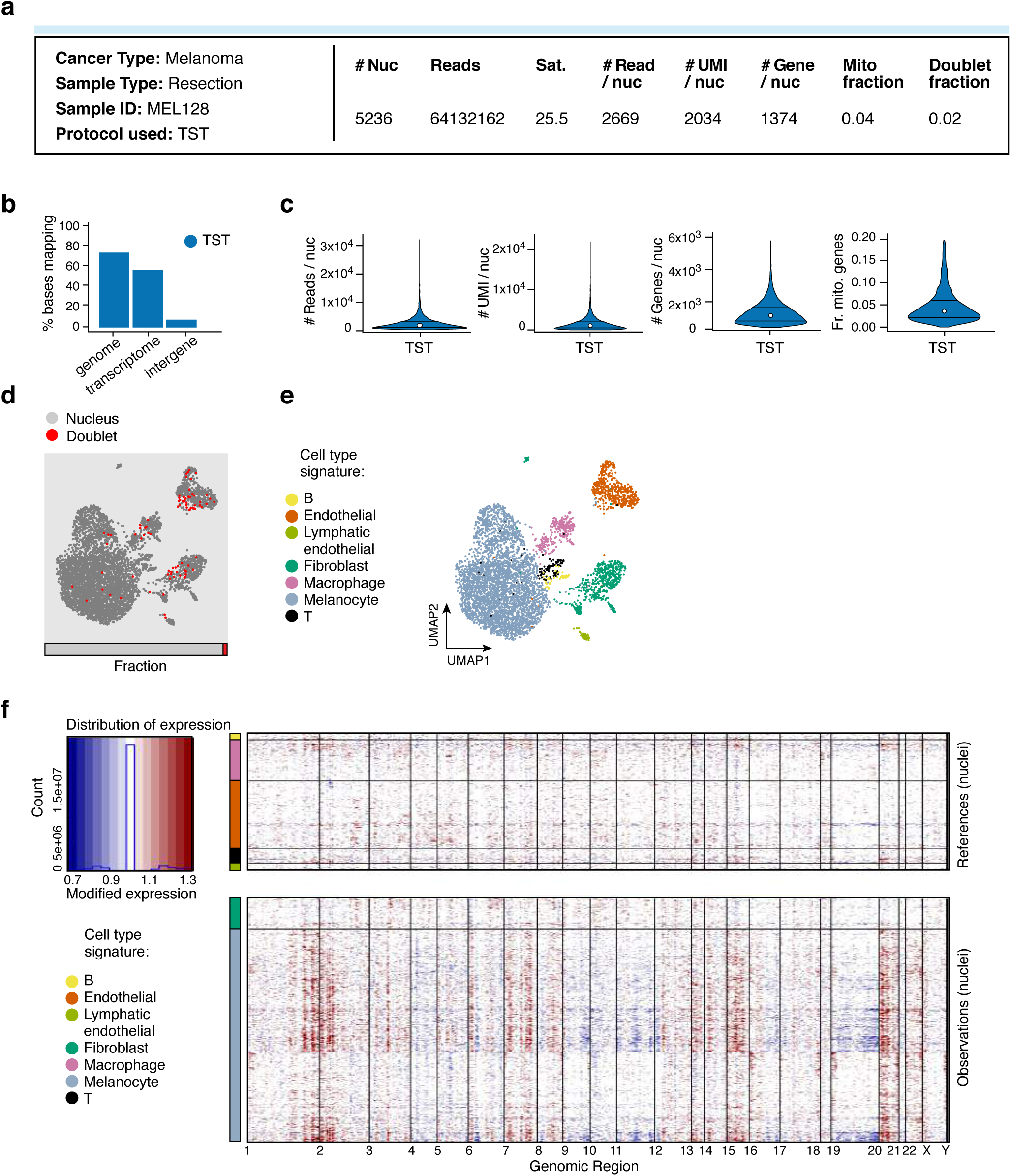
Evaluation of snRNA-Seq protocol for a melanoma resection. **(A)** Sample processing and QC overview. Shown are the number of nuclei passing QC, the number of sequencing reads, and sequencing saturation across all nuclei. The remaining metrics are reported for those nuclei passing QC: median number of reads per nucleus, median number of UMIs per nucleus, median number of genes per nucleus, median fraction of UMIs mapping to mitochondrial genes in each nucleus, and fraction of nucleus barcodes called as doublets. **(B)** Read mapping QCs. The percent of bases in the sequencing reads (*y* axis) mapping to the genome, transcriptome, and intergenic regions (*x* axis). **(C)** Overall QCs. Distribution (median and first and third quartiles) of the number of reads per nucleus, number of UMIs per nucleus, number of genes per nucleus, and fraction of UMIs mapping to mitochondrial genes in each nucleus (*y* axes) for all nuclei passing QC. **(D)** Relation of doublets to cell types. UMAP embedding and fraction (horizontal bar) of single nucleus (grey) and doublet (red) profiles. **(E)** Cell type assignment. UMAP embedding of single nucleus profiles colored by assigned cell type signature. **(F)** Inferred CNA profiles for nuclei. Chromosomal amplification (red) and deletion (blue) inferred in each chromosomal position (columns) across the single nuclei (rows). Top: reference nuclei not expected to contain CNA in this cancer type. Bottom: nuclei tested for CNA relative to the reference nuclei. Color bar: assigned cell type signature for each nucleus.

**Supplementary Figure 29.**
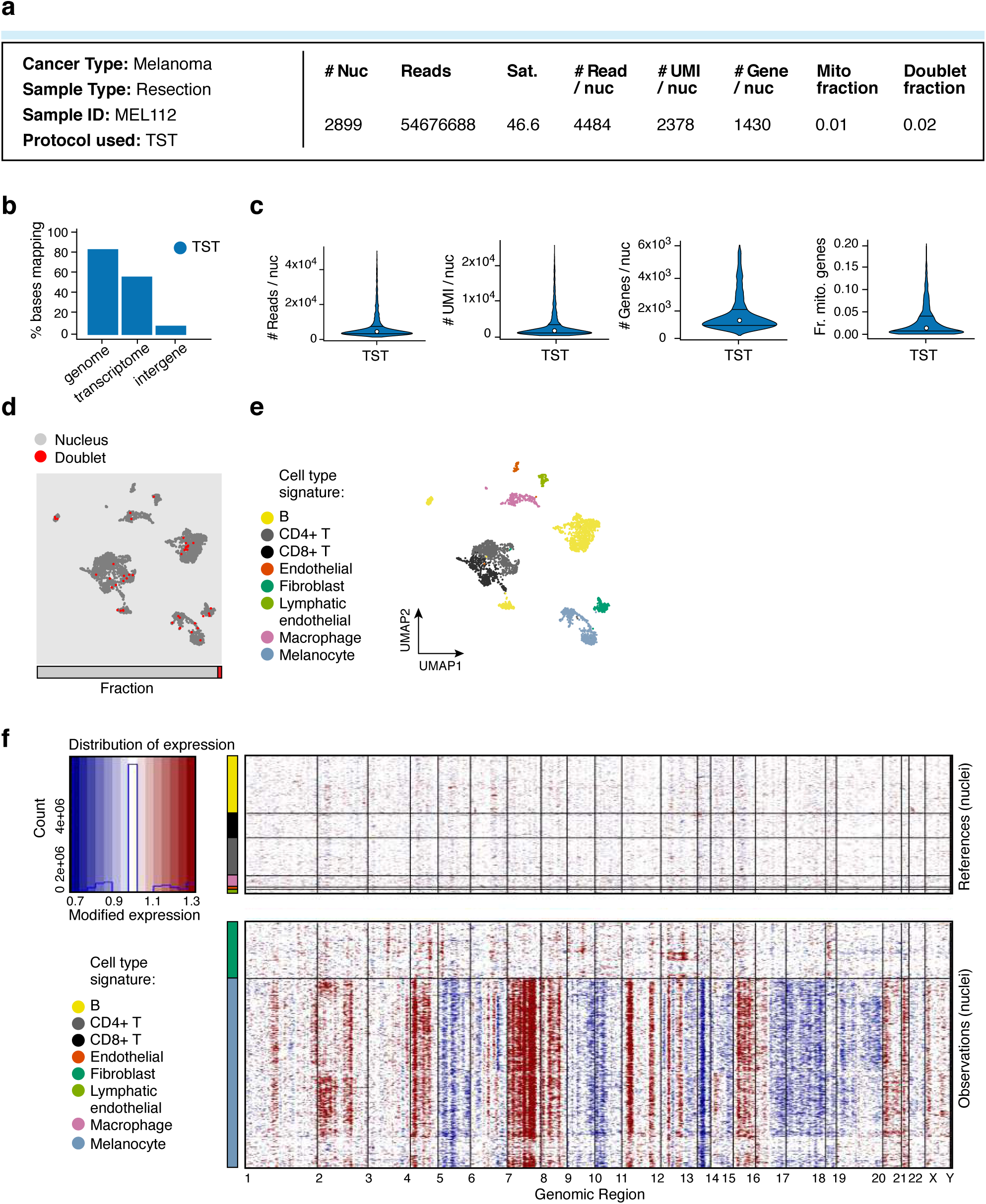
Evaluation of snRNA-Seq protocol for another melanoma resection. **(A)** Sample processing and QC overview. Shown are the number of nuclei passing QC, the number of sequencing reads, and sequencing saturation across all nuclei. The remaining metrics are reported for those nuclei passing QC: median number of reads per nucleus, median number of UMIs per nucleus, median number of genes per nucleus, median fraction of UMIs mapping to mitochondrial genes in each nucleus, and fraction of nucleus barcodes called as doublets. **(B)** Read mapping QCs. The percent of bases in the sequencing reads (*y* axis) mapping to the genome, transcriptome, and intergenic regions (*x* axis). **(C)** Overall QCs. Distribution (median and first and third quartiles) of the number of reads per nucleus, number of UMIs per nucleus, number of genes per nucleus, and fraction of UMIs mapping to mitochondrial genes in each nucleus (*y* axes) for all nuclei passing QC. **(D)** Relation of doublets to cell types. UMAP embedding and fraction (horizontal bar) of single nucleus (grey) and doublet (red) profiles. **(E)** Cell type assignment. UMAP embedding of single nucleus profiles colored by assigned cell type signature. **(F)** Inferred CNA profiles for nuclei. Chromosomal amplification (red) and deletion (blue) inferred in each chromosomal position (columns) across the single nuclei (rows). Top: reference nuclei not expected to contain CNA in this cancer type. Bottom: nuclei tested for CNA relative to the reference nuclei. Color bar: assigned cell type signature for each nucleus.

**Supplementary Figure 30.**
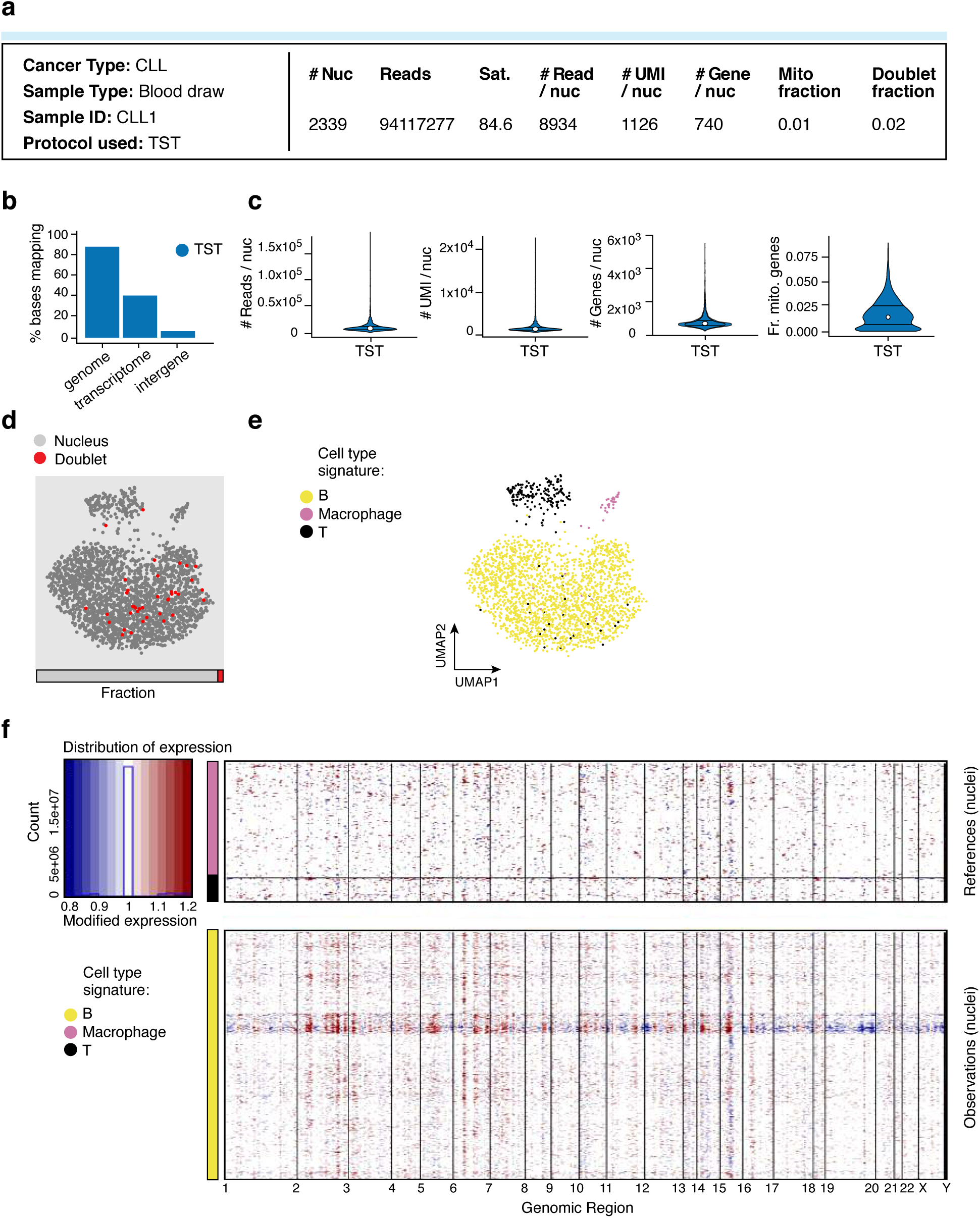
Evaluation of snRNA-Seq protocol for a cryopreserved CLL sample. **(A)** Sample processing and QC overview. Shown are the number of nuclei passing QC, the number of sequencing reads, and sequencing saturation across all nuclei. The remaining metrics are reported for those nuclei passing QC: median number of reads per nucleus, median number of UMIs per nucleus, median number of genes per nucleus, median fraction of UMIs mapping to mitochondrial genes in each nucleus, and fraction of nucleus barcodes called as doublets. **(B)** Read mapping QCs. The percent of bases in the sequencing reads (*y* axis) mapping to the genome, transcriptome, and intergenic regions (*x* axis). **(C)** Overall QCs. Distribution (median and first and third quartiles) of the number of reads per nucleus, number of UMIs per nucleus, number of genes per nucleus, and fraction of UMIs mapping to mitochondrial genes in each nucleus (*y* axes) for all nuclei passing QC. **(D)** Relation of doublets to cell types. UMAP embedding and fraction (horizontal bar) of single nucleus (grey) and doublet (red) profiles. **(E)** Cell type assignment. UMAP embedding of single nucleus profiles colored by assigned cell type signature. Note that the cell type signature used for macrophages contains macrophage and monocyte markers. **(F)** Inferred CNA profiles for nuclei. Chromosomal amplification (red) and deletion (blue) inferred in each chromosomal position (columns) across the single nuclei (rows). Top: reference nuclei not expected to contain CNA in this cancer type. Bottom: nuclei tested for CNA relative to the reference nuclei. Color bar: assigned cell type signature for each nucleus.

**Supplementary Figure 31.**
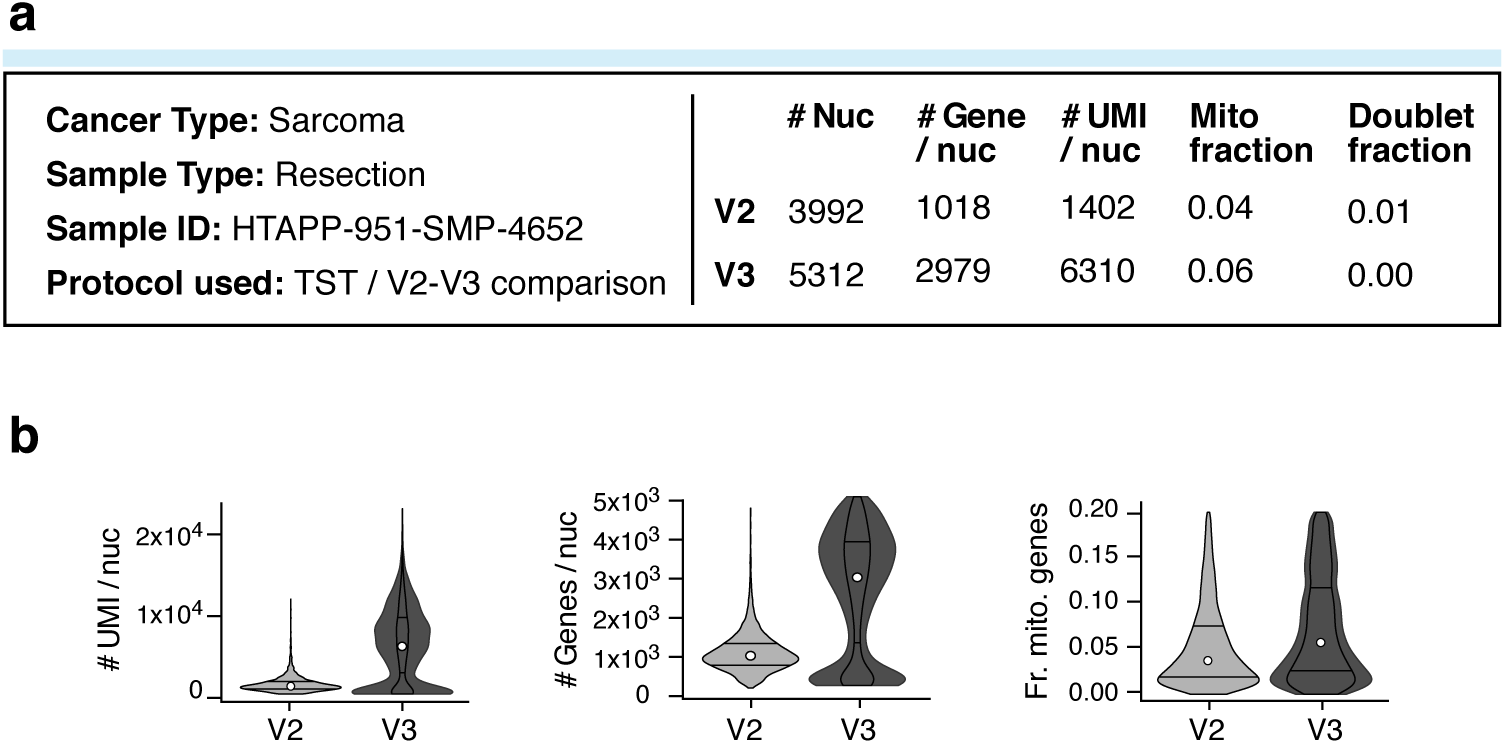
SnRNA-seq protocol comparison of V2 and V3 chemistry from 10x Genomics on a resection of sarcoma. **(A)** Sample processing and QC overview. For each protocol, shown are the number of nuclei passing QC, after the total number of sequencing reads from the V3 protocol data was down-sampled to match the number of reads in the V2 data. The remaining metrics are reported for those nuclei passing QC: median number of UMIs per nucleus, median number of genes per nucleus, median fraction of UMIs mapping to mitochondrial genes in each nucleus, and fraction of nucleus barcodes called as doublets. **(B)** Overall QCs. Distribution (median and first and third quartiles) of number of UMIs per nucleus, number of genes per nucleus, and fraction of UMIs mapping to mitochondrial genes in each nucleus (*y* axes) for all nuclei passing QC.

**Supplementary Figure 32.**
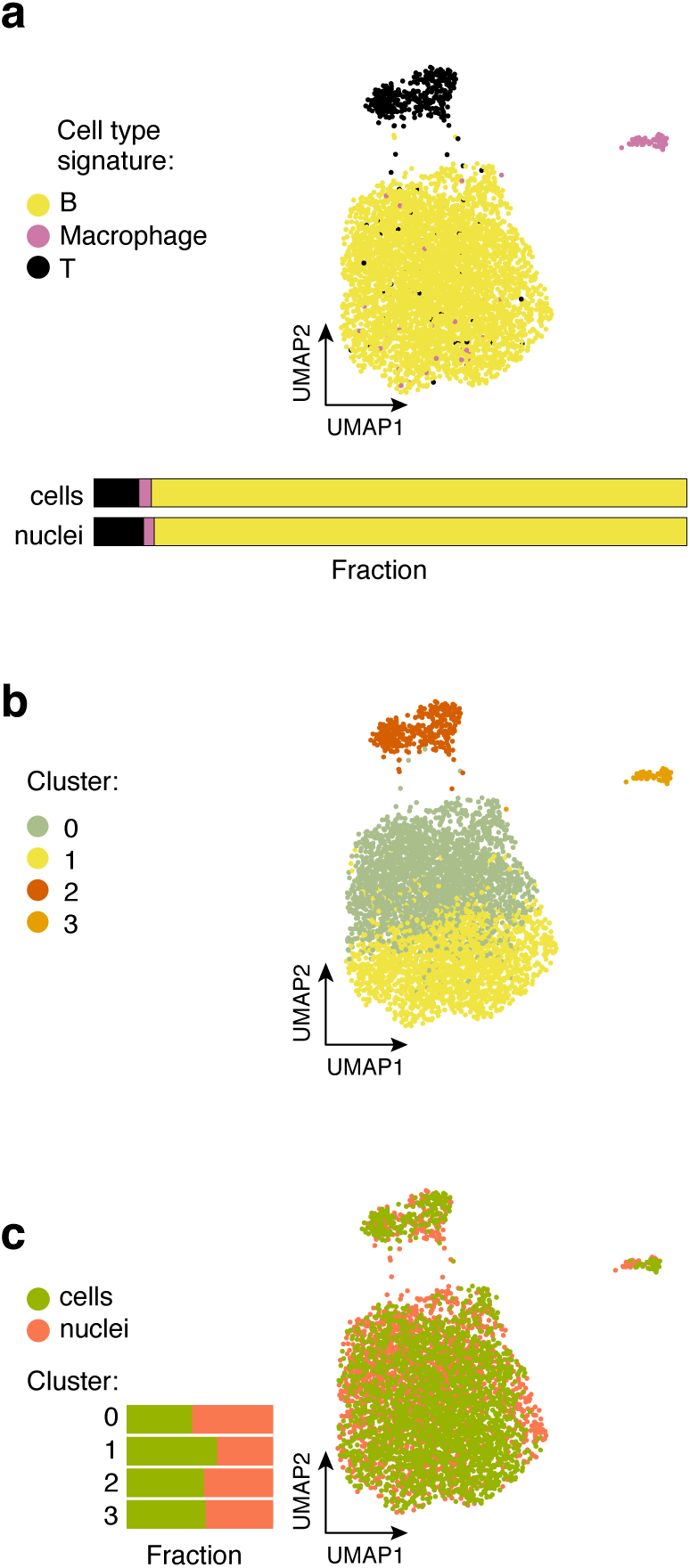
Comparison of scRNA-Seq and snRNA-Seq from a single cryopreserved CLL sample. **(A-C)** UMAP embedding of single cell and single nucleus profiles after batch correction by CCA (**Methods**) colored by either assigned cell type signature (**A**; fractions in horizontal bar), cluster assignment (**B**) or data type (**C**, cells or nuclei; horizontal bar: cluster assignment).

**Supplementary Figure 33.**
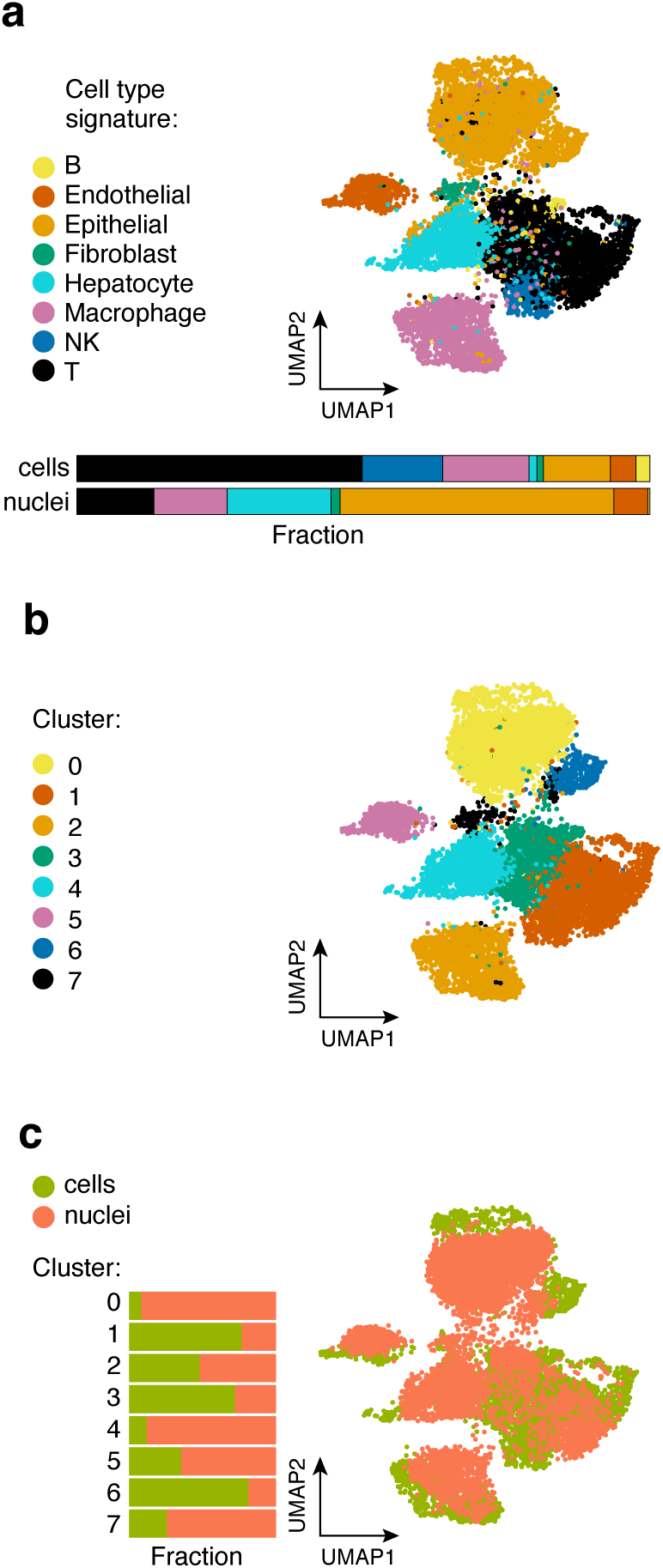
Comparison of scRNA-Seq and snRNA-Seq from a single metastatic breast cancer sample (HTAPP-963-SMP-4741). (A-C) UMAP embedding of single cell and single nucleus profiles after batch correction by CCA (**Methods**) colored by either assigned cell type signature (**A**; fractions in horizontal bar), cluster assignment (**B**) or data type (**C**, cells or nuclei; horizontal bar: cluster assignment).

**Supplementary Figure 34.**
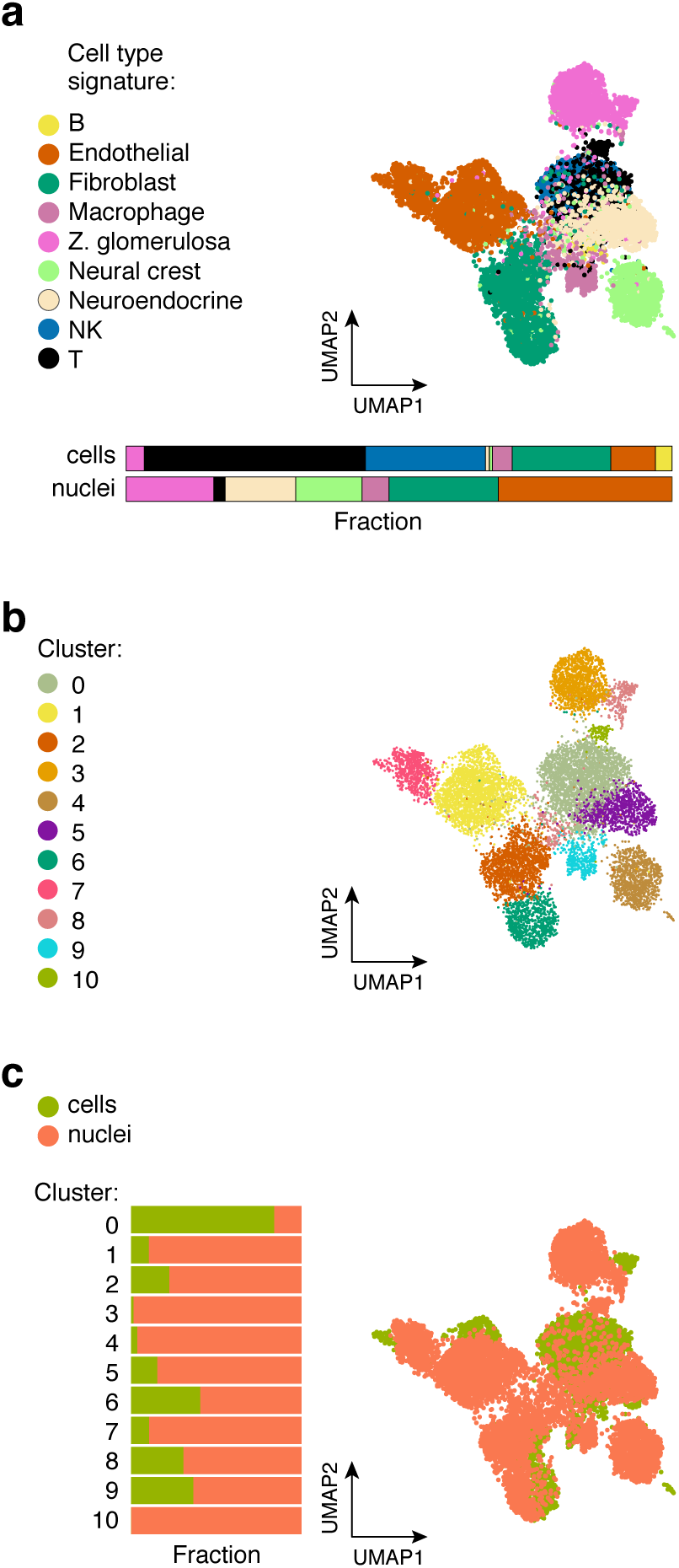
Comparison of scRNA-Seq and snRNA-Seq from a single neuroblastoma sample (HTAPP-656-SMP-3481). **(A-C)** UMAP embedding of single cell and single nucleus profiles after batch correction by CCA (**Methods**) colored by either assigned cell type signature (**A**; fractions in horizontal bar), cluster assignment (**B**) or data type (**C**, cells or nuclei; horizontal bar: cluster assignment).

**Supplementary Figure 35.**
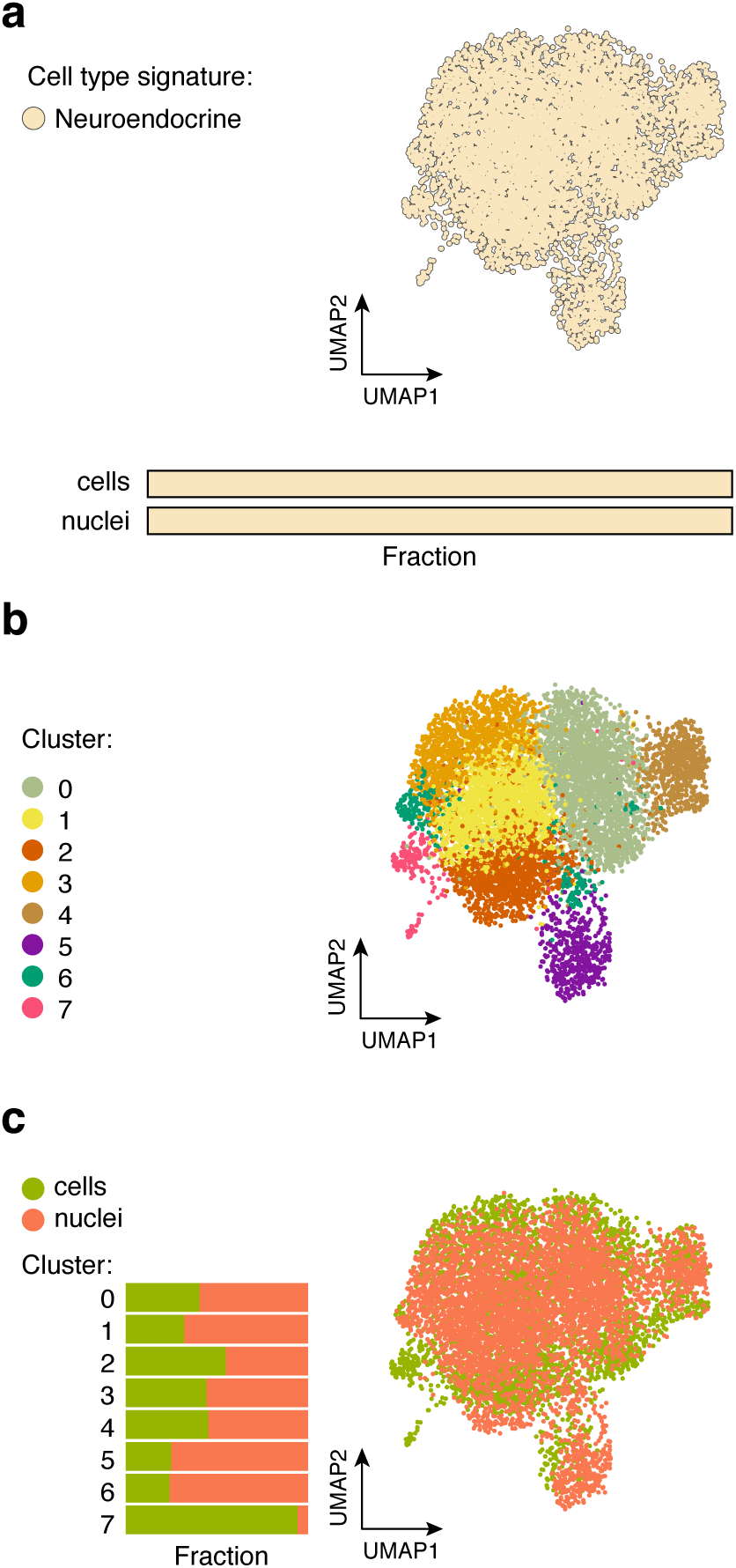
Comparison of scRNA-Seq and snRNA-Seq from a single O-PDX neuroblastoma sample. **(A-C)** UMAP embedding of single cell and single nucleus profiles after batch correction by CCA (**Methods**) colored by either assigned cell type signature (**A**; fractions in horizontal bar), cluster assignment (**B**) or data type (**C**, cells or nuclei; horizontal bar: cluster assignment).

## Methods

### Human patient samples

External sample cohorts were added to the Broad Institute’s Molecular Classification of Cancer protocol (15-370B) and reviewed and approved by the Dana Farber Cancer Institute’s Institutional Review Board (IRB). No subject recruitment or ascertainment was performed as part of the Broad protocol. Samples added to this protocol also underwent IRB review and approval at the institutions where the samples were originally collected. Specifically, Dana-Farber Cancer Institute IRB approved the following protocols: lung cancer (IRB protocol 98-063), metastatic breast cancer (IRB protocol 05-246), neuroblastoma (IRB protocols 11-104 and 17-104), ovarian cancer (IRB protocol 02-051), melanoma (IRB protocol 11-104), sarcoma (IRB protocol 17-104), GBM (IRB protocol 10-417), and chronic lymphocytic leukemia (IRB protocol 99-224), and the St. Jude Children’s Research Hospital IRB approved the following protocol: pediatric high-grade glioma (IRB protocol 97BANK).

The XPD 09-234 MAST (Molecular Analysis of Solid Tumor) protocol for creating the neuroblastoma O-PDX sample was reviewed and approved by the St. Jude Children’s Research Hospital IRB.

### Laboratory animals

For the neuroblastoma O-PDX sample, animal use was restricted to one female nude athymic mouse for para-adrenal injection of O-PDX cells. This study was carried out in strict accordance with the recommendations in the Guide to Care and Use of Laboratory Animals of the National Institute of Health. The protocol was approved by the Institutional Animal Care and Use Committee at St. Jude Children’s Research Hospital. All efforts were made to minimize suffering. All mice were housed in accordance with approved IACUC protocols. Animals were housed on a 12-12 light cycle (light on 6 am and off 6 pm) and provided food and water ad libitum. Athymic nude female mice were purchased from Charles River Laboratories (strain code 553).

### Collection of fresh tissue for scRNA-Seq

Collection of fresh solid tumor tissue for lung cancer, ovarian cancer, and metastatic breast cancer at BWH/DFCI, was performed following protocols established to reduce the time elapsed between removal of the tumor tissue from the body, placement of the specimen in media, and processing for scRNA-Seq. To this end, we established procedures between the hospital team (surgeon/clinical research coordinator (CRC)/clinical pathologist), the coordinating team (project managers/ pathology technician), and the processing team (staff scientists/research technicians) prior to procedure day. This included providing the hospital team with collection containers with appropriate media and pre-defining allocation priorities to ensure quick handling by the pathology technician of the sample received. On the day of the procedure, timely communication between the teams ensured quick specimen transfer from the hospital team to the research team, timely transport to the Broad Institute for processing, and immediate loading of the single cell suspension into the 10x Genomics Single-Cell Chromium Controller (below).

In all cases, the tissue received from the hospital team was examined by the research pathology technician and following procurement of a specimen for anatomic pathology review, the highest quality portion (or core) was allocated for scRNA-Seq, placed in media and transported to the Broad Institute for dissociation following the appropriate protocol (below). Tissue quality is assessed based on visual examination and rapid pathology interpretation at the time of collection, and determined based on tumor content, necrosis, calcification, fat, and hemorrhage.

For ovarian cancer ascites, approximately ∼300 mL were usually received from the hospital team within one hour after taken out of the body, which contained a vast majority of non-malignant (mainly immune) cells. Hence, all ascites samples were subjected to CD45^+^ cell depletion (below) to enrich for malignant cells.

For CLL, samples were generated from peripheral blood mononuclear cells isolated using density centrifugation (Ficoll-Paque) and stored in freezing media (FBS +10% DMSO) in liquid nitrogen until processing.

For orthotopic PDX of neuroblastoma samples (O-PDX), *Foxn1-/-* nude mice (Charles River Laboratories) were orthotopically injected via ultrasound-guided para-adrenal injection with cells derived from a patient MYCN-amplified neuroblastoma (available as sample SJNBL046_X1 through the Childhood Solid Tumor Network)(Stewart et al., 2017; Stewart et al., 2015). A portion of O-PDX tumor was flash-frozen for future single-nucleus RNA-Seq, while the remainder underwent dissociation as described below.

### Preservation of tissue for snRNA-Seq

For those samples that we prospectively collected for snRNA-seq (Neuroblastoma HTAPP-244-SMP-451, HTAPP-656-SMP-3481), freezing of tumor samples was performed as quickly as possible after sample collection using standard biobanking technique and the dates when samples were frozen were recorded. (Other samples were obtained from tissue banks with limited record on how they were frozen, which is a typical scenario.) Samples were placed in cryo-tubes without any liquid. Complete removal of liquid from the sample was accomplished by gently wiping it (not patting, as this would damage the tissue) on the side of the container, before placing in the cryotube. The tubes were then covered in dry-ice and transferred to −80°C for long term storage.

The other frozen samples from snRNA-Seq were obtained from tissue banks as follows: ovarian OCT-frozen archival samples were obtained from the Dana-Farber Cancer Institute Gynecology Oncology Tissue Bank; sarcoma snap-frozen samples were obtained from the Boston Children’s Hospital Tissue Bank; pediatric snap-frozen glioma samples were obtained from the St. Jude Children’s Research Hospital Biorepository; neuroblastoma snap-frozen samples were obtained from the St. Jude Children’s Research Hospital Biorepository and the Boston Children’s Hospital Precision Link Biobank for Health Discovery; metastatic breast cancer OCT-frozen samples were obtained from the Center for Cancer Precision Medicine Bank; snap-frozen melanoma samples were obtained through the laboratory of Dr. Charles Yoon at BWH.

### Dissociation workflow from fresh solid tumor samples to a single-cell suspension for scRNA-seq

#### MBC, NSCLC (protocols PDEC and LE), ovarian cancer solid tumor, and neuroblastoma workflows

Fresh tissue dissociation of MBC, NSCLC (protocols PDEC and LE), ovarian cancer solid tumor, and neuroblastoma were performed using a similar workflow (**Fig. 2A**), with different components of the dissociation mixture for each tumor type, as described in the next section.

Samples were transferred from interventional radiology (biopsies) or the operating room (resections) in DMEM (MBC), RPMI (NSCLC), or RPMI with HEPES (ovarian cancer and neuroblastoma) medium. Upon arrival to the laboratory, the sample was washed in cold PBS and transferred into either a 2 mL Eppendorf tube containing dissociation mixture (for biopsies) or a 5 mL Eppendorf tube containing 3 mL dissociation mixture (for resections). Next, the sample was minced in the Eppendorf tube using spring scissors (Fine Science Tools, catalog no. 15514-12) into fragments under ∼0.4 mm, and incubated at 37°C, while rotating at approximately 14 RPM, for 10 minutes. After 10 minutes, the sample was pipetted 20 times with a 1 mL pipette tip at room temperature, and placed back into incubation with rotation for an additional 10 minutes. The sample was pipetted again 20 times using a 1 mL pipette tip, transferred to a 1.7 mL Eppendorf tube and centrifuged at 300 g-580 g for 4-7 minutes at 4°C. The supernatant was removed and the pellet was resuspended in 200-500 µL of ACK red blood cell lysis buffer (Thermo Fisher Scientific, A1049201). The ACK volume added depended on the size of the pellet; while pellet size is hard to quantify we suggest adding about 100 µL ACK lysis buffer per 100,000 cells, with a minimum volume of 200 µL. The sample was incubated in ACK red blood cell lysis buffer for 1 minute on ice, followed by the addition of cold PBS at twice the volume of the ACK. The cells were pelleted by a short centrifugation for 8 seconds at 4°C using the short spin setting with centrifugal force ramping up to, but not exceeding, 11,000 g. The supernatant was removed. The pellet color was assessed, if RBCs remained (pellet color pink or red), the ACK step was repeated up to two additional times. To remove cell clumps in the MBC protocol (or sample), the pellet was resuspended in 100 µL of TrypLE (Life Technologies, catalog no. 12604013) and incubated while constantly pipetting at room temperature for 1 minute with a 200 µL pipette tip. TrypLE was inactivated by adding 200 µL of cold RPMI 1640 with 10% FBS. The cells were pelleted using short centrifugation as described above. The pellet was resuspended in 50 µL of 0.4% BSA (Ambion, catalog no. AM2616) in PBS. To assess the single cell suspension, viability, and cell count, 5 µL of Trypan blue (Thermo Fisher Scientific, catalog no. T10282) was mixed with 5 µL of the sample and loaded on INCYTO C-Chip Disposable Hemocytometer, Neubauer Improved (VWR, catalog no. 82030-468). The cell concentration was adjusted if necessary to a range of 200-2,000 cells/µL. A total of 8,000 cells were loaded into each channel of the 10x Genomics Single-Cell Chromium Controller. Due to differences between clinical samples, some steps may need to be repeated or adjusted; for a general overview of guidelines see **Fig. 2A**.

#### NSCLC-C4 protocol workflow

A similar workflow was used for protocol NSCLC-C4 with the following modifications: Following mechanical chopping as above, sample was dissociated for 15 minutes in a 15 mL falcon tube, with gentle vortex every 5 minutes, followed by filtration through a 70 µm filter, and washed with 20 mL of ice cold PBS and centrifuged at 580 g for 5 minutes. RBS lysis was performed similarly to the above workflow by resuspending the pellet in 1 mL ACK lysis buffer with incubation on ice for 1 minute. 20 mL of ice cold PBS were added to quench the ACK lysis buffer, followed by filtration through a 70 µm filter, and centrifugation at 580 g for 5 minutes. Sample NSCLC14 was further cleaned using Viahance™ dead-cell removal kit (BioPAL, catalog no. CP-50VQ02) according to manufacturer’s instructions. Cells were then re-suspended in M199 and loaded on the 10x Genomics Single-Cell Chromium Controller as described above.

#### GBM workflow

All steps were done on ice. Sample was minced thoroughly in Petri dish, thereafter, 4 mL HBSS were added (Life Technologies, catalog number 14175095), transferred to 15 mL tubes and centrifuged at 1000 rpm for 2 minutes. After centrifugation, supernatant was removed, pre-heated Buffer X was added, and the sample was incubated while shaking at 37°C for 15 minutes. Sample was pipetted up-down 20 times, incubated at 37°C for an additional 15 minutes, and pipetted again. After dissociation, the sample was filtered through a 100 μm cell strainer (Fisher Scientific, Cat # 22-363-547) into 50 mL tube. We recommend keeping any tissue fragments left in the cell strainer, as they can be reprocessed with the same protocol if initial cell recovery is low. Filtrate was centrifuged at 1000 rpm for 3 minutes, and the supernatant was removed. If the pellet was bloody, RBC removal was performed when needed using LYMPHOLYTE H (CedarLAne, Cat.# CL5015) or Red Blood Cell (RBC) Lysis Solution (10x) (Miltenyi Biotech, Cat# 130-094-183). The pellet was washed with 10 mL of cold PBS/1% BSA, transferred to 15 mL tube and centrifuged at 1200 rpm for 3 minutes. Supernatant was removed and the pellet was resuspended in 0.4 BSA in PBS. Single cell suspension was visualized, counted and loaded on the 10x Genomics Single-Cell Chromium Controller as described above.

### Dissociation mixtures for different tumor types

Dissociation mixtures were prepared approximately 5-10 minutes before sample processing from frozen aliquoted stocks, as follows:

#### MBC, LD protocol

950 µL of RPMI 1640 (Thermo Fisher Scientific, catalog no. 11875093), 10 µL of 10 mg/mL DNAse I (Sigma Aldrich, catalog no. 11284932001) to a final concentration of 100 µg/mL, and 40 µL of 2.5 mg/mL Liberase TM.

#### Ovarian cancer resection

Dissociation mixture was based on Miltenyi Human Tumor Dissociation Kit (Miltenyi Biotec, catalog no. 130-095-929). Before starting, Enzymes H, R, and A were resuspended according to manufacturer’s instructions. Dissociation mix containing 2.2 mL RPMI, 100 µL enzyme H, 50 µL enzyme R, and 12.5 enzyme A, was prepared immediately before use.

#### Neuroblastoma, NB-C4 protocol

Medium 199, Hanks Balanced Salts Buffer (Thermo Fisher Scientific) with 100 µg/mL of DNAse I (Millipore Sigma, catalog no. 11284932001), 100 µg/mL Collagenase IV (Worthington; catalog no. LS004186).

#### Orthotopic PDX neuroblastoma

Worthington Papain Dissociation System (catalog no. LK003150). Dissociation was performed according to manufacturer’s instructions, with deviation of the dissociation duration, which was shortened to 15 minutes.

#### NSCLC, PDEC protocol

2692 HBSS (Thermo Fisher Scientific, catalog no. 14170112), 187.5 µL of 20 mg/mL pronase (Sigma Aldrich, catalog no. 10165921001) to a final concentration of 1,250 µg/mL, 27.6 µL of 1 mg/mL elastase (Thermo Fisher Scientific, catalog no. NC9301601) to a final concentration of 9.2 µg/mL, 30 µL of 10 mg/mL DNase I (Sigma Aldrich, catalog no. 11284932001) to a final concentration of 100 µg/mL, 30 µL of 10 mg/mL Dispase (Sigma Aldrich, catalog no. 4942078001) to a final concentration of 100 µg/mL, 30 µL of 150 mg/mL Collagenase A (Sigma Aldrich, catalog no. 10103578001) to a final concentration of 1,500 µg/mL, 3 µL of 100 µg/mL collagenase IV (Thermo Fisher Scientific, catalog no. NC9836075) to a final concentration of 1250 µg/mL.

#### NSCLC, LE protocol

5 mL RPMI 1640 (Thermo Fisher Scientific, catalog no. 11875093), 200 µL of 2.5 mg/mL Liberase TM (Millipore Sigma, 5401119001) to a final concentration of 100 µg/mL, 50 µL of 10 mg/mL DNase I (Sigma Aldrich, catalog no. 11284932001) to a final concentration of 100 µg/mL, 27.6 µL of 1 mg/mL elastase (Thermo Fisher Scientific, catalog number NC9301601) to a final concentration of 9.2 µg/mL.

#### NSCLC, C4 protocol

5 mL M199 with DNase 1 (final concentration of 10 µg/mL) and Collagenase iv (final concentration of 100 µg/mL).

#### GBM

Brain Tumor Dissociation Kit (P) (Miltenyi Biotech. Catalog number 130-095-942). 4 mL Buffer X, 40 µL Buffer Y, 50 µL Enzyme N, 20 µL Enzyme A.

### Processing of non-solid tumor samples for scRNA-Seq

#### CLL

Frozen (cryopreserved) cells were thawed in 10 mL RPMI, pelleted and washed with an additional 10 mL RPMI. Live cells were sorted using the MoFlo Astrios EQ Cell Sorter, and 8,000 cells were loaded on one channel of the 10x Genomics Single-Cell Chromium Controller. Remaining cells were pelleted by short centrifugation, the supernatant was discarded and the pellet was frozen on dry ice and stored in −80°C.

#### Ovarian cancer ascites

Ascites samples without spheres were selected and delivered in four 50 mL conical tubes, for a total of 200 mL of fluid. Tubes were spun down at 580 g for 5 minutes in a 4°C pre-cooled centrifuge and supernatants was aspirated.

Pellets were resuspended in 5 mL cold ACK Lysing Buffer, and combined from all tubes at this step. ACK lysis was done on ice for 3 minutes, and quenched by adding 10 mL of cold PBS, followed by centrifugation at 580 g for 5 minutes at 4°C. The pellet color was assessed and if it was pink or red, revealing a significant portion of erythrocytes, ACK treatment steps were repeated as needed at most two additional times. Post ACK treatment, the pellet was resuspended in 20 mL cold PBS, filtered through a 70 µm cell strainer into a 50 mL conical tube, and the filter was washed with additional 20 mL cold PBS to recover as many cells as possible. The sample was then centrifuged at 580 g for 5 minutes at 4°C. To reduce the fraction of immune cells in the sample, CD45^+^ cell depletion was performed using the MACS CD45 depletion protocol described below.

### Depletion of CD45^+^ cells for scRNA-Seq

Depletion of CD45^+^ cells in ovarian cancer ascites samples and NSCLC was performed using CD45 MicroBeads (Miltenyi Biotec, catalog no. 130-045-801) according to the manufacturer’s protocol. Briefly, following filtration of the ovarian cells from ascites or dissociation of NSCLC tissue samples, cells were counted. The single-cell suspension was centrifuged at 500 g for 4 minutes at 4°C. The supernatant was removed and the pellet was resuspended in 80 µL of MACS buffer (PBS supplemented with 0.5% BSA, and 2mM EDTA) per 10^6^ cells. 20 µL of the MACS CD45 microbeads were added to the cell suspension per 10 million cells. The cells incubated on ice for 15 minutes. During the incubation, the column (MS for NSCLC and LS for ovarian ascites) was prepared by attaching the column to a MidiMACS separator and rinsing the column with 3 mL MACS buffer. Following the incubation, the cells and bead conjugate was washed with 900 µL MACS buffer per 10 million cells. The cells were centrifuged at 500 g for 4 minutes at 4°C. The supernatant was removed and the pellet was resuspended in 500 µL MACS buffer. The cell suspension was transferred to the column and the effluent was collected (CD45^-^ fraction). The column was washed three times with 3 mL MACS buffer. The CD45^-^ fraction was centrifuged at 500 g for 4 minutes at 4°C. In the ascites sample, bead attachment and column separation can be repeated to increase the number of tumor and stromal cells relative to immune cells. The pellet was resuspended in 50 µL of 0.4% BSA (Ambion, catalog no. AM2616) in PBS. Cells were counted by mixing 5 µL of Trypan blue (Thermo Fisher Scientific, catalog no. T10282) with 5 µL of the sample and loaded on INCYTO C-Chip Disposable Hemocytometer, Neubauer Improved (VWR, catalog no. 82030-468). The cell concentration was adjusted if necessary to a range of 200-2,000 cells/µL. 8,000 cells were loaded into each channel of the 10x Genomics Single-Cell Chromium Controller.

### Flow cytometry analysis

For flow cytometry analysis of CD45^+^ depletion in the ovarian cancer ascites sample, cells were resuspended in PBS complemented with 2% fetal bovine serum and stained with FITC anti-human CD45 antibody (BioLegend #304006CD45, 1:200 dilution) and PE anti-human EPCAM antibody (Miltenyi Biotech #130-113-264, 1:50 dilution) for 20 minutes, and with 7-AAD (Invitrogen #A1310, 1:200 dilution) for 5 minutes. The same cells were also used for single-stain and unstained controls in order to perform compensation and adjust gating. Analysis was performed on a BD LSRFortessa cell analyzer with BD FACSDiva Software Version 8.0.1 and plots were generated with FlowJo Version 10.5.3. Gating was performed as described in **Supp. Fig. 5**.

### ST based buffers for snRNA-seq

2X stock of salt-Tris solution (ST buffer) containing 292 mM NaCl (Thermo Fisher Scientific, catalog no. AM9759), 20 mM Tris-HCl pH 7.5 (Thermo Fisher Scientific, catalog no. 15567027), 2 mM CaCl2 (VWR International Ltd, catalog no. 97062-820), and 42 mM MgCl_2_ (Sigma–Aldrich, catalog no. M1028) in ultrapure water was made and used to prepare three buffers. <u>For CST</u>: 1 mL of 2X ST buffer, 980 µ of 1% CHAPS (Millipore), 10 µ of 20 mg/mL BSA (New England BioLabs), and 10 µ of nuclease-free water. <u>For TST</u>: 1 mL of 2X ST buffer, 60 µ of 1% Tween-20 (Sigma-aldrich, catalog no. P-7949), 10 µ of 20 mg/mL BSA (New England Biolabs, catalog no. B9000S), and 930 µ of nuclease-free water. <u>For NST</u>: 1 mL of 2X ST buffer, 40 µ of 10% Nonidet™ P40 Substitute (Fisher Scientific), 10 µ of 20 mg/mL BSA (NEB), and 950 µ of nuclease-free water. 1x ST buffer was prepared by dilution 2x ST with ultra-pure water (Thermo Fisher Scientific catalog no. 10977023) in a ratio of 1:1.

### Nucleus isolation from frozen samples for snRNA-seq

On dry ice, tissue was split and subjected to one of three salt-Tris (ST)-based nucleus isolation protocols (Drokhlyansky et al., 2019) and the EZ nuclei isolation buffer (Habib et al., 2017), as detailed below.

#### Nucleus isolation workflow for ST-based buffers

On ice, a piece of frozen tumor tissue was placed into a well of a 6-well plate (Stem cell Technologies, catalog no. 38015) with 1 mL of either CST, TST, or NST buffer. For samples frozen in OCT, an additional step of removing the surrounding OCT, and washing any residual OCT from the sample with PBS was performed in a 10 cm Petri dish. Tissue was then chopped using Noyes Spring Scissors (Fine Science Tools, catalog no. 15514-12) for 10 minutes on ice. For cell pellets, such as for CLL frozen cells, sample was pipetted in the buffer on ice, instead of chopping. The homogenized solution was then filtered through a 40 µm Falcon™ cell strainer (Thermo Fisher Scientific, catalog no. 08-771-1). An additional 1 mL of the detergent buffer solution was used to wash the well and filter. The volume was brought up to 5 mL with 3 mL of 1X ST buffer. The sample was then transferred to a 15 mL conical tube and centrifuged at 4°C for 5 minutes at 500 g in a swinging bucket centrifuge. The pellet was resuspended in 1X ST buffer. Resuspension volume was dependent on the size of the pellet, usually within the range of 100-200 µL. The nucleus solution was then filtered through a 35 µm Falcon™ cell strainer (Corning, catalog no. 352235). Nuclei were counted using C-chip disposable hemocytometer (VWR, International Ltd, catalog no. 82030-468). 10,000 or 8,000 nuclei (V2 or V3 10x genomics, receptively) of the single-nucleus suspension were loaded onto the Chromium Chips for the Chromium Single Cell 3′ Library (V2, PN-120233; V3 PN-1000075) according to the manufacturer’s recommendations (10x Genomics).

#### Nucleus isolation workflow using EZ lysis buffer

Nucleus isolation was done as previously described(Habib et al., 2017). Briefly, tissue samples were cut into pieces <0.5 cm and homogenized using a glass dounce tissue grinder (Sigma, Catalog no. D8938). The tissue was homogenized 25 times with pestle A and 25 times with pestle B in 2 mL of ice-cold nuclei EZ lysis buffer. The sample was then incubated on ice for 5 minutes, with an additional 3 mL of cold EZ lysis buffer. Nuclei were centrifuged at 500 g for 5 minutes at 4°C, washed with 5 mL ice-cold EZ lysis buffer and incubated on ice for 5 minutes. After centrifugation, the nucleus pellet was washed with 5 mL Nuclei Suspension Buffer (NSB; consisting of 1x PBS, 0.01% BSA and 0.1% RNAse inhibitor (Clontech, Catalog no.2313A)). Isolated nuclei were resuspended in 2 mL NSB, filtered through a 35 μm cell strainer (Corning-Falcon, Catalog no. 352235) and counted. A final concentration of 1,000 nuclei/µL was used for loading on 10x v2 channel.

### Droplet-based sc/snRNA-seq

An input of 8,000 single cells or 10,000 single nuclei (8,000 for v3 10x technology) were loaded into each channel of the Chromium single cell 3’ Chip. Single cells/nuclei were partitioned into droplets with Gel Beads in the Chromium. After emulsions were formed, barcoded reverse transcription of RNA took place. This was followed by cDNA amplification, fragmentation and adaptor and sample index attachment, all according to the manufacturer’s recommendations.

Libraries from four 10x channels were pooled together and sequenced on one lane of an Illumina HiSeqX with paired end reads, Read 1: 26 nt, Read 2: 55 nt, Index 1: 8 nt, Index 2: 0 nt.

### scRNA-seq data processing

We used Cell Ranger `mkfastq` (v2.0 and v3.0) (10x Genomics) to generate demultiplexed FASTQ files from the raw sequencing reads. We aligned these reads to the human GRCh38 genome and quantified gene counts as UMIs using Cell Ranger `count` (v2.0 and v3.0) (10x Genomics). For single-nucleus RNA-seq reads, we counted reads mapping to introns as well as exons, as this results in a greater number of genes detected per nucleus, more nuclei passing quality control, and better cell type identification, as previously described (Bakken et al., 2018). To count introns during read mapping, we followed the approach described at https://support.10xgenomics.com/single-cell-gene-expression/software/pipelines/latest/advanced/references. Briefly, we built a “pre-mRNA” human GRCh38 reference using Cell Ranger `mkref` (v3.0) (10x Genomics) and a modified gene transfer format (GTF) file, where for each transcript, the feature type had been changed from transcript to exon. The starting GTF files came from refdata-cellranger-GRCh38-1.2.0.tar.gz or refdata-cellranger-GRCh38-3.0.0.tar.gz, and are available for download at https://support.10xgenomics.com/single-cell-gene-expression/software/downloads/3.0.

To down-sample sequencing reads or gene counts (UMIs) when comparing protocols, we used *downsampleReads* and *downsampleMatrix,* respectively from the `R` package(Lun et al., 2019) `DropletUtils` (v1.0.3 or higher). Reads were down-sampled to match the protocol with the lowest number of total reads. After down-sampling by total reads, we used *write10xCounts* from `DropletUtils` and a custom python script to generate an HDF5 file for input into our analysis pipelines described below.

### Quality control of scRNA-seq data

To maintain explicit control over all gene and cell quality control filters, in all our downstream analyses we used the raw feature-barcode matrix, rather than the filtered feature-barcode matrix generated by Cell Ranger. We removed low quality cells by requiring each cell to have a minimal number of UMIs and genes detected. We used different thresholds depending on the experimental modality (single cell or single nucleus) and on the 10x kit (V2 or V3 chemistry). For single nucleus data, we retained nuclei with at least 200 genes and 400 UMIs detected by V2 chemistry and with at least 500 genes and 1,000 UMIs detected by V3 chemistry. For single cell data, we retained cells with at least 500 genes and 1,000 UMIs detected by either V2 or V3 chemistry. For both data types, we filtered out those cells or nuclei where >20% of UMIs came from mitochondrial genes. Finally, we normalized the total UMIs per cell or nucleus to one-hundred thousand (CP100K), and log-transformed these values to report gene expression as E = log(CP100K+1).

We reported the following QC metrics: number of total reads per library sample, sequencing saturation (fraction of reads originating from an already-observed UMI as reported by Cell Ranger count), total recovered cells or nuclei, number of reads per cell or nucleus, number of UMIs per cell or nucleus, number of genes detected per cell or nucleus, fraction of UMIs in a cell or nucleus aligned to mitochondrial genes, fraction of droplets estimated to contain only ambient RNA (“empty drops”), fraction of cell or nucleus doublets, the number of detected cell types, and the pattern of copy number aberration (CNA) for malignant cells. For a subset of samples, we also calculated the UMI saturation for each cell or nucleus by subsampling from the total number of sequencing reads in the cell or nucleus(Wallrapp et al., 2017), the number of cells or nuclei per detected cell type, and the estimated level of ambient RNA in droplets containing cells.

We predicted droplets containing only ambient RNA and no cells using `EmptyDrops` (part of `DropletUtils`, v1.0.3 or higher), with the retain parameter set by the knee of the curve in the barcode rank plot (cell barcodes ranked by their total UMIs)(Lun et al., 2019). We predicted potential doublets using `Scrublet` (v0.2) with expected_doublet_rate = 0.06(Wolock et al., 2019). We estimated the levels of ambient RNA using `SoupX` (v0.3.1) and a set of cell-type specific marker genes(Young and Behjati, 2018) (**Supplementary Table 1**). Importantly, we flagged the doublets and empty drops and retained them in our analysis, instead of immediately filtering them out. Droplets that appear to contain doublets or empty drops can arise from many different effects, such as cellular differentiation or insufficient sequencing, and by carrying them through the analysis, potential doublets or empty drops can be more clearly interpreted in the context of the full dataset.

### Dimensionality reduction, clustering, and visualization

For each tumor sample, we analyzed the filtered expression matrix to identify cell subsets, as previously described(Shekhar et al., 2016; Wolf et al., 2018). We chose highly variable genes with a z-score cutoff of 0.5(Macosko et al., 2015). centered and scaled the expression of each gene to have a mean of zero and standard deviation of one, and performed dimensionality reduction on the variable genes using principal component analysis (PCA). We used the top 50 principal components (PCs) as input to Louvain graph-based clustering, with the resolution parameter set to 1.3. For each cluster of cells, we identified cluster-specific differentially expressed genes using the following tests: an AUC classifier, Welch’s *t*-test, and Fisher’s exact test. For tests that returned a p-value, we controlled the false discovery rate at 5% with the Benjamini-Hochberg procedure(Benjamini, 1995) We visualized gene expression and clustering results by embedding cells or nuclei profiles in a Uniform Manifold Approximation and Projection (UMAP)(Leland McInnes, 2018) of the top 50 PCs, with min_dist = 0.5, spread = 1.0, the number of neighbors = 15, and the Euclidean distance metric.

### Annotating cell subsets

For each cell subset identified by clustering, we assigned a cell type from the malignant, parenchymal, stromal, and immune compartments of the tumor microenvironment using a combination of differentially expressed genes, known gene signatures (**Supplementary Table 1**), and `SingleR` (v0.2.2)(Aran et al., 2019), an automated annotation package. When running `SingleR`, only cell types assigned to 30 or more cells were considered. When scoring cells for the expression of known gene signatures, we used the *AddModuleScore* function in `Seurat` (v2.3.4)(Butler et al., 2018). We note that overlapping expression programs between T cells and NK cells make these cell types sometimes more difficult to accurately identify.

We identified the malignant cells by inferring chromosomal copy number aberrations (CNAs) from the gene-expression data using inferCNV (v1.1.0)(Tickle T, 2019). On a sample-by-sample basis, we used the immune and endothelial cells as a healthy reference to estimate CNAs in the malignant cells. We created the count matrix file and annotation file for inferCNV by randomly subsetting the counts data to sample at most 2,000 cells or nuclei. We created a gene ordering file from the human GRCh38 assembly, which contains the chromosomal start and end positions for each gene. To run inferCNV, we used a cutoff of 0.1 for the minimum average read counts per gene among reference cells or nuclei, clustered according to the annotated cell types, denoised our output, ran an HMM to predict CNA level, implemented inferCNV’s i6 HMM model, and requested 8 threads for parallel steps.

### Comparing single cell and single nucleus RNA-Seq data

To compare profiles between single cell and single nucleus RNA-Seq data collected from the same sample, we used a batch-correction approach.

We performed batch correction using canonical correlation analysis (CCA) as implemented in `Seurat` (v2.3.4) (Butler et al., 2018). We selected 1,500 genes that were variable across both the cell and nucleus data, used those genes as input to *RunCCA* to compute the first 20 canonical components, and aligned the first 12 canonical components with *AlignSubspace*. The aligned canonical components represent a co-embedding of the cell and nucleus data, and we carried out clustering in this dimensionality-reduced space using *FindClusters*.

## Data availability

All main and supplementary figures have associated raw data. The counts matrices for each sample will be publicly available in GEO under data repository accession no. GSEXXX. Raw data will be available in the controlled access repository DUOS (https://duos.broadinstitute.org/#/home).

## Code availability

We implemented all major analysis steps, from FASTQ files to identifying cell subsets, in pipelines executed in a Cloud environment. We named this collection of pipelines scCloud, which may be executed in both a Cloud-based environment and a local, python environment(Butler et al., 2018).

Pipelines were written in the Workflow Description Language (WDL) and run on Cromwell in the Terra Cloud platform (https://app.terra.bio/), and data was stored in Google Cloud Plaform storage buckets. We wrote two WDL workflows: cellranger_workflow, a wrapper for running Cell Ranger mkfastq and count, and scCloud, a novel, fast, and scalable analysis pipeline for single cell and single nucleus RNA-Seq data. All analysis workflows will be publicly available through https://github.com/klarman-cell-observatory/scCloud.

We ran additional quality control steps, cell-subset annotations, and protocol comparison steps in R (v3.5) by converting the single-cell AnnData objects from scCloud into Seurat objects. An example script for this analysis will be made publicly available at https://github.com/klarman-cell-observatory/HTAPP-Pipelines.

## References

Al-Hajj, M., Wicha, M. S., Benito-Hernandez, A., Morrison, S. J., and Clarke, M. F. (2003). Prospective identification of tumorigenic breast cancer cells. Proc Natl Acad Sci U S A 100, 3983–3988.

Aran, D., Looney, A. P., Liu, L., Wu, E., Fong, V., Hsu, A., Chak, S., Naikawadi, R. P., Wolters, P. J., Abate, A. R., et al. (2019). Reference-based analysis of lung single-cell sequencing reveals a transitional profibrotic macrophage. Nat Immunol 20, 163–172.

Bakken, T. E., Hodge, R. D., Miller, J. A., Yao, Z., Nguyen, T. N., Aevermann, B., Barkan, E., Bertagnolli, D., Casper, T., Dee, N., et al. (2018). Single-nucleus and single-cell transcriptomes compared in matched cortical cell types. PLoS One 13, e0209648.

Benjamini, Y., and Yosef Hochberg (1995). “Controlling the false discovery rate: a practical and powerful approach to multiple testing”. Journal of the Royal statistical society: series B (Methodological) 571 289–300.

Butler, A., Hoffman, P., Smibert, P., Papalexi, E., and Satija, R. (2018). Integrating single-cell transcriptomic data across different conditions, technologies, and species. Nature biotechnology 36, 411–420.

Cieslik, M., and Chinnaiyan, A. M. (2018). Cancer transcriptome profiling at the juncture of clinical translation. Nat Rev Genet 19, 93–109.

Drokhlyansky, E., Smillie, C. S., Van Wittenberghe, N., Ericsson, M., Griffin, G. K., Dionne, D., Cuoco, M. S., Goder-Reiser, M. N., Sharova, T., and Aguirre, A. J. (2019). The enteric nervous system of the human and mouse colon at a single-cell resolution. bioRxiv, 746743.

Filbin, M. G., Tirosh, I., Hovestadt, V., Shaw, M. L., Escalante, L. E., Mathewson, N. D., Neftel, C., Frank, N., Pelton, K., Hebert, C. M., et al. (2018). Developmental and oncogenic programs in H3K27M gliomas dissected by single-cell RNA-seq. Science 360, 331–335.

Gao, R., Kim, C., Sei, E., Foukakis, T., Crosetto, N., Chan, L. K., Srinivasan, M., Zhang, H., Meric-Bernstam, F., and Navin, N. (2017). Nanogrid single-nucleus RNA sequencing reveals phenotypic diversity in breast cancer. Nat Commun 8, 228.

Gaublomme, J. T., Li, B., McCabe, C., Knecht, A., Yang, Y., Drokhlyansky, E., Van Wittenberghe, N., Waldman, J., Dionne, D., Nguyen, L., et al. (2019). Nuclei multiplexing with barcoded antibodies for single-nucleus genomics. Nat Commun 10, 2907.

Guillaumet-Adkins, A., Rodriguez-Esteban, G., Mereu, E., Mendez-Lago, M., Jaitin, D. A., Villanueva, A., Vidal, A., Martinez-Marti, A., Felip, E., Vivancos, A., et al. (2017). Single-cell transcriptome conservation in cryopreserved cells and tissues. Genome Biol 18, 45.

Habib, N., Avraham-Davidi, I., Basu, A., Burks, T., Shekhar, K., Hofree, M., Choudhury, S. R., Aguet, F., Gelfand, E., Ardlie, K., et al. (2017). Massively parallel single-nucleus RNA-seq with DroNc-seq. Nat Methods.

Habib, N., Li, Y., Heidenreich, M., Swiech, L., Avraham-Davidi, I., Trombetta, J. J., Hession, C., Zhang, F., and Regev, A. (2016). Div-Seq: Single-nucleus RNA-Seq reveals dynamics of rare adult newborn neurons. Science 353, 925–928.

Hermansen, J. U., Tjonnfjord, G. E., Munthe, L. A., Tasken, K., and Skanland, S. S. (2018). Cryopreservation of primary B cells minimally influences their signaling responses. Sci Rep 8, 17651.

Jerby-Arnon, L., Shah, P., Cuoco, M. S., Rodman, C., Su, M. J., Melms, J. C., Leeson, R., Kanodia, A., Mei, S., Lin, J. R., et al. (2018). A Cancer Cell Program Promotes T Cell Exclusion and Resistance to Checkpoint Blockade. Cell 175, 984–997.e924.

Leland McInnes, J. H., James Melville (2018). UMAP: Uniform Manifold Approximation and Projection for Dimension Reduction. In, (eprint arXiv:1802.03426).

Li, B., Gould, J., Rosen, Y., Rozenblatt-Rosen, O., and Regev, A. (2019). https://github.com/klarman-cell-observatory/KCO. In.

Lun, A. T. L., Riesenfeld, S., Andrews, T., Dao, T. P., Gomes, T., participants in the 1st Human Cell Atlas, J., and Marioni, J. C. (2019). EmptyDrops: distinguishing cells from empty droplets in droplet-based single-cell RNA sequencing data. Genome Biol 20, 63.

Macosko, E. Z., Basu, A., Satija, R., Nemesh, J., Shekhar, K., Goldman, M., Tirosh, I., Bialas, A. R., Kamitaki, N., Martersteck, E. M., et al. (2015). Highly Parallel Genome-wide Expression Profiling of Individual Cells Using Nanoliter Droplets. Cell 161, 1202–1214.

McDivitt, R. W., Stone, K. R., and Meyer, J. S. (1984). A method for dissociation of viable human breast cancer cells that produces flow cytometric kinetic information similar to that obtained by thymidine labeling. Cancer Res 44, 2628–2633.

Neftel, C., Laffy, J., Filbin, M. G., Hara, T., Shore, M. E., Rahme, G. J., Richman, A. R., Silverbush, D., Shaw, M. L., Hebert, C. M., et al. (2019). An Integrative Model of Cellular States, Plasticity, and Genetics for Glioblastoma. Cell.

Puram, S. V., Tirosh, I., Parikh, A. S., Patel, A. P., Yizhak, K., Gillespie, S., Rodman, C., Luo, C. L., Mroz, E. A., Emerick, K. S., et al. (2017). Single-Cell Transcriptomic Analysis of Primary and Metastatic Tumor Ecosystems in Head and Neck Cancer. Cell 171, 1611–1624 e1624.

Shekhar, K., Lapan, S. W., Whitney, I. E., Tran, N. M., Macosko, E. Z., Kowalczyk, M., Adiconis, X., Levin, J. Z., Nemesh, J., Goldman, M., et al. (2016). Comprehensive Classification of Retinal Bipolar Neurons by Single-Cell Transcriptomics. Cell 166, 1308–1323 e1330.

Stewart, E., Federico, S. M., Chen, X., Shelat, A. A., Bradley, C., Gordon, B., Karlstrom, A., Twarog, N. R., Clay, M. R., Bahrami, A., et al. (2017). Orthotopic patient-derived xenografts of paediatric solid tumours. Nature 549, 96–100.

Stewart, E., Shelat, A., Bradley, C., Chen, X., Federico, S., Thiagarajan, S., Shirinifard, A., Bahrami, A., Pappo, A., Qu, C., et al. (2015). Development and characterization of a human orthotopic neuroblastoma xenograft. Dev Biol 407, 344–355.

Tickle T, T. I., Georgescu C, Brown M, Haas B (2019). inferCNV of the Trinity CTAT Project, https://github.com/broadinstitute/inferCNV. In.

Tirosh, I., Venteicher, A. S., Hebert, C., Escalante, L. E., Patel, A. P., Yizhak, K., Fisher, J. M., Rodman, C., Mount, C., Filbin, M. G., et al. (2016). Single-cell RNA-seq supports a developmental hierarchy in human oligodendroglioma. Nature 539, 309–313.

Venteicher, A. S., Tirosh, I., Hebert, C., Yizhak, K., Neftel, C., Filbin, M. G., Hovestadt, V., Escalante, L. E., Shaw, M. L., Rodman, C., et al. (2017). Decoupling genetics, lineages, and microenvironment in IDH-mutant gliomas by single-cell RNA-seq. Science 355.

Wallrapp, A., Riesenfeld, S. J., Burkett, P. R., Abdulnour, R. E., Nyman, J., Dionne, D., Hofree, M., Cuoco, M. S., Rodman, C., Farouq, D., et al. (2017). The neuropeptide NMU amplifies ILC2-driven allergic lung inflammation. Nature 549, 351–356.

Wolf, F. A., Angerer, P., and Theis, F. J. (2018). SCANPY: large-scale single-cell gene expression data analysis. Genome Biol 19, 15.

Wolock, S. L., Lopez, R., and Klein, A. M. (2019). Scrublet: Computational Identification of Cell Doublets in Single-Cell Transcriptomic Data. Cell Syst 8, 281–291 e289.

Young, M. D., and Behjati, S. (2018). SoupX removes ambient RNA contamination from droplet based single cell RNA sequencing data. 303727.

